# Representational similarity and pattern classification of fifteen emotional states induced by movie clips and text scenarios

**DOI:** 10.1101/2025.10.17.682958

**Authors:** Yaohui Ding, Nathan M. Muncy, John L. Graner, Joel S. White, Alisa C. Schutz, Leonard Faul, John M. Pearson, Kevin S. LaBar

## Abstract

How do different emotional states relate to each other and how are they represented in the human brain? These are important questions in the field of affective neuroscience, with profound scientific and vast clinical implications. Different theories of emotion tend to emphasize the relative importance of distinct psychological constructs, for example categorical labels (e.g., fear, joy, or sadness) versus dimensional ratings (e.g., valence and arousal), for understanding human emotions. To investigate whether categorical or dimensional constructs correspond better to patterns of brain activity associated with human affective experiences, we experimentally induced 15 emotions (spanning positive, negative, and neutral valence) in 136 participants using 150 short movie clips and 150 one- or two-sentence text scenarios, while their blood oxygenation-level dependent activity was recorded in a magnetic resonance imaging scanner. Results from our representational similarity analyses suggest participants’ brain activity significantly correlated with their categorical labeling of, but not their dimensional rating of, the movie clips and text scenarios. Subsequently, we were also able to decode the categorical labels of these emotional stimuli using whole-brain multi-voxel pattern classification, with important voxels found in many cortical, limbic, subcortical, cerebellar, and brainstem regions. Finally, we found similar clusters of emotions through exploratory hierarchical clustering analyses of participants’ categorical labeling of and brain responses to these stimuli. Taken together, these findings greatly advance our understanding of how a large set of human emotions are related to each other both in terms of the participants’ self-report and their brain activity.

**Significance Statement:** We successfully decoded fifteen emotional states induced by movie clips and text scenarios based on participants’ brain responses (blood oxygenation-level dependent signals) to these stimuli using multi-voxel pattern classification (a supervised machine learning approach). We also found that participants’ brain responses correlated with their self-reported emotional experience, i.e. which emotion they felt while they were presented with these stimuli, using representative similarity analysis (an unsupervised approach). Finally, we found that these emotions are organized in very similar ways both in terms of participants’ categorical labeling and their brain responses to these stimuli. Together, these data-driven and computational modeling-based findings greatly advance our understanding of how a large set of emotions are organized and represented in the human brain.

## Introduction

How do different emotional states relate to each other and how are they represented in the human brain? This is one of the central questions in the field of affective neuroscience, with profound scientific and vast clinical implications. Different theoretical orientations make different predictions about the facets of emotional experience that are important for distinguishing emotions from one another. Discrete emotion theories predict that emotions tend to be experienced in distinct and universal categories with unique evolutionary origins and brain systems, at least for a few basic emotions (1,2). Dimensional theories instead postulate that specific emotions emerge from a small set of intersecting affective dimensions such as arousal and valence (3). Appraisal theories include other dimensions of experience that are important for cognitively evaluating the meaning of an emotional situation as it dynamically unfolds, some of which are more universal, like curiosity, and some of which are more culturally specific, like moral appraisals that contribute to normative significance evaluation (4). Constructionist theories postulate that each instance of emotion is constructed uniquely in the service of allostatic goals based on contextual demands, internal models of the body in the world, the learning history of the individual, and sociocultural factors (5,6). Finally, according to the semantic space theory (7,8), emotions can be understood in a semantic space that is characterized by three major aspects of our emotional experiences: dimensionality, distribution, and conceptualization. The dimensionality aspect is concerned with determining the number of meaningful experiences or expressions that are distinct (7) within the semantic space; the distribution aspect captures the clustering or blending of emotional states; and the conceptualization aspect is concerned with how emotions are perceived by those experiencing them.

The variety of theories of how emotion arise are also reflected in the understanding of emotional representation in the brain. There is no current consensus other than that there is no one-to-one correspondence between a given emotion and univariate activity in a given brain region (9). With the rise and maturation of multivariate pattern classification (MVPA) techniques (10), researchers have started to investigate whether emotional states could be decoded or classified from multi-voxel patterns of blood oxygenation level dependent (BOLD) activity. For example, Kassam and colleagues (11) decoded nine emotional states based on the neural responses from the same participant, other participants, and a different emotion induction modality. Kragel and LaBar (12) decoded seven emotional states induced from movie and music and showed that the neural biomarkers of these emotional states are categorically distinct and widely distributed across the entire brain. In this work (12) they also found that classification errors, the extent to which one emotion was confused with another emotion, were best explained by the pairwise distances of the emotions in a 7-dimensional space spanned by the seven categories. Similarly, Saarimäki and colleagues (13) found specific neural signatures of six basic emotions (disgust, fear, happiness, sadness, anger, and surprise) induced by two stimulus modalities: short movies and mental imagery. Further, Saarimäki et al. showed there was a robust correspondence between the behavioral and neural similarities of fourteen different emotional states (14). They also found that the neural bases of these emotions included brain regions widely distributed across cortical and subcortical areas. These multivariate studies overcame some limitations of earlier studies: the static nature of the emotional stimuli, a small set of emotions, and the traditional univariate approach to brain mapping. Accordingly, they have begun to answer a wider array of questions regarding how emotions are represented in the brain. However, the results from these studies have not converged on a clear answer, and they were still limited by relatively small sample sizes (11-13), minimal involvement of subcortical regions (12), under-sampling of positive emotions (12), and a single theoretical perspective (14).

Cowen and Keltner (7) showed that semantic space theory was able to capture more complex and nuanced aspects of human subjective experiences to various types of emotionally evocative stimuli, e.g., ancient art, face and voice, short video clips, music, and prosody. These studies emphasized that discrete emotions were represented with fuzzy categorical boundaries foundational to the semantic spaces, while dimensions of emotional experience like valence and arousal provided means to bridge these fuzzy categories. In addition, different induction modalities elicited semantic spaces that differed in dimensionality and structure. These studies supporting semantic space theory have primarily focused on the subjective experience of emotions based on self-report (7,15). In an extension to neural data, Horikawa and colleagues (16) found that the brain representations of 27 emotions were categorical in nature and involved distributed, transmodal brain regions. Though this study investigated a large number of emotions, its generalizability was limited by the very small sample size (N=5), single induction method, and a within-subject approach to representational similarity and decoding analyses.

### Research Questions and Hypotheses

In the current study, we experimentally induced fifteen emotional states using two modalities, short movie clips and text scenarios, while participants underwent whole-brain functional magnetic resonance imaging (fMRI). We expanded upon previous work by including a large number of emotions (N = 15) that spanned negative, neutral, and positive valence as well as low-to-high arousal levels, multiple induction methods (movies and scenarios), a large number of stimuli (N = 150 for each induction modality), and a large sample size (N=136). To investigate the neural underpinnings of human emotional experiences and how different emotional states relate to each other, we utilized three complementary computational modeling techniques, namely representational similarity analysis, hierarchical clustering analysis, and partial least squares discriminant analysis (PLS-DA, a type of supervised machine learning technique). We sought to answer the following questions: (1) Do participants’ subjective experience (e.g., self-reported categorical endorsement and valence/arousal ratings) of the fifteen emotional states correlate with their neural response to the emotional stimuli? (2) Relatedly, how are these fifteen emotional states organized in terms of participants’ self-report and brain response, i.e., which emotions are more similar or dissimilar to each other? (3) Can we use participants’ brain responses to classify the normative emotion labels of movie and scenario emotional stimulus blocks experienced in the scanner? (4) Finally, if the classification is successful, what properties of these emotional stimuli (e.g. categorical endorsement or valence/arousal ratings) better explain the classification errors? We hypothesized we would achieve above-chance supervised classification across emotions for both modalities, but that the representations would differ somewhat across induction methods due to their different natures (e.g., reading and imagining hypothetical scenarios vs. viewing dynamic movie clips). Given that some prior multivariate studies have favored the primacy of categorical over dimensional representations (7,12), we hypothesized that classification errors would be better explained by categorical rather than dimensional ratings (3). Consistent with some constructionist theories (17) and the prior whole-brain fMRI decoding studies reviewed above, we hypothesized that emotions would broadly span cortical, limbic, and subcortical structures.

## Results

### Successful Emotion Induction Based on Participant Self-Report

Our task successfully induced all fifteen emotions in our participants using both movie and scenario stimuli, based on participants’ self-report. In total, we gathered participants’ emotion endorsement from 3556 blocks of movie and 3788 blocks of scenario stimuli. Performance metrics (SI Appendix, Table S1) computed from participants’ emotion endorsement of the movie blocks were as follows: balanced accuracy (0.7395, CI = [0.7255, 0.7534]; Fig. 1-B1), macro-averaged sensitivity (0.7395, CI = [0.7255, 0.7534]), macro-averaged specificity (0.9814, CI = [0.9803, 0.9824]), macro-averaged precision (0.7488, CI = [0.7347, 0.7627]), and macro-averaged F1-score (0.7441, CI = [0.7303, 0.7580]); all p < 0.001. Emotion-specific performance metrics of participants’ endorsement of movie blocks are tabulated in SI Appendix, Table S2. Additionally, SI Appendix, Fig. S1-A1 shows the endorsed and expected frequencies of each emotion and SI Appendix, Fig. S1-B1 plots the averaged intensity rating of each emotion from the movie blocks. Similarly, performance metrics (SI Appendix, Table S3) computed from participants’ subjective emotion endorsement of the scenario blocks were as follows: balanced accuracy (0.7031, CI = [0.6882, 0.7174]; Fig. 1-B2), macro-averaged sensitivity (0.7031, CI = [0.6882, 0.7174]), macro-averaged specificity (0.9788, CI = [0.9777, 0.9798]), macro-averaged precision (0.7238, CI = [0.7100, 0.7371]), and macro-averaged F1-score (0.7133, CI = [0.6992, 0.7268]); all p < 0.0001. Emotion-specific performance metrics of participants’ endorsement of scenario blocks are tabulated in SI Appendix, Table S4. Finally, SI Appendix, Fig. S1-A2 shows the endorsed and expected frequencies of each emotion and SI Appendix, Fig. S1-B2 plots the averaged intensity rating of each emotion from the scenario blocks.

**Fig. 1.**
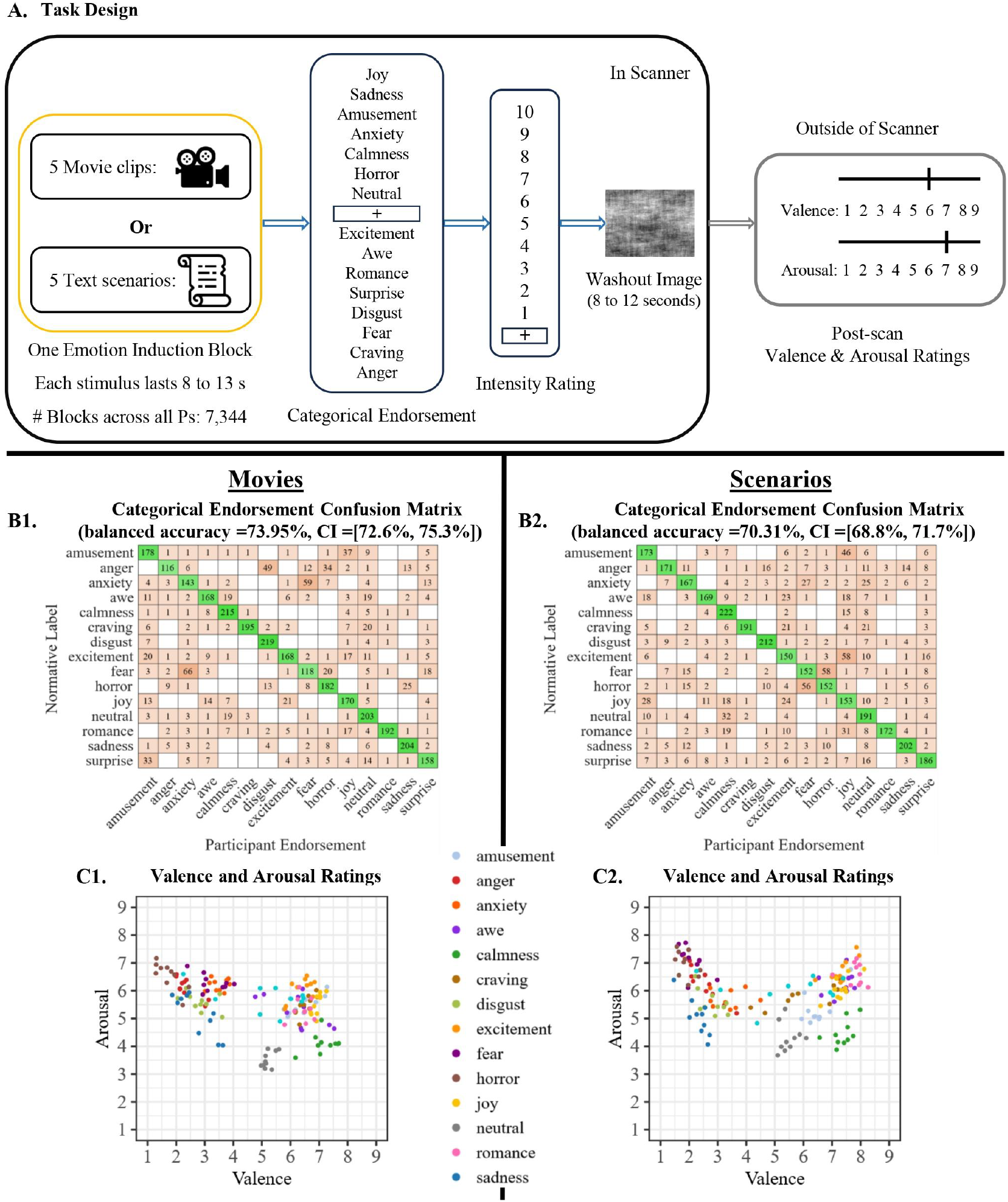
Task Design and Participant In-Scanner Emotion Endorsement and Post-Scan Valence-Arousal Ratings. *A:* An illustration of the task design. Movie clips and scenarios were presented in two separate MRI sessions, and a block of 5 stimuli all came from the same emotion category. Within each MRI session (movies or scenarios), each participant encountered two blocks of each of the 15 emotion categories (order randomized). Each emotion block is followed by emotion endorsement (forced choice), intensity rating, and an 8- to 12-second washout period. After the participant comes out of the scanner, they are asked to provide valence and arousal ratings of half the stimuli they encountered in the scanner, using a 9-point Likert scale. *B1, B2:* Confusion matrices of participant emotion endorsement of movie (balanced accuracy = 73.95%, CI = [72.6%, 75.35%]) and scenario (balanced accuracy = 70.31%, CI = [68.8%, 71.7%]) blocks, respectively. *C1, C2:* Scatter plots of post-scan valence and arousal ratings. Each dot represents the averaged (across all participants) valence and arousal rating for a stimulus. There are 150 movie stimuli and 150 scenario stimuli in total.

### Representational Similarity between Participant Self-Report and Brain Responses

There is a strong and positive correlation between the pairwise categorical distances and pairwise dimensional distances for both the movie (Spearman rho = 0.5932, CI = [0.4563, 0.7001], p = 0.0004; Fig. 2-B2) and scenario (Spearman rho = 0.7509, CI = [0.6713, 0.8057], p = 0.0002; Fig. 2-D2) stimuli (see “Materials and Methods: Representation Similarity Analysis” section for definitions of categorical and dimensional distances). These results suggest that emotions which are more distant from each other in the valence-arousal space tend to be categorized in different ways. However, the relationship between participants’ self-report and brain responses to these stimuli is more nuanced. Specifically, there is a moderate positive correlation between pairwise categorical distances and pairwise neural distance for the movie stimuli (Spearman rho = 0.4113, CI = [0.2132, 0.5902], p = 0.0004; Fig. 2-B1) but not for the scenario stimuli (Spearman rho = 0.1673, CI = [-0.02761, 0.3518], p = 0.1208; Fig. 2-d=D1). In terms of the relationship between pairwise neural distance and pairwise dimensional distance, we did not find a statistically significant correlation for movie (Spearman rho =0.0846, CI = [-0.1193, 0.2859], p = 0.3004; Fig. 2-B3) or scenario (Spearman rho = 0.1465, CI = [-0.06299, 0.344], p = 0.0562; Fig. 2-D3) stimuli. Overall, these results suggest participants’ categorical and dimensional ratings of the emotional stimuli are highly consistent with each other, whereas the correspondence between participants’ self-report and brain responses to the stimuli is stronger for the categorical ratings, at least for movie inductions.

**Fig. 2.**
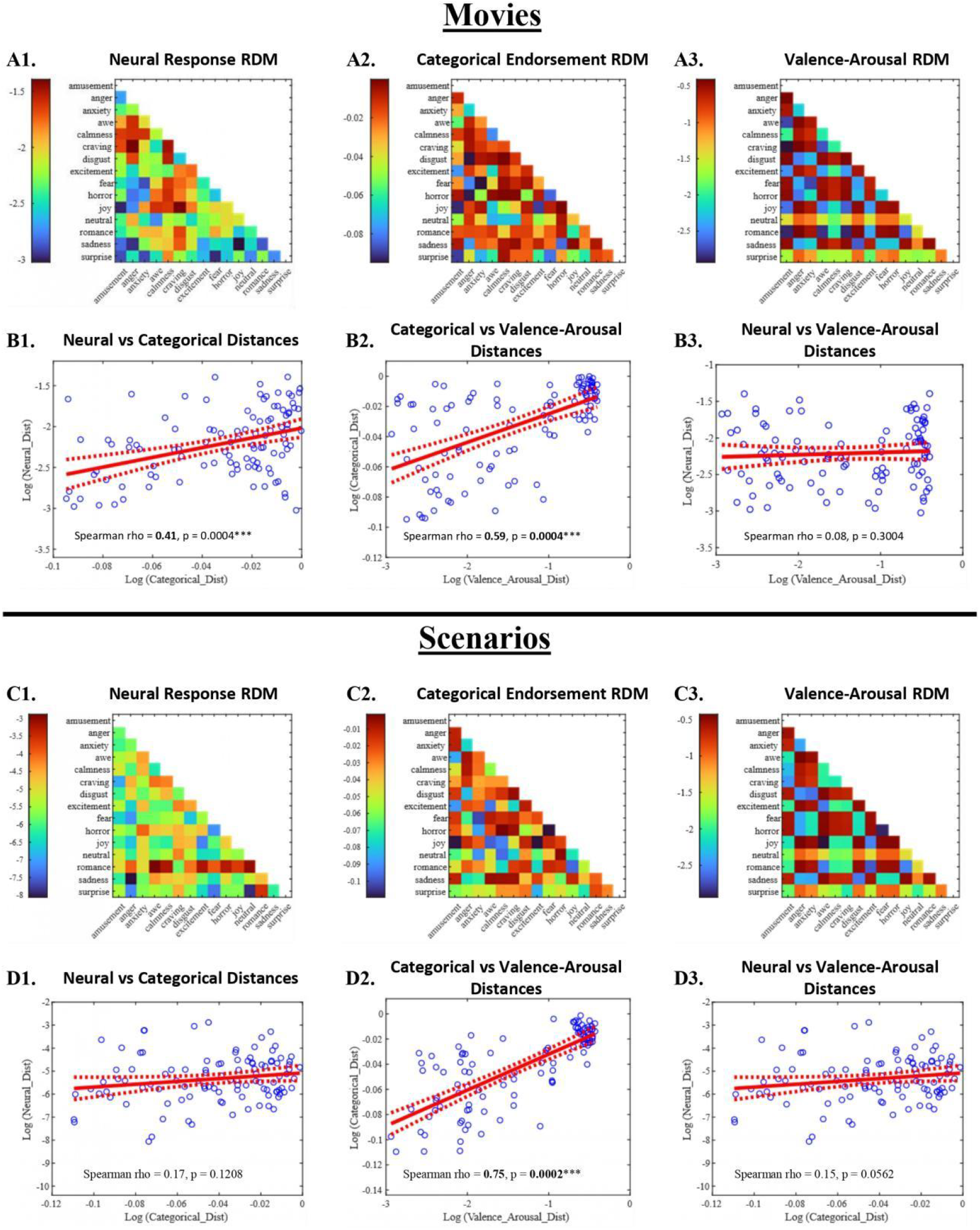
Representational Similarity between Participant Self-Report and Brain Responses to Movie and Scenario Blocks. *A1, A2, A3:* Representational dissimilarity matrices (RDM) of patterns of BOLD activity, participant categorical endorsement of the movie blocks, and participant valence/arousal ratings of the movie stimuli. *B1, B2, B3:* Spearman correlation between pairwise neural distance and pairwise categorical distance (rho = 0.4113, permutation p = 0.0004), Spearman correlation between pairwise categorical distance and pairwise valence/arousal distance (rho = 0.5932, p = 0.0004), and Spearman correlation between pairwise neural distance and pairwise valence/arousal distance (rho = 0.0846, p = 0.3004) for movie blocks. *C1, C2, C3:* Representational dissimilarity matrices (RDM) of patterns of BOLD activity, participant categorical endorsement of the scenario blocks, and participant valence/arousal ratings of scenario stimuli. *D1, D2, D3:* Spearman correlation between pairwise neural distance and pairwise categorical distance (rho = 0.1673, p = 0.1208), Spearman correlation between pairwise categorical distance and pairwise valence/arousal distance (rho = 0.7509, p = 0.0002), and Spearman correlation between pairwise neural distance and pairwise valence/arousal distance (rho = 0.1465, p = 0.0562) for scenario blocks. Number of permutations =10000.

### Successful Decoding of Emotional States from Movie and Scenario Blocks

After anomaly detection and outlier removal, 3143 blocks of whole-brain, voxel-wise beta weights were retained from the movie stimuli and 3247 blocks from the scenario stimuli. The PLS-DA classifiers trained on beta weights from movie blocks achieved a strong performance for decoding the emotional states from these stimuli: balanced accuracy (0.5371, CI = [0.5208, 0.5533]; Fig. 3-A1), macro-averaged sensitivity (0.5371, CI = [0.5208, 0.5533]), macro-averaged specificity (0.967, CI = [0.9657, 0.9682]), macro-averaged precision (0.5274, CI = [0.5076, 0.5469]), and macro-averaged F1-score (0.5322, CI = [0.5146, 0.5493]); all p < 0.001 (see SI Appendix, Table S7 for more information). Emotion-specific performance metrics of PLS-DA classification of movie blocks are tabulated in SI Appendix, Table S8. The macro-averaged area under the curve (AUC) value of the PLS-DA movie classifiers is 0.91 (CI = [0.89, 0.94]; Fig. 3-C1). In contrast, the PLS-DA classifiers trained on beta weights from the scenario blocks performed more poorly, though still better than chance (SI Appendix, Fig. S2-B2): balanced accuracy (0.1862, CI = [0.1733, 0.199]; Fig. 3-A2), macro-averaged sensitivity (0.1862, CI = [0.1733, 0.199]), macro-averaged specificity (0.9418, CI = [0.9409, 0.9427]), macro-averaged precision (0.1812, CI = [0.1663, 0.1963]), and macro-averaged F1-score (0.1837, CI = [0.1701, 0.1973]); all p < 0.001 (see SI Appendix, Table S9 for more information). Emotion-specific performance metrics of PLS-DA classification of scenario blocks are tabulated in SI Appendix, Table S10. Macro-averaged AUC value of the PLS-DA scenario classifiers is 0.66 (CI = [0.89, 0.94]; Fig. 3-C2). When comparing the four types of Poisson regression models described in “Materials and Methods: Post-hoc classification error analysis and Bayesian model comparison”, we found that the Poisson regression model with the categorical rating regressor explained the errors the best for both movies (log Bayes factor = 21.89; Fig. 3-D1) and scenarios (log Bayes factor = 20.5241; Fig. 3-D2). The beta coefficients from the categorical Poisson models are the following: intercept (3.7192, CI = [3.4976, 3.9408], t = 16.781, p<0.0001), categorical pairwise distance regressor (-1.6995, CI = [-1.9108, -1.4882], t = - 8.0411, p < 0.0001) for movies, see Fig. 3-E1. Similarly, we have the following estimates for scenarios: intercept (4.0192, CI = [3.8180, 4.2204], t = 19.977, p < 0.0001), categorical pairwise distance regressor (-1.4124, CI = [-1.6136, -1.2112], t = -7.3913, p < 0.0001), see Fig. 3-E2. The fact that the estimates of the categorical pairwise distance regressor are negative tells us an increase in categorical pairwise distance between two emotions leads to lower confusion between those two emotions, as predicted by categorical theories of emotion. The correlations between PLS-DA classification errors and pairwise categorical endorsement distance for the movie and scenario blocks are shown in Fig. 3-F1 and Fig. 3-F2, respectively. There are negative correlations between PLS-DA errors and pairwise categorical distance for both movie and scenario stimuli. Similarly, the correlations between PLS-DA classification errors and pairwise valence/arousal distance for the movie and scenario blocks are shown in Fig. 3-G1 and Fig. 3-G2. Here there is a lack of correlation between PLS-DA errors and pairwise valence/arousal distance, which is not predicted by dimensional theories of emotion.

**Fig. 3.**
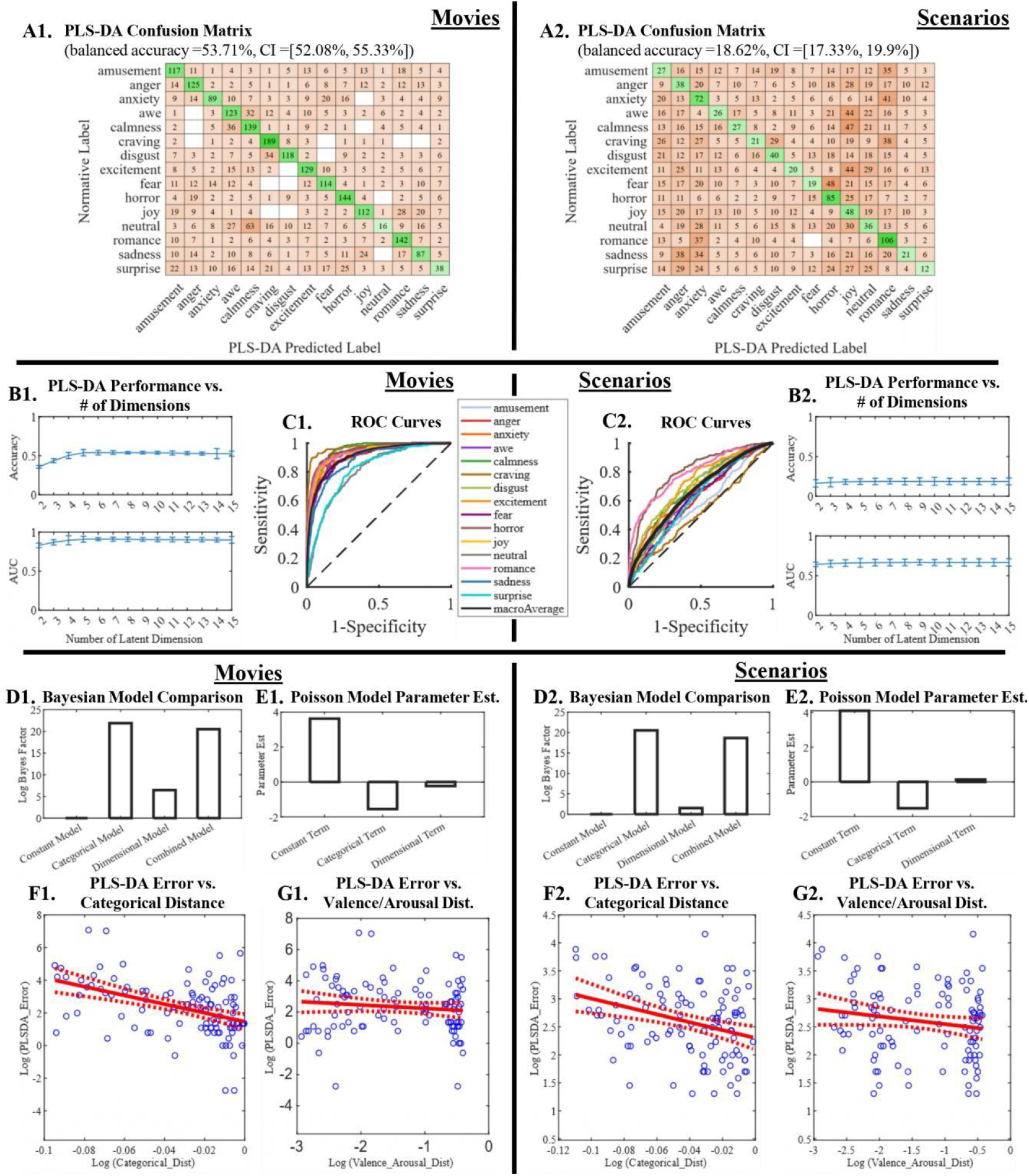
PLS-DA Classification Results. *A1, A2:* Confusion matrices from PLS-DA classification of movie (balanced accuracy = 0.5371, CI = [0.5208, 0.5533], p = 0.0002) and scenario (balanced accuracy = 0.1862, CI = [0.1733, 0.199], p = 0.0002) blocks. *B1, B2:* PLS-DA classification accuracy versus number of latent dimensions used. *C1, C2:* Receiver operating characteristic (ROC) curves of PLS-DA classification of movie (macro-averaged AUC = 0.91, CI = [0.89, 0.94]) and scenario (macro-averaged AUC = 0.66, CI = [0.62, 0.70]) blocks. *D1, D2:* Bar plots of the log Bayes factors from four Poisson regression models predicting PLS-DA classification errors from movie and scenario blocks. *E1, E2:* Bar plots of the parameter estimate from the combined Poisson regression model for predicting PLS-DA classification errors from movie and scenario blocks. *F1, F2:* Spearman correlation between PLS-DA classification errors and pairwise categorical distance from movie (rho = -0.4308, p = 0.0004) and scenario (rho = - 0.2926, p = 0.0048) blocks. *G1, G2:* Spearman correlation between PLS-DA classification errors and pairwise valence/arousal distance from movie (rho = -0.108, p = 0.2412) and scenario (rho = -0.1432, p = 0.104) blocks. Dist = distance; Est. = Estimates.

### Similar Emotion Clusters based on Participant Categorical Endorsement and Neural Responses to Emotion Induction

Several interesting and meaningful clusters of emotions emerged from hierarchical clustering of participant’s categorical endorsement of the movie (Fig. 4-A1) and scenario (Fig. 4-A2) inductions. For example, for both movies and scenarios, we see a disgust-anger cluster for negative emotions and an amusement-joy-excitement cluster for positive emotions. In contrast, the emotion clusters from participants’ valence/arousal ratings of the movie (Fig. 4-B1) and scenario stimuli (Fig. 4-B2) are more dichotomous. Additionally, we observe that the clusters from the movie and scenario subjective endorsement data are similar. However, this correspondence between movie and scenario stimuli does not hold for the clusters computed from participants’ brain responses to the emotional stimuli, for which the clusters from the movie stimuli (Fig. 4-C1) are quite different from those from the scenario stimuli (Fig. 4-C2). Finally, the emotion clusters computed from participants’ categorical endorsement (Fig. 4-A1) and the clusters from participants’ neural responses (Fig. 3-C1) are very similar for the movie stimuli. This similarity is supported by the result (Fig. 2-B1) from the RSA analysis.

**Fig. 4.**
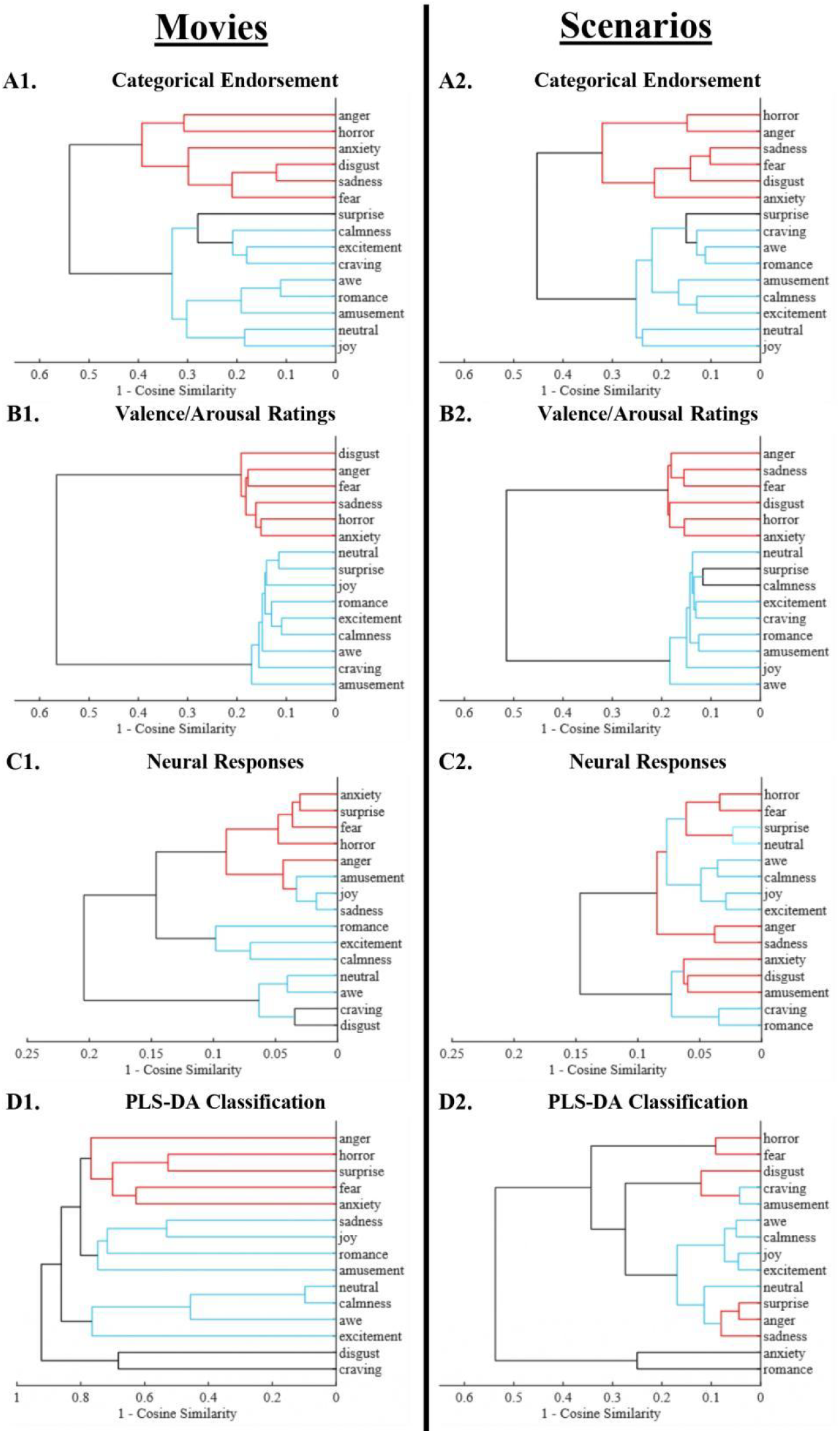
Dendrograms Showing the Emotion Clusters from Participants’ Categorical Endorsement, Valence/Arousal Ratings, and Brain Responses to Movie and Scenario Blocks. *A1, A2:* Hierarchical clustering of the categorical representation dissimilarity matrices (RDMs) from movie and scenario blocks. *B1, B2:* Hierarchical clustering of the Valence/Arousal RDMs from movie and scenario blocks. *C1, C2:* Hierarchical clustering of the neural RDMs from the movie and scenario blocks. D1, D2: Hierarchical clustering of the confusion matrices of the PLS-DA classifier. Note: Red represents, generally, negatively valenced emotions whereas teal represents positively valenced emotions.

### Maps of PLS-DA coefficients

When inspecting the voxel maps of the informative PLS-DA coefficients (Fig. 5 and 6), the first observation we can make is that brain regions important for decoding each emotion are widely distributed across cortical, limbic, and subcortical regions. Notably, the PLS-DA classifiers identified several subcortical regions, including in the brainstem and the cerebellum, which were missing in prior studies. Our preliminary analysis also showed that the classification accuracy from whole-brain patterns of activity exceeds the classification accuracies obtained with activity from each of the seven canonical resting-state networks separately (see SI Appendix, Fig. S3-A1). This distributed pattern of PLS-DA coefficients suggests that our emotional experience is an integrative process, where neural information from the sensory, limbic, subcortical, and cortical association regions all contribute to the classifier’s ability to differentiate one emotion from another, as predicted by constructionist theories. Additionally, we see that visual and other sensory areas are more involved in decoding the movie stimuli, suggesting that the emotion representations could be more perceptually-driven. In contrast, the default mode and other associative regions play more important roles in decoding the hypothetical scenario stimuli, which possibly indicates more abstract and internally-generated emotion representations. These differences suggest that neural representations of emotion, especially during their initial processing, could be partly modality-dependent, as predicted by semantic space theory. Taken together, the results from our study support the idea that neural representation of emotional states is distributed, high-dimensional, and categorical, and that the regions (or networks) engaged may depend in part on the induction modality and the perceptual richness of the stimuli.

**Fig. 5.**
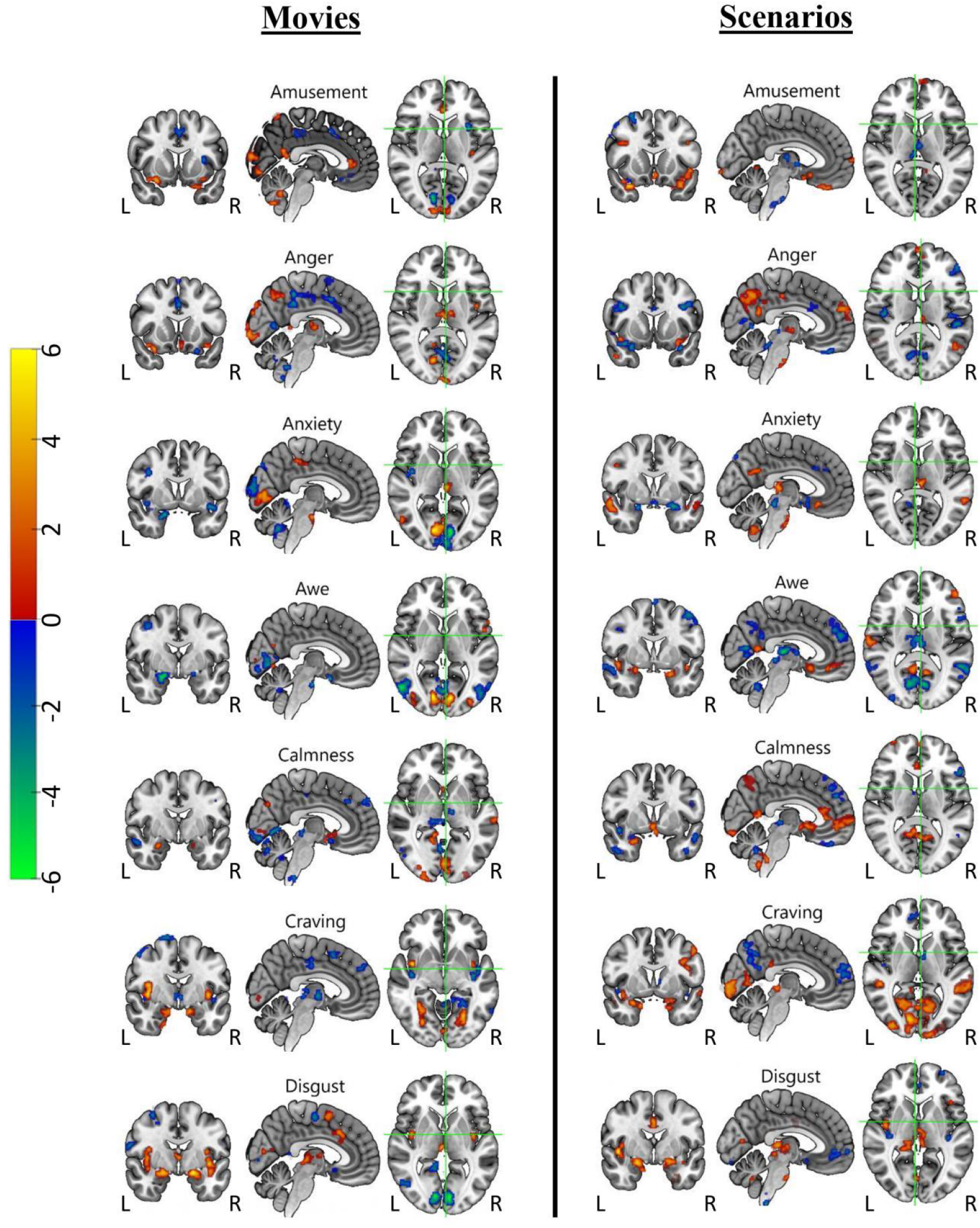
Maps of Informative Coefficients from PLS-DA Classification of Movie (left) and Scenario (right) Inductions (see SI Appendix for selection and clustering details).

**Fig. 6.**
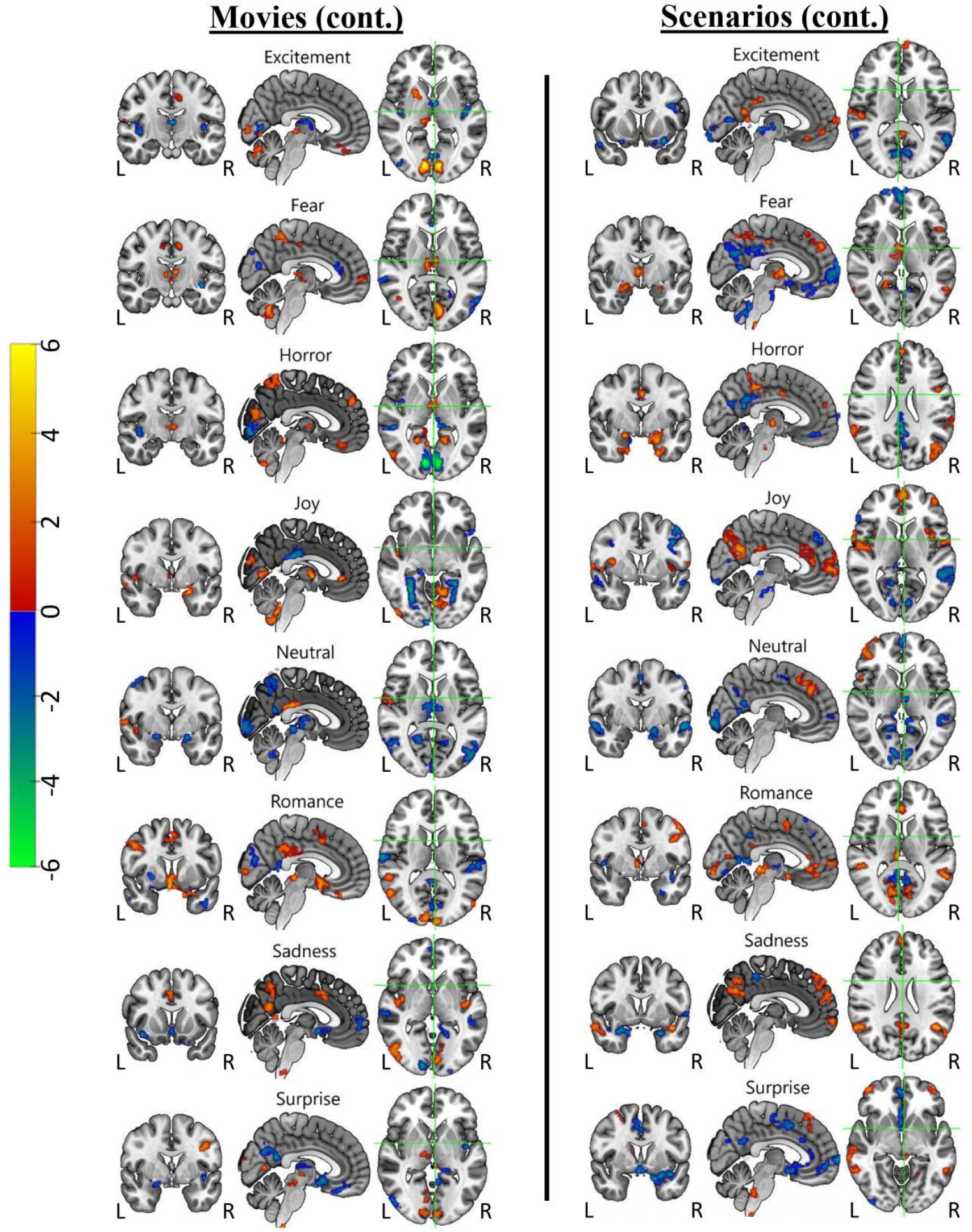
Maps of Informative Coefficients from PLS-DA Classification of Movie (left) and Scenario (right) Inductions (continued from Fig. 5; see SI Appendix for selection and clustering details).

## Discussion

### There is a moderately robust relationship between self-report and brain responses to emotion induction

The results from the unsupervised RSA analyses suggest there is a relationship between participants’ self-reported experiences in response to the emotion inductions and their brain responses. However, this relationship was significant for the movie stimuli (Fig. 2-B1) but not for the scenario stimuli (Fig. 2-D1). Supervised multivariate pattern classification of the emotional content from these stimuli further corroborates this finding. The PLS-DA classifiers trained on movie stimuli achieved strong performance (see Fig. 3-A1 and B1 and SI Appendix, Fig. S2-B1), whereas the classifiers trained on scenario stimuli exhibited weak performance (see Fig. 3-A2 and B2 and SI Appendix, Fig. S2-B2), though still above-chance-level accuracy. These results suggest contextually richer movie stimuli may elicit more robust and consistent patterns of brain activity for each emotion, compared to the hypothetical text scenarios. If we assume the scenario stimuli successfully elicited all fifteen target emotions in our participants, which the self-report data indicate (Fig. 1-B2), then there is a possibility that the neural representation of emotions might be modality-dependent and less distinct for text-based scenarios. We note that these scenarios required the participants to be more motivated to imagine themselves in the hypothetical, second-person textual statements and to self-generate the resulting emotional state. Because these processes take time to evolve and vary in temporal onset and duration across participants, the resulting data may be noisier and less amenable to the time-locked, group-averaged signal processing methods employed. During debriefing, participants reported that the scenario task was more tedious, especially given that the task duration was approximately 1.5 hours in the scanner. Inspection of the canonical resting-state network analyses supports the idea that the movies relied relatively more on visual networks whereas the scenarios relied relatively more on the default-mode network, consistent with the modality specificity of the induction methods.

### Neural pattern classification performance tracked predictions of categorical but not dimensional theories

A major theoretical question about emotion representation in the brain is whether emotions can be better predicted using categorical or dimensional factors or a hybrid combination of them (18). The Bayesian model comparison directly compared evidence for these models in predicting the classifier performance on the neural data. For both movies and scenarios, the analysis found strong evidence for the categorical model (or the hybrid model) but relatively weaker evidence for the dimensional model. Although the hybrid model performed nearly as well as the categorical model alone, the parameter estimate for the dimensional model was close to zero, meaning that performance was largely driven by the categorical factor. Furthermore, our PLS-DA decoding error analyses on the movies indicated that pairwise neural distance between emotions is positively correlated with their pairwise distance in categorical space (Fig. 2-B1). This implies that if the (mean) patterns of BOLD activity associated with two emotions are more distinct (as measured by the metric of 1-cosine similarity), then these two emotions tend to be categorized in different ways. However, pairwise neural distance was neither positively nor negatively correlated with pairwise distance in the 2-dimensional valence/arousal space (Fig. 2-B3). These findings provide insight into the putative mechanisms the pattern classification algorithm was relying on to make its predictions, with evidence favoring a categorical architecture, as previously reported by Kragel and LaBar (12) using similar analyses but on a smaller set of emotions. Because our supplemental analyses (SI Appendix, Fig. S3-B1) showed that maximal PLS-DA classifier performance required a minimum of a 7-dimensional latent space to optimally decode the movies, it is likely that a circumplex model of arousal/valence is simply too low-dimensional. The poor decoding performance of the scenarios precludes strong conclusions from similar error analyses; yet we note that the Bayesian model comparison still favored the categorical model in this case. The lower-dimensional space needed for optimal decoding performance of the scenarios (SI Appendix, Fig. S3-B2) may also be tied to the poor decoding performance and simply reflect a weaker neural response or representation. It may be the case that more dimensions would be indicated if the neural encoding were strong to begin with.

### Emotions are organized into intuitive and meaningful clusters at both the subjective and neural levels of representation

The results from hierarchical clustering analysis provide complementary evidence to suggest emotions are organized into both larger-scale and finer-scale clusters that are meaningful, intuitive, yet nuanced, at both the subjective and neural levels of representation. First, we see roughly four clusters based on participants’ categorical endorsement of the emotional stimuli. These clusters are roughly the same for movie (Fig. 4-A1) and scenario stimuli (Fig. 4-A2), for example the disgust–anger-horror cluster for negative emotions and the amusement–joy– excitement cluster for positive emotions. In terms of participants’ valence and arousal ratings of these stimuli, we see two large clusters that are quite distant from each other (Fig. 4-B1 and B2), one for positively valenced emotions and one for negatively valenced emotions. Within each cluster there are, however, smaller clusters. Comparing these clusters from the categorical endorsement and valence/arousal ratings, we see that the clusters from categorical endorsement are more equidistant from each other and fine-grained, whereas the clusters from valence/arousal ratings are more dichotomous. This difference suggests participants are able to make finer distinctions and categorizations of their emotional experiences when presented with emotion labels that may help them tap into the emotion concepts they have already acquired. Other studies in the literature have shown the effectiveness of developing finer emotional granularity through learning more diverse and fine-grained sets of emotion concepts (19). In contrast, participants seem less able to differentiate their emotional experiences when asked to rate the valence/arousal of the emotional stimuli encountered in this study. It also seems that participants tend to rely more on the valence property of the stimuli to make such coarse-grain evaluations, i.e. an emotion is either good or bad, as evidenced by the two large-scale clusters from participants’ valence/arousal ratings that are split along the valence dimension (see Fig. 4-B1 and B2). This bias is consistent with the finding in the literature that people find it harder to report their level of arousal than valence when it comes to affective experiences, perhaps in part due to the multiple varieties of arousal experiences (20). When examining the clusters obtained from participants’ brain activity, we see they resemble more closely the clusters from participants’ categorical endorsement of the emotional stimuli, i.e. the clusters are also less dichotomous and more fined-grained. This pattern suggests the brain representation of emotional experiences may be more categorically distinct. However, this was only found to be the case for movie stimuli (Fig. 4-C1). For scenario stimuli, the clusters from participants’ brain activity were more intermixed and less intuitive (Fig. 4-C2). Not surprisingly, the clusters derived from the confusion matrix of the PLS-DA classifier (Fig. 4-D1) resemble closely the clusters computed from participants’ brain activity. Consequently, these clusters also resemble those computed from participants’ categorical endorsement of the movie stimuli. These results from hierarchical clustering analysis corroborate and provide additional details to the main finding from the RSA analysis, which is that there is a robust relationship between participants’ subjective experience (self-report) of and their brain responses to the emotional stimuli in the current study. Taken together, these data-driven and computational modeling-based findings from our study highlight the categorical, distributed, conceptually distinctive, and high-dimensional nature of our emotional experiences, consistent with parallel observations in support of Semantic Space Theory (7,8,16).

### Limitations and future directions

Readers should keep in mind several limitations when interpreting the results from the current study. The valence/arousal ratings were obtained outside of the scanner. This decision was made for practical considerations due to the length of the study in the scanner (1.5 hours) and the number of stimuli presented (N=150) for each modality. We note that the arousal/valence ratings obtained did correlate with those in the normative database for the stimuli, and that our prior study on a subset of 7 emotions did obtain these ratings in the scanner (12) and yielded similar findings as we did here. Because our supplemental analysis showed that roughly seven dimensions are needed for the PLS-DA classifiers to achieve maximum classification accuracies for decoding movies, future studies could investigate whether a more complex dimensional model may account better for neural classification performance. In order for such a model to be valuable for tracking classification errors, however, it would need to specify how each of the 7 dimensions relates to the others so that pairwise distances between emotions can be calculated. Our classification analyses assumed that each of the 15 categories is independent; however, we acknowledge that there are versions of basic-emotions theories that argue for only a smaller subset of independent emotions while other complex emotions are combinations of the more basic ones, see (14). Another direction for future investigation is how emotions are organized in the body, examining the representational similarity among self-report, brain response, and physiological responses to emotion induction. Finally, future studies should include participants with certain psychiatric conditions such as mood and anxiety disorders in order to investigate differences in emotional organization within clinical populations at either the subjective or neural level. Overall, the results from the current study advance our understanding of how a relatively large set of emotions are organized and represented in the brain, with complementary evidence across supervised and unsupervised modeling approaches that there is a relationship between our subjective experience of and brain responses to emotional inductions that highlight the richness and complexity of human affective experiences.

## Methods

### Participants

The study was approved by the Institutional Review Board (Approval Number: Pro00105943) at Duke University Health System. All participants provided verbal and written informed consent prior to taking part in the study. One hundred sixty six healthy English-speaking participants between 18-49 years old (93 female; mean age = 30.44 ± 8.55) enrolled in the current study. Of these participants, one hundred thirty six (77 female; mean age = 30.35 ± 8.46) completed at least one session of fMRI scanning. Additionally, five pilot participants (data not included in final analyses) were run before the experimental and scan protocol were finalized. Inclusion criteria consisted of having obtained at least a high school diploma or equivalent, no history of neurological illness, and no current psychiatric diagnoses or psychotropic medication use. Participants were excluded if they had a substance abuse disorder, history of breathing or heart-related conditions, intracranial or medical implants, transdermal delivery systems, non-removable piercings, were pregnant or claustrophobic. Each participant was compensated at a rate of 25 USD per hour.

### Experimental Design and Stimuli

The study consisted of three sessions, termed Visit 1-3. Visit 1 and Visit 2 were separated by at least 7 days, and Visit 2 and Visit 3 were separated by at least 5 days. Visit 1 was entirely online and consisted of participants filling out seven surveys: the Affect Intensity Measure (21), Affect Liability Scale (22), Emotion Regulation Questionnaire (23), Penn State Worry Questionnaire (24), Ruminative Response Style (25), State-Trait Anxiety Inventory (26), and the Toronto Alexithymia Scale (27). We note that because we did between-subject (limited by the number of samples from each participant; two blocks per emotion per participant) representational similarity and decoding analyses, we were not able to correlate any metrics from these analyses with individual differences from these psychometric surveys. Visit 2 and 3 were fMRI experiments where participants took part in an emotion induction task while they were in the scanner. We used 150 validated, emotionally-evocative, short (3 to 8 s) movie clips (15) and 150 validated, one- or two-sentence, hypothetical, second-person scenarios (28) for the emotion induction tasks. Both the movie clips and scenarios were divided into 15 emotion categories based on normative ratings (10 stimuli per category per induction method). Five consecutive stimuli of the same emotion category were grouped into a block (two blocks per category per induction method) and presentation order was randomized across emotion blocks. The movie clips and scenarios were presented to the participants in two separate MRI sessions, with one induction method conducted on each day (e.g., movies on Visit 2 and scenarios on Visit 3). For each stimulus trial, participants were first presented with either a movie clip or text scenario on a TV screen for 8 to 13 seconds. They were then instructed to replay the stimulus in their mind for 5 seconds. Afterwards, they responded to a question about where the events of the stimulus took place (indoor/outdoor) via button press. After each emotional block (consisting of 5 trials), they were asked to endorse an emotion they subjectively experienced the most during that block, and to what degree they experienced it on a scale from 1 to 10 (Figure 1-A). Finally, participants were presented with a scrambled black and white image for 8 to 12 seconds, which was referred to as “washout”. See SI Appendix for more detailed descriptions of these stimuli and a more detailed description of the emotion induction task.

### Representational Similarity Analysis (RSA)

The following steps were implemented for representational similarity analysis (29). First, we calculated three types dissimilarity matrices, namely, neural dissimilarity matrix, dissimilarity matrix based on participant categorical endorsement, and dissimilarity matrix based on participant valence and arousal rating for movie and scenario blocks separately. Then, we took the off-diagonal elements of these dissimilarity matrices and compute the corresponding Spearman correlation coefficients. Finally, we computed the bootstrap confidence interval (30) and permutation p values of these Spearman correlation coefficients. See SI for more details.

### Multi-Voxel Pattern Classification

The following steps were taken to decode the emotional content of the movie and scenarios blocks: multiclass to binary classification, data partition for nested cross validation, anomaly detection, feature transformation, computing multiclass classification performance metrics, ensemble averaging and thresholding PLS-DA coefficients for determining voxel importance, and finally post-hoc classification error analysis and Bayesian model comparison. See SI for more details.

### Hierarchical Clustering Analysis

For the exploratory hierarchical clustering analysis, a weighted linkage was applied to the distance matrices from the Representation Similarity Analysis to construct hierarchical cluster trees. To improve interpretability, an optimal leaf order was computed that minimizes pairwise distances between adjacent leaves. Finally, the resulting emotion clusters from each dissimilarity matrix was visualized using the dendrogram function in Matlab (31), which shows visually whether certain emotions tend to cluster together.

## Data, Materials, and Software Availability

All raw data for the study are deposited in the U.S. National Institute of Mental Health Data Archive under protocol \#3639 “Neurocomputational Approaches to Emotion Representation” (PI: Kevin S. LaBar). Code is available on Github: https://github.com/labarlab-emorep-PatternClassify.

## ACKNOWLEDGEMENTS

The current study was funded by the U.S. National Institutes of Health grant RO1 MH124112-01.L. F. was supported by an NSF Graduate Fellowship. The authors also wish to thank Mary Baumann, Shannan Chen, and Sebastian Sanchez for their contribution to participant recruitment and data collection for the current project. The lead author would also like to thank Kevin Kuper and Miles Martinez for their helpful comments on the draft of this manuscript posted on BioRxiv.

## Notes

### Competing Interest Statement

The authors have declared no competing interest.

### Summary of Updates

Introduction, methods, results, and discussion sections have all undergone major revisions. All figures have been updated with new layouts, larger font sizes, and some new plots for more clarity and better visual presentation.

