## Supporting Information for "Representational similarity and pattern classification of fifteen emotional states induced by movie clips and text scenarios"

#### Movie and Scenario Stimuli

Movie stimuli (10 per emotion category) were selected based on normative ratings acquired from (Cowen & Keltner, 2017), as well as additional ratings we collected for new movie clips that were not in the Cowen and Keltner (2017) dataset. New movie clips were especially needed to fill categories such as anger, calmness, excitement, and neutral, which were underrepresented in Cowen and Keltner (2017). Following similar procedures as Cowen and Keltner (2017) and those detailed in Faul et al. (2023), we presented movie clips to participants recruited from Amazon's Mechanical Turk (MTurk) and asked them to select among 34 categories the emotion(s) that best described how they felt when watching each movie (we replaced the boredom category label with neutral). Movie clips were rated by at least 9 individuals in Cowen and Keltner (2017) and at least 11 individuals in our own normative data. We then determined the agreement among participants when choosing category labels (i.e., the concordance rate). Movie clips were only selected for the current study if the concordance rate for the category of interest was greater than 0.5 (reflecting a true majority among raters), and we also required this concordance rate to be greater than any of the other emotion categories in the study. All movie clips lasted between 3-12 seconds (M = 6.8 s, SD = 2.3 s). Although a one-way ANOVA test indicated a significant difference in average movie length across the 15 emotion categories ($F_{14,135} = 1.99$, $p = 0.023$), none of the follow-up pairwise comparisons with a Tukey HSD test were significant (all *p*s > 0.05). Follow-up tests only indicated a marginally shorter duration for the romance category compared to the horror ($p = 0.062$), excitement ($p = 0.075$), and surprise ($p = 0.089$) categories. Thus, movie clips were generally comparable in duration across the 15 emotion categories. The average normative concordance rate for the movie stimuli was 0.73 (SD = 0.13). The category with the highest mean concordance rate was disgust (0.97), and the lowest categories were anxiety and excitement (0.62). We observed a significant difference in concordance rates between categories ($F_{14,135} = 16.89$, $p < .001$). Follow-up pairwise comparisons with a Tukey HSD test revealed that the mean concordance rate for disgust was significantly higher than all other categories except amusement (all *p*s < .05). The mean concordance rates for amusement, sadness, craving, fear, and calmness were also significantly higher when compared to some of the other categories, whereas anxiety, excitement, and awe had comparatively lower concordance rates.

Text scenarios (10 per emotion category) were selected based on data from Faul et al. (2023), who normed custom scenarios as well as scenarios originally created by Fields and Kuperberg (2012). Comprehensive details on these scenarios and their normative data are further detailed in Faul et al. (2023). In brief, all text scenarios were rated by at least 11 MTurk participants who selected emotion categories that best described how they felt when imagining each scenario. As with the movie stimuli, we selected a text scenario for the current study only if the concordance rate was greater than 0.5 for the intended category, and we also ensured that this rate was greater compared to all other categories of interest. All scenarios were written in the second-person perspective and consisted of two sentences (mean words = 19.31, SD = 4.52). With a one-way ANOVA test, we confirmed that the 15 emotion categories did not differ in the number of words per scenario ($F_{14,135} = 1.53$, $p = .108$), average text concreteness ($F_{14,135} = 1.25$, $p = .245$), and average text imageability ($F_{14,135} = 0.79$, $p = .682$). Concreteness and imageability values were obtained with TAALES version 2.0 (Kyle et al., 2018), using metrics from Brysbaert et al. (2014) and the MRC Psycholinguistic Database (Coltheart, 1981), respectively. The average normative concordance rate for the text scenarios was 0.75 (SD = 0.09). The category with the highest mean concordance rate was fear (0.90), and the lowest was neutral (0.66). As with the movie stimuli, we observed a significant difference in concordance rates between categories ($F_{14,135} = 8.45$, $p < .001$). Follow-up comparisons with a Tukey HSD test revealed that the mean concordance rate for fear was significantly higher than all other categories except disgust (all *p*s < .05). The mean concordance rate for disgust was also significantly higher than most other categories, whereas neutral and awe had comparatively lower concordance rates compared to some of the other categories.

#### Task Design

The task was completed over the course of two MRI sessions, separated by at least seven days. The two imaging sessions were identical except for the type of stimulus used: on one day, participants were presented with movie stimuli and on the other day they were presented with text scenarios (order of stimulus type was balanced across participants). Each imaging session contained eight functional task runs. A total of 30 task blocks were presented over the eight runs: runs 1, 2, 3, 5, 6, and 7 contained 4 blocks; runs 4 and 8 contained 3 blocks. Every run began and ended with an 8-second presentation of a fixation cross (small black cross presented in the middle of the screen). Each task block consisted of 5 induction trials followed by emotion endorsement questions and passive viewing of a grayscale washout picture. Each trial contained three parts: passively viewing a movie or reading a text scenario, mentally replaying the movie or scenario, and responding to a question about where the events of the movie or scenario took place. Each movie was presented without sound for its full duration (see below for average movie lengths by emotion) and was immediately followed by the instruction “Replay the movie in your mind”. The “replay” instruction time for each movie was set such that the total movie and “replay” time was always 13 seconds. Each text scenario was displayed for an amount of time equal to the average movie length of its associated emotional induction category (e.g. the average length of anxiety-inducing movie clips was eight seconds, so all anxiety-inducing scenarios were displayed for eight seconds). Text scenario display was immediately followed by the instruction “Replay the scenario in your mind”. As with the movie stimuli, the duration of the “replay” instruction time following each text scenario was set to make the total scenario and “replay” time combine to 13 seconds. Following each “replay” instruction period, a fixation cross was presented for 2 seconds (average, jittered between 1 and 3 seconds). Participants were then presented with the question “Where did the [movie/scenario] take place?” and the possible responses: “1: Indoors”, “2:Outdoors”, “3:Both”, “4:Uncertain”. Participants selected one of these options by pressing the appropriate button on the button box held in their right hand. Participants had 4 seconds to select a response. This question was included to verify participant attention to movie and scenario stimuli while avoiding having participants make explicit on-line ratings of the affective characteristics of the stimuli, since such affect labeling can reduce brain activation in the relevant circuits during emotion tasks (Lieberman et al., 2007).

After the final trial of each block, participants were asked to report the emotion they experienced the most during that block. A selection screen was presented with a list of the 15 emotion options (amusement, anger, anxiety, awe, calmness, craving, disgust, excitement, fear, horror, joy, neutral, romance, sadness, surprise) in a column down the center of the screen and the question “Which of these did you experience most during the last group of movies?”. The order of emotions on the screen was randomized for each presentation. Participants selected an emotion using a selection box that began on a small plus symbol set between the 7th and 8th emotions in the list. Participants used the index finger on the button box to move the selection cursor up the list and the middle finger button to more the selection cursor down the list. Once moved away from the plus symbol in the middle of the list, the cursor would never go back to that position, instead jumping between the 7th and 8th emotions. Participants had 11 seconds to leave the selection cursor on the emotion they wished to select. Following the emotion endorsement screen, participants were asked to rate to what degree they experienced the emotion they selected. The message “You selected [emotion]. How intensely did you experience it?” ([emotion] replaced with the actual emotion chosen) was displayed on the left-hand portion of the screen and the integers 1 through 10 were displayed in a column down the center of the screen in descending order. A plus symbol was displayed below the number 1 and was again the starting position of a selection cursor. Participants used the same button controls to move the cursor up or down to highlight a response. They had 7 seconds to leave the selection cursor on the integer they wished to select. Following emotion intensity endorsement, one of three grayscale “washout” pictures was randomly chosen and displayed for 10 seconds (average, jittered between 8 and 12 seconds). Three neutral pictures (of flowers and trains) were first converted to grayscale and then phase-scrambled. This procedure created pictures which retained the low-level image characteristics of the originals, but contained no discernable objects.

#### MRI Scanner

Magnetic resonance imaging was performed on a Siemens Magnetom 3.0T scanner. The functional task was presented on a MRI-compatible 4K monitor (InroomViewingDevice, NordicNeurolabs, Bergen, Norway) placed at the rear of the scanner. Participants viewed the monitor with a mirror attached to the head coil. Participant responses were collected via a 4-button response box held in the right hand (Current Designs, Inc., Philadelphia, PA).

##### MRI acquisition

We collected T1, BOLD, and resting state fMRI data from each participant. The MRI protocol on each session consisted of a localizer acquisition, a high-resolution T1-weighted acquisition, a reverse-phase-encode-direction (RPED) echo planar imaging (EPI) acquisition, nine EPI acquisitions (8 task runs and a resting-state run), and another RPED EPI acquisition. The T1-weighted acquisition had the following parameters: acquisition matrix = 256 x 256, repetition time = 2250 ms, echo time = 3.12 ms, field of view = 256 mm, in-plane voxel size = 1.0 x 1.0 mm, slice thickness = 1.0 mm, spacing between slices = 0 mm, 192 axial anterior commissure (AC) – posterior commissure (PC) aligned slices. EPI acquisitions had the following parameters: acquisition matrix = 128 x 128, repetition time = 2000 ms, echo time = 30 ms, field of view = 256 x 256 mm, in-plane voxel size = 2.0 x 2.0 mm, slice thickness = 2.0 mm, spacing between slices = 0 mm, phase-encode direction = anterior-to-posterior, 69 axial slices angled 30 degrees positive of AC-PC, multiband acceleration factor = 3, in-plane acceleration factor =2. Runs 1, 2, 3, 5, 6, and 7 of the functional task had 262 time-points and were 8 minutes and 44 seconds long. Runs 4 and 8 had 200 time-points and were 6 minutes and 40 seconds long. The resting state acquisition had 240 volumes and were 8 minutes long. The RPED EPI acquisitions used the same parameters as the functional task EPI acquisitions except a phase-encode direction of posterior-to-anterior and 2 time-points.

##### fMRI Preprocessing

Preprocessing and modeling of functional MRI (fMRI) data were conducted with the following software suites: dcm2niix v1.0.20211006 (Li et al., 2016), Convert 3D v1.1.0 (Yushkevich et al., 2006), FreeSurfer v7.2.0 (Fischl, 2012), fMRIPrep v23.0.2 (Esteban et al., 2019), MRIQC v23.1.0 (Esteban et al., 2017), FMRIB Software Library v6.0.6.4 (Jenkinson et al., 2012), and Analysis of Functional NeuroImages v24.2.01 (Cox, 2012). *Preprocessing.* Anatomic T1-weighted and functional T2*-weighted DICOMs were converted into NIfTI files, and FreeSurfer was then pre-run via the recon-all -all command to generate a set of anatomic parcellations. FMRIPrep next utilized the FreeSurfer output, T2*w run, and T2*w field map files to conduct a workflow which produced preprocessed fMRI data warped into MNI152NLin6Asym template space; see Supplemental Section [S1] for full details. Finally, additional preprocessing was conducted on fMRIPrep output to prepare the fMRI data for first-level modeling. These extra preprocessing steps consisted of bandpass filtering the functional data and then scaling the output such that all fMRI data having a common y-axis range. Additionally, we note that while the ICA-AROMA procedures were used during the fMRIPrep.

#### MRI Preprocessing

Results included in this manuscript come from preprocessing performed using *fMRIPrep* 23.0.2 (Esteban et al., 2018; Esteban et al., 2019), which is based on *Nipype* 1.8.6 (Gorgolewski et al., 2011; Gorgolewski et al., 2018).

"The following boilerplate text was automatically generated by fMRIPrep with the express intention that users should directly copy it into their manuscripts unchanged. It is released under the CC0 license."

##### Preprocessing of $B_0$ inhomogeneity mappings

A total of 9 fieldmaps were found available within the input BIDS structure for this particular subject. A $B_0$-nonuniformity map (or *fieldmap*) was estimated based on two (or more) echo-planar imaging (EPI) references with topup in FSL 6.0.5.1 (Andersson et al., 2003).

#### Anatomical data preprocessing

A total of 1 T1-weighted (T1w) images were found within the input BIDS dataset.The T1-weighted (T1w) image was corrected for intensity non-uniformity (INU) with N4BiasFieldCorrection (Tustison et al., 2010), distributed with ANTs 2.3.3 (Avants et al., 2008), and used as T1w-reference throughout the workflow. The T1w-reference was then skull-stripped with a *Nipype* implementation of the antsBrainExtraction.sh workflow (from ANTs), using MNI152NLin6Asym as target template. Brain tissue segmentation of cerebrospinal fluid (CSF), white-matter (WM) and gray-matter (GM) was performed on the brain-extracted T1w using fast FSL 6.0.5.1 (Zhang et al., 2001). Brain surfaces were reconstructed using recon-all in FreeSurfer 7.3.2 (Dale et al., 1999), and the brain mask estimated previously was refined with a custom variation of the method to reconcile ANTs-derived and FreeSurfer-derived segmentations of the cortical gray-matter of Mindboggle (Klein et al., 2017). Volume-based spatial normalization to two standard spaces (MNI152NLin6Asym, MNI152NLin2009cAsym) was performed through nonlinear registration with antsRegistration (ANTs 2.3.3), using brain-extracted versions of both T1w reference and the T1w template. The following templates were were selected for spatial normalization and accessed with *TemplateFlow* (23.0.0, (Ciric et al., 2022)): *FSL's MNI ICBM 152 non-linear 6th Generation Asymmetric Average Brain Stereotaxic Registration Model (Evans et al., 2012)*; TemplateFlow ID: MNI152NLin6Asym, *ICBM 152 Nonlinear Asymmetrical template version 2009c* (Fonov et al., 2009); TemplateFlow ID: MNI152NLin2009cAsym.

#### Functional data preprocessing

For each of the 9 BOLD runs found per subject (across all tasks and sessions), the following preprocessing was performed. First, a reference volume and its skull-stripped version were generated using a custom methodology of *fMRIPrep*. Head-motion parameters with respect to the BOLD reference (transformation matrices, and six corresponding rotation and translation parameters) are estimated before any spatiotemporal filtering using mcflirt (FSL 6.0.5.1:57b01774, (Jenkinson et al., 2002)). The estimated *fieldmap* was then aligned with rigid-registration to the target EPI (echo-planar imaging) reference run. The field coefficients were mapped on to the reference EPI using the transform. BOLD runs were slice-time corrected to 0.946s (0.5 of slice acquisition range 0s-1.89s) using 3dTshift from AFNI ((Cox & Hyde, 1997),RRID:SCR_005927). The BOLD reference was then co-registered to the T1w reference using bbregister (FreeSurfer) which implements boundary-based registration (Greve & Fischl, 2009). Co-registration was configured with six degrees of freedom. Several confounding time-series were calculated based on the *preprocessed BOLD*: framewise displacement (FD), DVARS and three region-wise global signals. FD was computed using two formulations following (Power et al., 2014) (absolute sum of relative motions, [Power2014]) and Jenkinson et al. (2002) (relative root mean square displacement between affines. FD and DVARS are calculated for each functional run, both using their implementations in *Nipype* (following the definitions by (Power et al., 2014)). The three global signals are extracted within the CSF, the WM, and the whole-brain masks. Additionally, a set of physiological regressors were extracted to allow for component-based noise correction (*CompCor*, (Behzadi et al., 2007)). Principal components are estimated after high-pass filtering the *preprocessed BOLD* time-series (using a discrete cosine filter with 128s cut-off) for the two *CompCor* variants: temporal (tCompCor) and anatomical (aCompCor). tCompCor components are then calculated from the top 2% variable voxels within the brain mask. For aCompCor, three probabilistic masks (CSF, WM and combined CSF+WM) are generated in anatomical space. The implementation differs from that of Behzadi et al. in that instead of eroding the masks by 2 pixels on BOLD space, a mask of pixels that likely contain a volume fraction of GM is subtracted from the aCompCor masks. This mask is obtained by dilating a GM mask extracted from the FreeSurfer's *aseg* segmentation, and it ensures components are not extracted from voxels containing a minimal fraction of GM. Finally, these masks are resampled into BOLD space and binarized by thresholding at 0.99 (as in the original implementation). Components are also calculated separately within the WM and CSF masks. For each CompCor decomposition, the *k* components with the largest singular values are retained, such that the retained components' time series are sufficient to explain 50 percent of variance across the nuisance mask (CSF, WM, combined, or temporal). The remaining components are dropped from consideration. The head-motion estimates calculated in the correction step were also placed within the corresponding confounds file. The confound time series derived from head motion estimates and global signals were expanded with the inclusion of temporal derivatives and quadratic terms for each (Satterthwaite et al., 2013). Frames that exceeded a threshold of 0.5 mm FD or 1.5 standardized DVARS were annotated as motion outliers. Additional nuisance timeseries are calculated by means of principal components analysis of the signal found within a thin band (*crown*) of voxels around the edge of the brain, as proposed by Patriat et al. (2017). The BOLD time-series were resampled into standard space, generating a *preprocessed BOLD run in MNI152NLin6Asym space*. First, a reference volume and its skull-stripped version were generated using a custom methodology of *fMRIPrep*. Automatic removal of motion artifacts using independent component analysis (ICA-AROMA,(Pruim et al., 2015)) was performed on the *preprocessed BOLD on MNI space* time-series after removal of non-steady state volumes and spatial smoothing with an isotropic, Gaussian kernel of 6mm FWHM (full-width half-maximum). Corresponding "non-aggresively" denoised runs were produced after such smoothing. Additionally, the "aggressive" noise-regressors were collected and placed in the corresponding confounds file. All resamplings can be performed with *a single interpolation step* by composing all the pertinent transformations (i.e. head-motion transform matrices, susceptibility distortion correction when available, and co-registrations to anatomical and output spaces). Gridded (volumetric) resamplings were performed using antsApplyTransforms (ANTs), configured with Lanczos interpolation to minimize the smoothing effects of other kernels (Lanczos, 1964). Non-gridded (surface) resamplings were performed using mri_vol2surf (FreeSurfer).

#### Study-specific canonical network mask generation

Masks of seven canonical brain networks (Thomas Yeo et al., 2011), control, default, dorsal attention, limbic, Salience/Ventral Attention, Somatomotor, and Visual, were created from ICA decompositions of the functional resting-state images collected as part of this study. Two sets of network masks were created, one from resting state data collected during movie sessions and one from resting state data collected during scenario sessions. All resting state data were first put through quality assurance. An image was excluded if any visual artifacts were present, more than 10% of the image time-points had FD > 0.5mm, or the maximum volume-to-volume motion exceeded 3.0mm (Birn, 2023). This lead to 30 out of 114 (26%) of movie session resting-state images being excluded and 38 out of 121 (31%) of scenario session resting-state images being excluded. These exclusion percentages are somewhat high. We suspect this is due to the resting-state scan occurring at the end of a relatively long (90-min) imaging session, which may have made it difficult for participants to remain still during acquisition. However, there were still 84 good resting state images from the movie sessions and 83 good resting state images from the scenario sessions. Only participants who contributed task-based data to the classification analysis were included in this resting-state analysis. Following QA, the remaining data sets from the movie sessions were put through FSL's MELODIC group ICA (Beckmann et al., 2005) using temporal concatenation with automatic dimensionality estimation. The threshold for IC maps was set to 0.5. This resulted in a group ICA decomposition comprised of 71 components. The network correspondence toolbox (Kong et al., 2025) was then used to calculate the correspondence of each component with each of the seven canonical networks of interest. For each network, the component with the lowest p-value (or several components if they shared the lowest) was visually inspected against the network mask. For six of the networks (default, dorsal attention, limbic, salience/ventral attention, somatomotor, and visual), a single component was found which well-matched the spatial distribution of the network. The thresholded z-stat images produced by MELODIC for these components were binarized to serve as the participant-derived canonical network masks. For the control network, one component was found which contained the right-lateralized portions of the network and another was found which contained the left-lateralized portions of the network. A mask was created from the union of these two component masks as the participant-derived control network mask. The same process was carried out using the resting state data sets from the scenario sessions. This resulted in an ICA decomposition comprised of 88 components. Five of the canonical networks (default, dorsal attention, salience/ventral attention, somatomotor, and visual) were well represented by single components. Masks for the limbic and control network were each created by taking the union of two component masks.

#### Feature Extraction

First-level models of the functional data were conducted via FSL's feat command (Woolrich et al., 2001), which fits a general linear model (GLM) to each run of the fMRI data. The main regressors were the general task events (indoor/outdoor judgments, washout screens, emotion label selection, and emotion intensity selection), and emotion events (stimulus and replay, modeled separately, for each emotion). Nuisance regressors in the GLM included averaged cerebrospinal fluid (CSF) and white matter (WM) signals, frame-wise motion derivative, and translation and rotation parameters with their derivative), and a confounds file (volumes with a framewise displacement > 0.5 mm were censored). The emotion stimuli blocks vs. washout were the GLM's parameters of interest. All of these regressors were convolved with a double-gamma hemodynamic response function (HRF) before entering into the GLM. Following first-level modeling, the voxel-wise emotion vs. washout parameter was extracted from the respective cope file, resulting in > 900,000 voxel values. This large number of parameters was due both to the spatial resolution of the data as well as the inclusion of extra cerebral, WM, and CSF voxels. To reduce the number of values, then, we masked the data with a gray matter mask (cortical and subcortical) constructed from the template prior tpl-MNI152NLin6Asym_res-02_atlas-HOSPA_desc-th25_dseg. Masking reduced the number of values to approximately 150,000. Masks of seven canonical brain networks (Thomas Yeo et al., 2011) -- control, default, dorsal attention, limbic, Salience/Ventral Attention, Somatomotor, and Visual -- were created from independent component analysis (ICA) decompositions of the functional resting-state images collected as part of this study.

#### Representational Similarity Analysis

**Neural dissimilarity matrix.** Whole-brain (grey matter) voxel-wise beta weights from mass univariate general linear models were used to calculate the neural dissimilarity matrix among the 15 experimentally-induced emotions. In total, we had 3,143 blocks of beta weights from 158,110 voxels for the movie stimuli and 3,247 blocks of beta weights from 158,110 voxels for the scenario stimuli. Each emotion category had roughly 240 blocks. We first averaged the beta weights within each emotion category to get a *mean* neural pattern for that emotion, resulting in 15 vectors, each with a length of 158,100. Then, we computed pairwise cosine similarities among these 15 mean neural patterns and used one minus these pairwise cosine similarities to form the neural dissimilarity matrix. Separate neural dissimilarity matrices are calculated for the neural responses to movie and scenario stimuli. Finally, we used the boxcox function from Matlab to transform the pairwise distances and estimate the transformation parameter lambda. The estimated lambda is 0.0192.

$$pairwise\_dist(\lambda) = \frac{pairwise\_dist^{\lambda} - 1}{\lambda}$$

**Dissimilarity matrix based on participant categorical endorsement**. We defined two types of behavioral dissimilarity matrices from participants' behavioral responses to the emotional stimuli. First, based on participants' categorical endorsement of the emotion blocks that they experienced in the MRI scanner (see Fig. 1A), we computed a confusion matrix (see Fig. 1-B1) by comparing participants' endorsements with the ground truth labels from previous studies (Cowen & Keltner, 2017; Faul et al., 2023). Then we computed pairwise distances (1-cosine similarity) among the 15 row vectors from this confusion matrix. These pairwise distances tell us, on average, how similar or dissimilar the blocks from different emotions are endorsed or categorized by the participants. Again, these pairwise distances were Box-Cox transformed (Box & Cox, 1964) and the estimated transformation parameter lambda is 10.56.

**Dissimilarity matrix based on participant valence and arousal rating**. We calculated another behavioral dissimilarity matrix based on participants' valence and arousal ratings of the movie and scenario stimuli. More specifically, we first averaged the valence and arousal ratings for each stimulus (10 stimuli per emotion category per modality) across the participants. These post-scan ratings are plotted in Fig. 1-C1 & C2. Then, we averaged the valence and arousal ratings within each emotion, i.e., across the stimuli. Therefore, we end up with 15 data points in a 2-D valence and arousal space, one data point for each emotion category. Next, we computed pairwise Euclidean distances among all 15 emotions. We treated this Euclidean distance matrix as comparable to the confusion matrix in the previous section. Then we computed pairwise distances (1-cosine similarity) among the 15 row vectors from this Euclidean distance matrix. Finally, these pairwise distances were Box-Cox transformed, and the estimated transformation parameter lambda is 0.3112.

**Bootstrap confidence interval.** We used a nonparametric bootstrap procedure (Tibshirani & Efron, 1993) to estimate the confidence intervals (CIs) of the representational similarity among the dissimilarity matrices defined in the previous three subsections. For example, we resampled, with replacement, the upper triangular entries of the neural representational dissimilarity matrix (RDM) 5,000 times to generate the bootstrap samples. We then computed the Spearman rank correlation coefficients between the original behavioral RDM and the resampled neural RDM. This procedure gives us a distribution of the Spearman correlation coefficients from these bootstrap resamples. From this distribution, we calculated the 95% confidence interval of the Spearman correlation coefficient using the percentile method (i.e., the 2.5th and 97.5th percentiles).

**Permutation p values.** We used permutation testing (number of permutations = 10,000) to derive the p values of the Spearman correlation coefficients. More specifically, we first permuted the labels of the row vectors of the neural RDM before taking its upper triangular entries. Then we computed the Spearman rank correlation between the permuted neural RDM and behavioral RDMs. All analyses were conducted in MATLAB using custom scripts. We used the following formula to calculate permutation p values:

$$p = \frac{number\left(rho_{permuted} > rho_{observed}\right) + 1}{n_{perm} + 1}$$

#### Multi-voxel Pattern Classification

**Multiclass to binary classification**. We used a one versus all coding scheme to convert the multiclass classification problem into 15 binary classifications, i.e., we trained a different classifier to distinguish each emotion (e.g., amusement) against all the other emotions. In addition to the computational efficiency of this approach, one of the main reasons that we took this approach was because we wanted to identify the voxels (emotion map) that differentiated a given emotion against all the other emotions.

**Nested cross validation.** Overfitting is a common problem in supervised machine learning, especially when the number of predictors is much larger than the number of samples. We used a nested cross-validation scheme to improve the generalizability of our classifiers (Lewis et al., 2023). More specifically, the data were first divided into eight subject independent folds (outer loop). The samples from each subject are only in the same fold, thus preventing information leakage. Each of the seven folds of the data were used as the training data, and the remaining eighth fold was used as the test data. Each training data set was then divided into five folds (inner loop), again in a subject-independent manner. Four of these inner folds were used as training set, and the remaining fifth fold was used as the validation set.

**Anomaly detection.** Outliers are hard to detect in high-dimensional spaces due to their geometric properties. In this study, we used the isolation forest algorithm (Liu et al., 2008) for anomaly detection. Crucially, the algorithm was applied to each training fold from the outer loop after the data had been partitioned according to the nested cross validation scheme described in the previous section. This procedure prevents information leakage. The algorithm outputs an anomaly score for each sample in the data set. We used 0.1% (1 out of 1,000) as the threshold for removing potential outliers; if a sample's anomaly score were in the top 0.1 percentile, then it would be considered as an outlier and removed from the data set.

**Feature transformation.** Voxel-wise beta weights were z-scored before being input to the classifiers to ensure that all voxels were weighted equally before the classifiers were trained. The z scoring was done separately on the training and test data sets to prevent information leakage.

**Partial least squares discriminant analysis.** There are many techniques available when it comes to multi-voxel pattern classification (Kragel et al., 2018). In this study, we chose partial least squares discriminant analysis (PLS-DA) (Li et al., 2018) to be consistent with our prior fMRI decoding study using a smaller number of emotions (Kragel & LaBar, 2015). The input to the classifier consists of beta weights from all emotion blocks (separate for movie and scenario blocks). The outcome variable is the normed label of each emotion block. PLS-DA is a linear supervised classification approach that consists of partial least squares (PLS) and linear discriminant analyses (LDA). During the PLS step, samples were first projected onto a lower-dimensional latent space. Crucially, during this dimensionality reduction step, the labels of the samples were used to weight (or constrain) the covariance matrix of the beta weights. As a result, the components extracted from PLS make the samples from different classes as separable as possible in the latent space, instead of extracting as much variance from the data set as possible. Thus, PLS can be understood as supervised principal component analysis (Ruiz-Perez et al., 2020). Once the samples were projected onto a latent dimensional space, linear discriminant analysis is done to classify samples into different classes. Because PLS is linear, the coefficients (or weights) associated with the voxels are straightforward to interpret. Another advantage of the PLS-DA approach is that there is only one hyperparameter that needs to be tuned, namely the number of latent dimensions.

**Multiclass classification performance metrics.** The predicted scores from each classifier were combined into a single matrix. With fifteen classifiers, we get a matrix of 15 columns of predicted scores. Each row of the matrix represents the predicted scores of the sample from the 15 classifiers. These predicted scores are then transformed into probability scores using a Sigmoid function. To decide which emotion category the sample came from, we used a winner-takes-all approach, i.e., the sample is classified into the class from the classifier with the highest probability score. We repeated this process for each row of the matrix to obtain a vector of predicted emotion labels for all samples. A confusion matrix was then computed by comparing the predicted emotion labels of all the samples with their ground truth labels. From this multiclass confusion matrix, we first counted the following class-specific quantities: true positive, true negative, false positive and false negative. Then we computed the following class specific metrics: accuracy, sensitivity, specificity, and precision. Finally, we computed these macro-averaged multiclass performance metrics, balanced accuracy, macro-averaged sensitivity, macro-averaged specificity, macro-averaged precision, and macro-averaged F1-score, using formulas defined in (Rainio et al., 2024). They are also tabulated in SI *Appendix*, Table S13 for easy reference.

**Bootstrap confidence intervals and permutation testing.** Like the RSA analysis, we used a bootstrap procedure (number of bootstraps = 5,000) to get the confidence intervals of these multiclass classification metrics defined in the previous section. For permutation testing, we randomly shuffled the ground truth labels of all the samples before computing the confusion matrix and subsequently the classification metrics. We repeated this process 5,000 times to get the null distributions of these metrics and calculated their p values using the same equation presented in the RSA section.

**Maps of PLS-DA coefficients.** The PLS-DA coefficients were first z-scored and then averaged across the eight outer folds. This process is repeated for the PLS-DA coefficients from each classifier. Hence, we obtained fifteen vectors of PLS-DA coefficients, one for classifying each emotion. Next, we applied a threshold of 1.96 (corresponding to a two-tailed p value of 0.05; uncorrected) to these vectors and only retained the voxels whose PLS-DA coefficients were greater than this threshold. This ensemble averaging approach gives a good estimate of relative importance of each voxel in classifying each emotion. These supra-threshold voxels were subsequently clustered using FSL's clustering algorithm (Jenkinson et al., 2012). We used a cluster-forming threshold of 30 voxels to reduce clusters that could potentially be false positives. To find the anatomical labels of these clusters, we used the default atlases in FSL and the Harvard Ascending Arousal Network Atlas (Edlow et al., 2023). Finally, the maps of these coefficients were visualized using MRIcroGL (Rorden, 2025).

**Post-hoc classification error analysis and Bayesian model comparison.** Similar to Kragel and LaBar (2015), we conducted a post-hoc analysis to examine whether the classification errors can be explained by theoretical predictions of the participants' patterns of behavioral ratings of the emotional stimuli. More specifically, we represented each row of the confusion matrix from participant in-scanner endorsement ratings as a vector in a 15-dimensional space, where each emotion category is a unique dimension. Then, we computed pairwise distances of these 15 vectors and operationalized these pairwise distances as the distances among these fifteen emotions in a categorical 15-dimensional space. We also defined pairwise distances using the averaged valence and arousal rating of each emotion in a 2-dimensional valence and arousal space (circumplex model). Finally, we examined the Spearman correlation between the classification errors and pairwise distances among these emotions in both the categorical and dimensional spaces. Additionally, we fit four Poisson regression models to the classification errors. The first model only had a constant intercept term. The second model had a constant intercept plus the pairwise distances from the categorical rating as a regressor. The third model had a constant intercept and the pairwise distances from the dimensional rating as an additional regressor. Finally, the fourth model had a constant intercept term and pairwise distances from both the categorical and dimensional model as regressors. Finally, we did a Bayesian model comparison to determine which model explained the classification errors the best.

### Figures

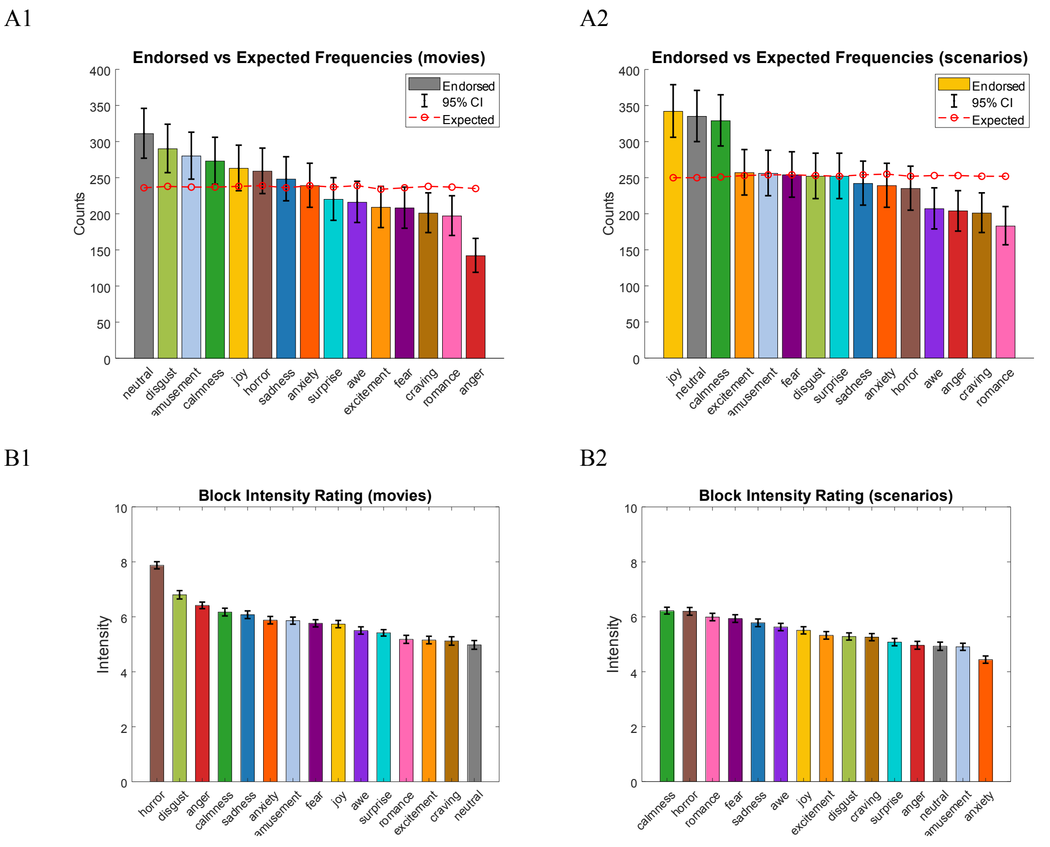


#### Fig. S1.

*A1*, *A2*: Bar plots of endorsed vs expected frequencies of each emotional state from movie and scenario blocks. *B1, B2:* Bar plots of the intensity rating of each emotional state from movie and scenario blocks.

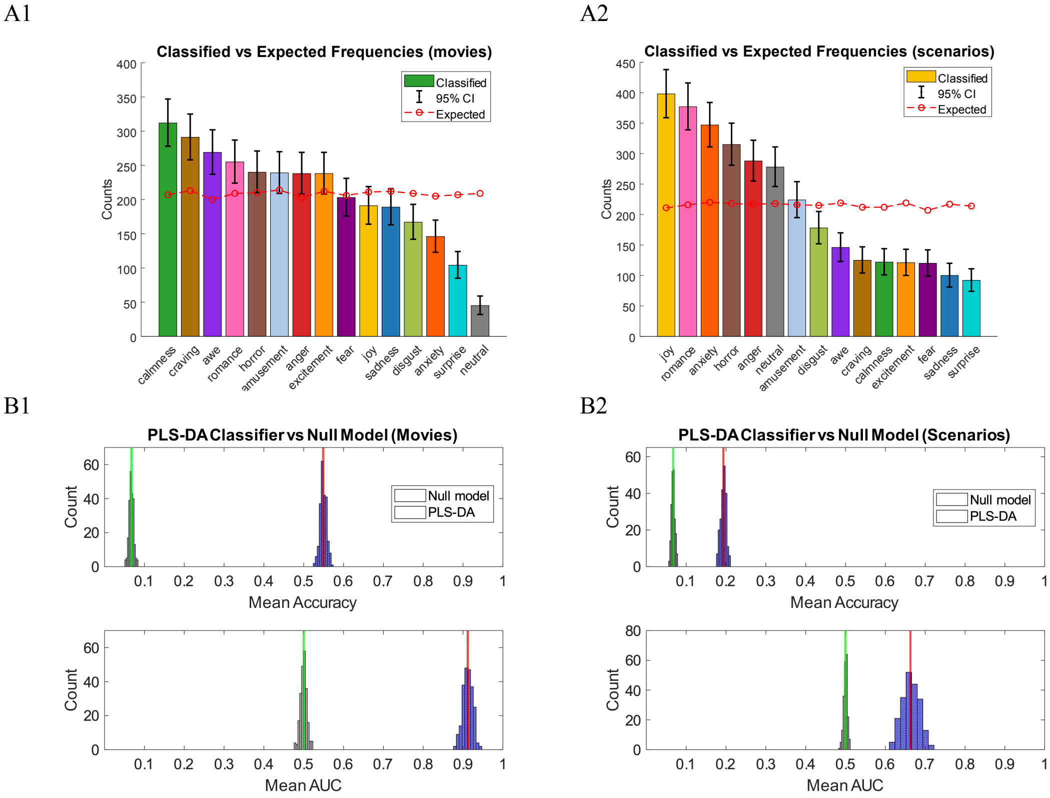


#### Fig. S2.

*A1*, *A2*: Bar plots of classified vs expected frequencies of each emotional state from movie and scenario blocks. *B1*, *B2*: Comparing performance metrics from the PLS-DA classifiers and the null models of movie and scenario blocks.

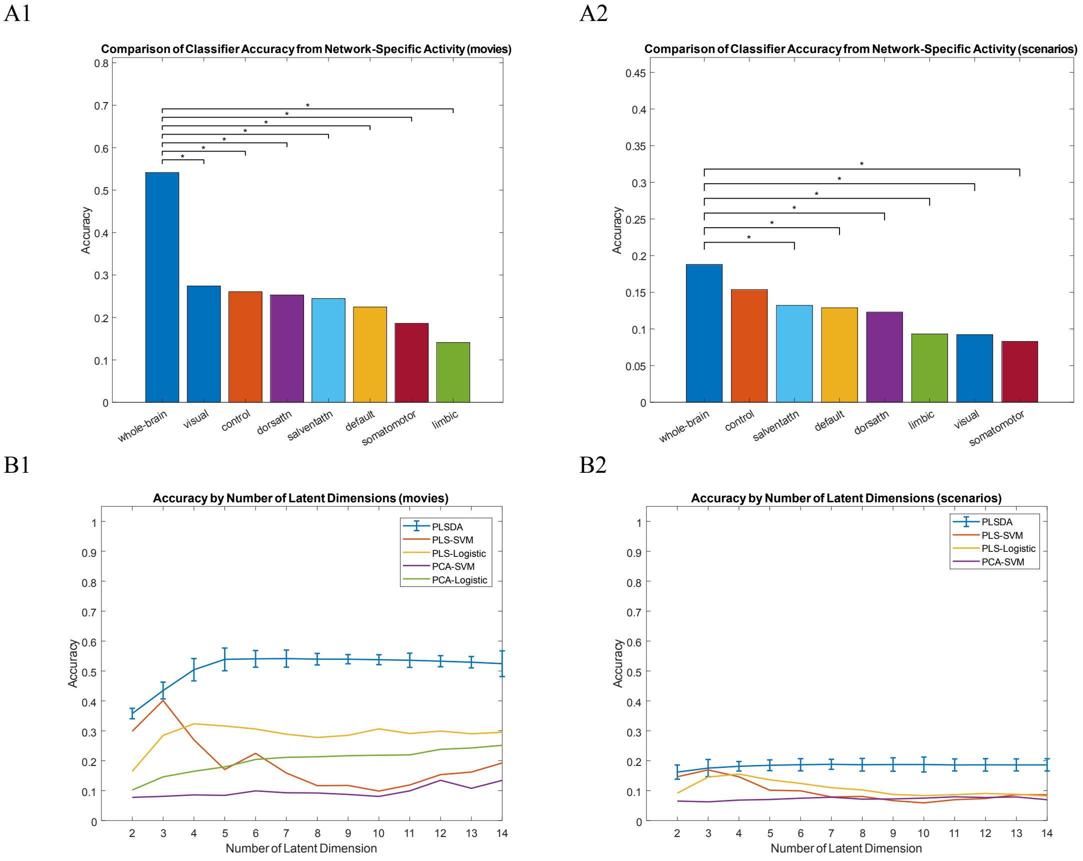


#### Fig. S3.

Results from additional exploratory analyses. *A1*, *A2*: Comparison of classification accuracies from whole-brain patterns of activity vs seven canonical resting state networks for both movie and scenario blocks. *B1, B2:* Classification accuracies as a function of number of latent dimensions for PLS-DA, PLS-SVM, PLS-Logistic, PCA-SVM, and PCA-Logistic classification of movie and scenario blocks. PCA=principle component analysis, PLS-DA=partial least squares discriminant analysis, SVM=support vector machine. * indicates $p < 0.05$ after adjusting for multiple comparisons.

#### Tables

##### Table S1. Performance Metrics of Participant In-Scanner Emotion Endorsement of **Movie** Blocks

| **Movies** | | | | |
|---|---|---|---|---|
| *Metrics* | Estimate | Bootstrap CI | Permutation P value | Chance (Null Distribution) |
| Balanced Accuracy | 0.7395 | [0.7255, 0.7534] | **0.0002*** | 0.0667 |
| Sensitivity (macro-average) | 0.7395 | [0.7255, 0.7534] | **0.0002*** | 0.0667 |
| Specificity (macro-average) | 0.9814 | [0.9803, 0.9824] | **0.0002*** | 0.9333 |
| Precision (macro-average) | 0.7488 | [0.7347, 0.7627] | **0.0002*** | 0.0677 |
| F1-Score (macro-average) | 0.7441 | [0.7303, 0.758] | **0.0002*** | 0.0677 |

Table S2. Emotion Specific Metrics of Participant In-Scanner Endorsement of **Movie** Blocks

| **Movies** | | | | |
|---|---|---|---|---|
| | Sensitivity | Specificity | Precision | F-1 Score |
| Amusement | 0.7542 | 0.9693 | 0.6357 | 0.6899 |
| Anger | 0.4874 | 0.9922 | 0.8169 | 0.6105 |
| Anxiety | 0.6034 | 0.9711 | 0.5983 | 0.6008 |
| Awe | 0.7089 | 0.9855 | 0.7778 | 0.7417 |
| Calmness | 0.9034 | 0.9825 | 0.7875 | 0.8415 |
| Craving | 0.8159 | 0.9982 | 0.9701 | 0.8864 |
| Disgust | 0.9280 | 0.9786 | 0.7552 | 0.8327 |
| Excitement | 0.7029 | 0.9876 | 0.8038 | 0.7500 |
| Fear | 0.4979 | 0.9729 | 0.5673 | 0.5303 |
| Horror | 0.7615 | 0.9768 | 0.7027 | 0.7309 |
| Joy | 0.7265 | 0.9720 | 0.6464 | 0.6841 |
| Neutral | 0.8602 | 0.9675 | 0.6527 | 0.7422 |
| Romance | 0.8067 | 0.9985 | 0.9746 | 0.8828 |
| Sadness | 0.8608 | 0.9867 | 0.8226 | 0.8412 |
| Surprise | 0.6723 | 0.9813 | 0.7182 | 0.6945 |

#### Table S3. Performance Metrics of Participant In–Scanner Emotion Endorsement of **Scenario** Blocks

| **Scenarios** | | | | |
|---|---|---|---|---|
| *Metrics* | Estimate | Bootstrap CI | Permutation P value | Chance (Null Distribution) |
| Balanced Accuracy | 0.7031 | [0.6882, 0.7174] | **0.0002*** | 0.0667 |
| Sensitivity (macro-average) | 0.7031 | [0.6882, 0.7174] | **0.0002*** | 0.0667 |
| Specificity (macro-average) | 0.9788 | [0.9777, 0.9798] | **0.0002*** | 0.9333 |
| Precision (macro-average) | 0.7238 | [0.71, 0.7371] | **0.0002*** | 0.0677 |
| F1-Score (macro-average) | 0.7133 | [0.6992, 0.7268] | **0.0002*** | 0.0677 |

Table S4. Emotion Specific Metrics of Participant In–Scanner Endorsement of **Scenario** Blocks

| **Scenarios** | | | | |
|---|---|---|---|---|
| | Sensitivity | Specificity | Precision | F-1 Score |
| Amusement | 0.6920 | 0.9765 | 0.6758 | 0.6838 |
| Anger | 0.6840 | 0.9907 | 0.8382 | 0.7533 |
| Anxiety | 0.6653 | 0.9796 | 0.6987 | 0.6816 |
| Awe | 0.6680 | 0.9893 | 0.8164 | 0.7348 |
| Calmness | 0.8740 | 0.9697 | 0.6748 | 0.7616 |
| Craving | 0.7520 | 0.9972 | 0.9502 | 0.8396 |
| Disgust | 0.8379 | 0.9887 | 0.8413 | 0.8396 |
| Excitement | 0.5952 | 0.9697 | 0.5837 | 0.5894 |
| Fear | 0.5984 | 0.9711 | 0.5984 | 0.5984 |
| Horror | 0.5961 | 0.9765 | 0.6468 | 0.6204 |
| Joy | 0.6071 | 0.9465 | 0.4474 | 0.5152 |
| Neutral | 0.7549 | 0.9593 | 0.5701 | 0.6497 |
| Romance | 0.6798 | 0.9969 | 0.9399 | 0.7890 |
| Sadness | 0.8016 | 0.9887 | 0.8347 | 0.8178 |
| Surprise | 0.7381 | 0.9813 | 0.7381 | 0.7381 |

#### Table S5. Representational Similarity Between Self-Report and Brain Responses to **Movie** Blocks

| **Movies** | | | | |
|---|---|---|---|---|
| RSA (Spearman's rho) | Estimate | Bootstrap CI | Permutation P value | Chance |
| Neural_vs_Categorical | **0.4113** | [0.2132, 0.5902] | **0.0004** | 0 |
| Neural_vs_Dimensional | 0.0846 | [-0.1193, 0.2859] | 0.3004 | 0 |
| Categorical _vs_Dimensional | **0.5932** | [0.4563, 0.7001] | **0.0004** | 0 |

#### Table S6. Representational Similarity Between Self-Report and Brain Responses to **Scenario** Blocks

| **Scenarios** | | | | |
|---|---|---|---|---|
| RSA (Spearman's rho) | Estimate | Bootstrap CI | Permutation P value | Chance |
| Neural_vs_Categorical | 0.1673 | [-0.02761, 0.3518] | 0.1208 | 0 |
| Neural_vs_Dimensional | 0.1465 | [-0.06299, 0.344] | 0.0562 | 0 |
| Categorical _vs_Dimensional | **0.7509** | [0.6713, 0.8057] | **0.0002** | 0 |

#### Table S7. Performance Metrics of PLSDA Classification of **Movie** Blocks

| **Movies** | | | | |
|---|---|---|---|---|
| *Metrics* | *Estimate* | *Bootstrap CI* | *Permutation P value* | *Chance* |
| Balanced Accuracy | 0.5371 | [0.5208, 0.5533] | **0.0002*** | 0.0667 |
| Sensitivity (macro-average) | 0.5371 | [0.5208, 0.5533] | **0.0002*** | 0.0667 |
| Specificity (macro-average) | 0.967 | [0.9657, 0.9682] | **0.0002*** | 0.9333 |
| Precision (macro-average) | 0.5274 | [0.5076, 0.5469] | **0.0002*** | 0.0677 |
| F1-Score (macro-average) | 0.5322 | [0.5146, 0.5493] | **0.0002*** | 0.0677 |

Table S8. Emotion Specific Metrics of PLSDA Classification of **Movie** Blocks

| **Movies** | | | | |
|---|---|---|---|---|
| | Sensitivity | Specificity | Precision | F-1 Score |
| Amusement | 0.5652 | 0.9582 | 0.4895 | 0.5247 |
| Anger | 0.5869 | 0.9612 | 0.5252 | 0.5543 |
| Anxiety | 0.4450 | 0.9805 | 0.6096 | 0.5145 |
| Awe | 0.5885 | 0.9500 | 0.4572 | 0.5146 |
| Calmness | 0.6619 | 0.9407 | 0.4455 | 0.5326 |
| Craving | 0.8832 | 0.9650 | 0.6495 | 0.7485 |
| Disgust | 0.5813 | 0.9832 | 0.7066 | 0.6378 |
| Excitement | 0.6085 | 0.9626 | 0.5420 | 0.5733 |
| Fear | 0.5534 | 0.9695 | 0.5616 | 0.5575 |
| Horror | 0.6825 | 0.9671 | 0.6000 | 0.6386 |
| Joy | 0.5283 | 0.9729 | 0.5864 | 0.5558 |
| Neutral | 0.0766 | 0.9901 | 0.3556 | 0.1260 |
| Romance | 0.6927 | 0.9613 | 0.5569 | 0.6174 |
| Sadness | 0.4203 | 0.9651 | 0.4603 | 0.4394 |
| Surprise | 0.1818 | 0.9774 | 0.3654 | 0.2428 |

Table S9. Performance Metrics of PLSDA Classification of **Scenario** Blocks

| **Scenarios** | | | | |
|---|---|---|---|---|
| *Metrics* | *Estimate* | *Bootstrap CI* | *Permutation P value* | *Chance* |
| Balanced Accuracy | 0.1862 | [0.1733, 0.199] | **0.0002*** | 0.0667 |
| Sensitivity (macro-average) | 0.1862 | [0.1733, 0.199] | **0.0002*** | 0.0667 |
| Specificity (macro-average) | 0.9418 | [0.9409, 0.9427] | **0.0002*** | 0.9333 |
| Precision (macro-average) | 0.1812 | [0.1663, 0.1963] | **0.0002*** | 0.0677 |
| F1-Score (macro-average) | 0.1837 | [0.1701, 0.1973] | **0.0002*** | 0.0677 |

Table S10. Emotion Specific Metrics of PLSDA Classification of **Scenario** Blocks

| **Scenarios** | | | | |
|---|---|---|---|---|
| | Sensitivity | Specificity | Precision | F-1 Score |
| Amusement | 0.1280 | 0.9348 | 0.1205 | 0.1241 |
| Anger | 0.1759 | 0.9171 | 0.1319 | 0.1508 |
| Anxiety | 0.3273 | 0.9087 | 0.2075 | 0.2540 |
| Awe | 0.1193 | 0.9602 | 0.1781 | 0.1429 |
| Calmness | 0.1244 | 0.9685 | 0.2213 | 0.1593 |
| Craving | 0.0963 | 0.9655 | 0.1680 | 0.1224 |
| Disgust | 0.1852 | 0.9542 | 0.2247 | 0.2030 |
| Excitement | 0.0930 | 0.9665 | 0.1653 | 0.1190 |
| Fear | 0.0868 | 0.9665 | 0.1583 | 0.1121 |
| Horror | 0.4009 | 0.9238 | 0.2698 | 0.3226 |
| Joy | 0.2264 | 0.8841 | 0.1206 | 0.1574 |
| Neutral | 0.1644 | 0.9197 | 0.1295 | 0.1449 |
| Romance | 0.5121 | 0.9104 | 0.2812 | 0.3630 |
| Sadness | 0.0968 | 0.9738 | 0.2100 | 0.1325 |
| Surprise | 0.0561 | 0.9735 | 0.1304 | 0.0784 |

#### Table S11. Correlations between PLSDA Errors and Various Distance Measures (**Movies**)

| **Movies** | | | | |
|---|---|---|---|---|
| Spearman's rho | Estimate | Bootstrap CI | Permutation P value | Chance |
| PLSDA-Error_vs_Categorical | **-0.4308** | [-0.5964, -0.2449] | **0.0004** | 0 |
| PLSDA-Error_vs_ Dimensional | -0.108 | [-0.3352, 0.1087] | 0.2412 | 0 |
| PLSDA-Error _vs_Neural | **-0.7424** | [-0.8187, -0.641] | **0.0002** | 0 |

#### Table S12. Correlations between PLSDA Errors and Various Distance Measures (**Scenarios**)

| **Scenarios** | | | | |
|---|---|---|---|---|
| Spearman's rho | Estimate | Bootstrap CI | Permutation P value | Chance |
| PLSDA-Error_vs_Categorical | **-0.2926** | [-0.4709, -0.09984] | **0.0048** | 0 |
| PLSDA-Error_vs_ Dimensional | -0.1432 | [-0.321, 0.04194] | 0.104 | 0 |
| PLSDA-Error _vs_Neural | **-0.5168** | [-0.6489, -0.3629] | **0.0002** | 0 |

#### Table S13. Formulas for Multiclass Classification Performance Metrics

| | | | |
|---|---|---|---|
| true positive (TP) | $TP_i = n_{ii}$ | true negative (TN) | $TN_i = \sum_{j \neq i} \sum_{h \neq i} n_{jh}$ |
| false positive (FP) | $FP_i = \sum_{j \neq i} n_{ji}$ | false negative (FN) | $FN_i = \sum_{j \neq i} n_{ij}$ |
| accuracy (acc) | $acc_i = \frac{TP_i}{TP_i + FN_i}$ | precision (prec) | $prec_i = \frac{TP_i}{TP_i + FP_i}$ |
| sensitivity (se) | $se_i = \frac{TP_i}{TP_i + FN_i}$ | specificity (sp) | $sp_i = \frac{TN_i}{TN_i + FP_i}$ |
| Balanced Accuracy | $acc_{Balanced} = \sum_{i=1}^{k} acc_i \Big/ k$ | Macro-Averaged Sensitivity | $se_{Macro} = \sum_{i=1}^{k} se_i \Big/ k$ |

| | | | |
|---|---|---|---|
| Macro-Averaged Specificity | $sp_{Macro} = \sum_{i=1}^{k} sp_i \Big/ k$ | Macro-Averaged Precision | $prec_{Macro} = \sum_{i=1}^{k} prec_i \Big/ k$ |
| Macro-Averaged F1-Score | $F1 - Score_{Macro} = 2\frac{(precision_{Macro} * se_{Macro})}{(precision_{Macro} + se_{Macro})}$ | | |

#### Table S14. Maps of PLSDA Coefficients (Movies)

*Amusement*

| Important Clusters from the PLSDA Model Classifying Amusement (movies) | | | | | | |
|---|---|---|---|---|---|---|
| Cluster Size (Voxels) | Max Coef | MNI | | | Hemisphere | Anatomical Label |
| | | x | y | z | | |
| 576 | 4.37 | 6 | -94 | 14 | R | Occipital Pole |
| | 3.99 | 0 | -84 | -6 | C | Lingual Gyrus |
| | 3.73 | -6 | -94 | 2 | L | Occipital Pole |
| 248 | -5.27 | 14 | -80 | 6 | R | Intracalcarine Cortex |
| | -4.39 | 10 | -84 | 4 | R | Intracalcarine Cortex |
| | -3.37 | 10 | -70 | -10 | R | Lingual Gyrus |
| 143 | -3.67 | -22 | -94 | -20 | L | Occipital Pole |
| | -3.52 | -30 | -98 | -10 | L | Occipital Pole |
| | -3.01 | -28 | -100 | -4 | L | Occipital Pole |
| 115 | 5.75 | -30 | 6 | -18 | L | Frontal Orbital Cortex |
| | 4.26 | -42 | 2 | -10 | L | Insular Cortex |
| | 3.8 | -40 | 6 | -14 | L | Insular Cortex |
| 91 | -4 | 56 | -4 | -14 | R | Superior Temporal Gyrus, anterior division |
| | -3.72 | 66 | -4 | -16 | R | Middle Temporal Gyrus, anterior division |
| | -3.14 | 60 | -2 | -10 | R | Superior Temporal Gyrus, anterior division |

| Important Clusters from the PLSDA Model Classifying Amusement (movies) | | | | | | |
|---|---|---|---|---|---|---|
| 90 | -5.82 | 14 | -14 | -20 | R | Right Hippocampus |
| | -5.19 | 20 | -20 | -20 | R | Parahippocampal Gyrus, anterior division |
| | -3.64 | 22 | -18 | -12 | R | Right Hippocampus |
| 84 | -6.57 | -8 | -82 | 2 | L | Intracalcarine Cortex |
| 80 | 3.69 | -40 | -16 | 46 | L | Precentral Gyrus |
| | 3.09 | -38 | -20 | 38 | L | Postcentral Gyrus |
| | 2.96 | -42 | -14 | 34 | L | Postcentral Gyrus |
| 75 | 4.67 | 26 | 8 | -18 | R | Frontal Orbital Cortex |
| | 2.77 | 26 | 14 | -22 | R | Frontal Orbital Cortex |
| | 2.63 | 40 | 8 | -16 | R | Insular Cortex |
| 65 | -3.75 | 12 | -46 | 0 | R | Lingual Gyrus |
| | -3.59 | 14 | -38 | -2 | R | Cingulate Gyrus, posterior division |
| | -2.86 | 18 | -48 | 0 | R | Lingual Gyrus |
| 60 | 4.07 | -46 | 54 | -4 | L | Frontal Pole |
| | 3.46 | -50 | 48 | -6 | L | Frontal Pole |
| | 3.09 | -48 | 52 | 2 | L | Frontal Pole |
| 52 | -2.69 | 30 | -90 | -6 | R | Lateral Occipital Cortex, inferior division / Occipital Pole |
| | -2.61 | 34 | -94 | -8 | R | Occipital Pole |
| | -2.45 | 40 | -90 | -6 | R | Lateral Occipital Cortex, inferior division |
| 52 | -3.29 | -10 | -74 | -10 | L | Lingual Gyrus |
| | -2.95 | -14 | -78 | -8 | L | Lingual Gyrus |
| | -2.5 | -14 | -70 | -12 | L | Lingual Gyrus |
| 50 | -5.43 | 12 | -6 | -26 | R | Parahippocampal Gyrus, anterior division |
| | -4.43 | 18 | -14 | -32 | R | Parahippocampal Gyrus, anterior division |
| | -3.5 | 18 | -6 | -32 | R | Parahippocampal Gyrus, anterior division |
| 44 | 3.78 | -26 | 16 | -24 | L | Frontal Orbital Cortex |
| | 2.66 | -20 | 10 | -26 | L | Frontal Orbital Cortex |
| 42 | 4.67 | 0 | -48 | 14 | C | Cingulate Gyrus, posterior division |
| | 3.81 | 2 | -52 | 20 | R | Cingulate Gyrus, posterior division |
| 42 | 2.99 | -10 | -58 | 74 | L | Lateral Occipital Cortex, superior division |
| | 2.5 | -8 | -54 | 68 | L | Precuneous Cortex |
| | 2.43 | -16 | -62 | 70 | L | Lateral Occipital Cortex, superior division |
| 42 | -3.35 | -2 | 12 | 44 | L | Paracingulate Gyrus |
| | -2.49 | 0 | 18 | 38 | C | Paracingulate Gyrus |

| Important Clusters from the PLSDA Model Classifying Amusement (movies) | | | | | | |
|---|---|---|---|---|---|---|
| | -2.44 | 2 | 10 | 48 | R | Paracingulate Gyrus |
| 41 | -4.57 | -20 | -14 | -14 | L | Left Amygdala |
| | -3.47 | -22 | -20 | -12 | L | Left Hippocampus |
| | -3.04 | -14 | -10 | -14 | L | Left Amygdala |
| 41 | -3.46 | -2 | 34 | -10 | L | Paracingulate Gyrus |
| | -3.07 | -2 | 30 | -16 | L | Subcallosal Cortex |
| | -2.96 | -2 | 24 | -16 | L | Subcallosal Cortex |
| 41 | 3.51 | 0 | -68 | -48 | C | Vermis VIIIb |
| | 3.13 | -2 | -60 | -46 | L | Vermis IX |
| 41 | -3.06 | 0 | -36 | 46 | C | Cingulate Gyrus, posterior division |
| | -2.86 | 0 | -26 | 44 | C | Cingulate Gyrus, posterior division |
| | -2.75 | 0 | -32 | 40 | C | Cingulate Gyrus, posterior division |
| 41 | -3.29 | 36 | -58 | 64 | R | Lateral Occipital Cortex, superior division |
| | -3.19 | 30 | -62 | 66 | R | Lateral Occipital Cortex, superior division |
| | -3.14 | 38 | -52 | 66 | R | Superior Parietal Lobule |
| 40 | -2.82 | -60 | -18 | 18 | L | Central Opercular Cortex |
| | -2.76 | -60 | -20 | 24 | L | Postcentral Gyrus |
| | -2.51 | -58 | -16 | 10 | L | Central Opercular Cortex |
| 39 | 3.8 | 6 | -62 | 66 | R | Precuneous Cortex |
| 38 | -3.35 | -38 | -78 | 14 | L | Lateral Occipital Cortex, superior division |
| | -2.79 | -46 | -78 | 18 | L | Lateral Occipital Cortex, superior division |
| | -2.45 | -36 | -82 | 10 | L | Lateral Occipital Cortex, inferior division |
| 38 | 4.44 | 0 | -56 | -34 | C | Vermis IX |
| 38 | 3.07 | -12 | -44 | 54 | L | Postcentral Gyrus |
| | 2.54 | -16 | -40 | 46 | L | Postcentral Gyrus |
| 37 | 3.18 | -42 | 2 | -44 | L | Temporal Pole |
| | 2.19 | -46 | 0 | -38 | L | Inferior Temporal Gyrus, anterior division |
| 37 | 3.11 | 14 | -42 | 52 | R | Precuneous Cortex |
| | 2.51 | 4 | -40 | 52 | R | Precuneous Cortex |
| 37 | -3.62 | -32 | 12 | 8 | L | Insular Cortex |
| 36 | 3.49 | -16 | -44 | -56 | L | Left VIIIb |
| | 3.09 | -14 | -40 | -50 | L | Left VIIIb |
| 35 | 4.18 | 40 | -78 | -20 | R | Lateral Occipital Cortex, inferior division |
| | 2.69 | 48 | -76 | -18 | R | Lateral Occipital Cortex, inferior division |
| 34 | 3.85 | -40 | -20 | 12 | L | Heschl's Gyrus (includes H1 and H2) |

| Important Clusters from the PLSDA Model Classifying Amusement (movies) | | | | | | |
|---|---|---|---|---|---|---|
| | 2.56 | -42 | -16 | 18 | L | Central Opercular Cortex |
| 33 | -3.16 | -28 | -86 | 24 | L | Lateral Occipital Cortex, superior division |
| | -2.22 | -30 | -78 | 20 | L | Lateral Occipital Cortex, superior division |
| 31 | -2.83 | 48 | 42 | -10 | R | Frontal Pole |
| 31 | -3.83 | 32 | -38 | 72 | R | Postcentral Gyrus |
| | -2.83 | 40 | -38 | 68 | R | Postcentral Gyrus |
| 30 | 4.77 | 12 | -42 | -54 | R | Right VIIIb |

*Anger*

| Important Clusters from the PLSDA Model Classifying Anger (movies) | | | | | | |
|---|---|---|---|---|---|---|
| **Cluster Size (Voxels)** | **Max Coef** | **MNI** | | | **Hemisphere** | **Anatomical Label** |
| | | **x** | **y** | **z** | | |
| 223 | 4.5 | -8 | -98 | -14 | L | Occipital Pole |
| | 4.05 | 10 | -98 | -8 | R | Occipital Pole |
| | 3.99 | 2 | -96 | -2 | R | Occipital Pole |
| 159 | -4.08 | 2 | -40 | 50 | R | Precuneous Cortex |
| | -3.77 | 6 | -38 | 38 | R | Cingulate Gyrus, posterior division |
| | -3.48 | 16 | -36 | 44 | R | Precuneous Cortex |
| 148 | 5.06 | 14 | -32 | -6 | R | Parahippocampal Gyrus, posterior division |
| | 3.93 | 12 | -42 | -2 | R | Lingual Gyrus |
| | 3.47 | 12 | -32 | 0 | R | Right Thalamus |
| 148 | 3.98 | -26 | -10 | 62 | L | Precentral Gyrus |
| | 3.93 | -24 | -10 | 50 | L | Precentral Gyrus |
| | 2.84 | -32 | -4 | 60 | L | Middle Frontal Gyrus |
| 141 | 3.57 | -22 | -62 | -14 | L | Temporal Occipital Fusiform Cortex |
| | 3.29 | -18 | -56 | -10 | L | Lingual Gyrus |
| | 2.92 | -24 | -66 | -10 | L | Occipital Fusiform Gyrus |
| 140 | 3.83 | -2 | -54 | 50 | L | Precuneous Cortex |

| Important Clusters from the PLSDA Model Classifying Anger (movies) | | | | | | |
|---|---|---|---|---|---|---|
| | 3.09 | 2 | -66 | 52 | R | Precuneous Cortex |
| | 2.78 | 2 | -64 | 44 | R | Precuneous Cortex |
| 120 | -3.9 | 0 | 22 | 26 | C | Cingulate Gyrus, anterior division |
| | -3.72 | 2 | 10 | 42 | R | Cingulate Gyrus, anterior division |
| | -3.6 | 0 | 14 | 36 | C | Cingulate Gyrus, anterior division |
| 112 | -4.34 | 34 | -50 | -22 | R | Temporal Occipital Fusiform Cortex |
| | -3.7 | 38 | -68 | -20 | R | Occipital Fusiform Gyrus |
| | -3.6 | 30 | -52 | -18 | R | Temporal Occipital Fusiform Cortex |
| 99 | -4.55 | -54 | -62 | -20 | L | Inferior Temporal Gyrus, temporooccipital part |
| | -3.41 | -40 | -64 | -20 | L | Temporal Occipital Fusiform Cortex |
| | -3.36 | -40 | -72 | -20 | L | Occipital Fusiform Gyrus |
| 91 | 3.47 | -10 | -14 | 10 | L | Left Thalamus |
| | 3.27 | 0 | -14 | 12 | C | Left Thalamus |
| | 3.11 | 0 | -14 | 6 | C | Left Thalamus |
| 79 | 4.05 | 0 | -90 | 32 | C | Occipital Pole |
| | 3.25 | -2 | -94 | 26 | L | Occipital Pole |
| | 3.16 | 4 | -92 | 26 | R | Occipital Pole |
| 77 | -3.37 | -4 | -72 | 10 | L | Intracalcarine Cortex |
| | -3.19 | -2 | -56 | 8 | L | Precuneous Cortex |
| | -2.9 | 4 | -64 | 8 | R | Lingual Gyrus |
| 70 | -3.3 | 0 | -24 | 48 | C | Precentral Gyrus |
| | -3.1 | 0 | -12 | 46 | C | Cingulate Gyrus, anterior division |
| | -2.39 | -2 | -10 | 40 | L | Cingulate Gyrus, anterior division |
| 69 | 3.27 | -56 | -40 | 26 | L | Parietal Operculum Cortex |
| | 2.98 | -64 | -42 | 22 | L | Supramarginal Gyrus, posterior division |
| | 2.92 | -58 | -36 | 22 | L | Parietal Operculum Cortex |
| 68 | 3.95 | 22 | -20 | -16 | R | Right Hippocampus |
| | 3.49 | 24 | -24 | -10 | R | Right Hippocampus |
| | 2.9 | 14 | -12 | -18 | R | Right Hippocampus |
| 67 | -5.44 | -32 | 6 | -20 | L | Temporal Pole |
| | -3.83 | -24 | 12 | -26 | L | Frontal Orbital Cortex |
| | -3.4 | -28 | 4 | -14 | L | Frontal Orbital Cortex |
| 66 | 3.92 | 32 | 14 | -20 | R | Frontal Orbital Cortex |
| | 3.14 | 34 | 22 | -24 | R | Frontal Orbital Cortex |
| | 3.03 | 40 | 14 | -16 | R | Insular Cortex |

| Important Clusters from the PLSDA Model Classifying Anger (movies) | | | | | | |
|---|---|---|---|---|---|---|
| 65 | -2.95 | 60 | -46 | 40 | R | Supramarginal Gyrus, posterior division |
| | -2.73 | 64 | -40 | 42 | R | Supramarginal Gyrus, posterior division |
| | -2.42 | 56 | -38 | 38 | R | Supramarginal Gyrus, posterior division |
| 62 | -3.57 | -42 | -88 | -10 | L | Lateral Occipital Cortex, inferior division |
| | -2.91 | -36 | -90 | -16 | L | Lateral Occipital Cortex, inferior division |
| | -2.56 | -34 | -94 | -12 | L | Occipital Pole |
| 61 | 3.43 | 46 | -28 | -4 | R | Middle Temporal Gyrus, posterior division |
| | 2.97 | 58 | -30 | -2 | R | Middle Temporal Gyrus, posterior division |
| | 2.66 | 50 | -34 | 0 | R | Superior Temporal Gyrus, posterior division |
| 55 | -3.8 | -50 | -40 | 58 | L | Superior Parietal Lobule |
| | -3.44 | -56 | -30 | 54 | L | Supramarginal Gyrus, anterior division |
| | -2.61 | -62 | -34 | 48 | L | Supramarginal Gyrus, anterior division |
| 53 | 4.55 | 14 | -76 | 8 | R | Intracalcarine Cortex |
| 53 | -3.53 | 0 | 10 | 68 | C | Superior Frontal Gyrus / Juxtapositional Lobule Cortex (formerly Supplementary Motor Cortex) |
| | -2.55 | 0 | 6 | 60 | C | Juxtapositional Lobule Cortex (formerly Supplementary Motor Cortex) |
| | -2.51 | 2 | 4 | 68 | R | Juxtapositional Lobule Cortex (formerly Supplementary Motor Cortex) |
| 51 | 3.9 | 12 | -58 | -54 | R | Right VIIIb |
| | 2.26 | 16 | -52 | -52 | R | Right VIIIb |
| | 2.17 | 12 | -56 | -46 | R | Right IX |
| 44 | -3.64 | 64 | -32 | 48 | R | Supramarginal Gyrus, anterior division |
| | -2.67 | 58 | -32 | 54 | R | Supramarginal Gyrus, anterior division |
| | -2.5 | 54 | -38 | 50 | R | Supramarginal Gyrus, posterior division |
| 38 | -2.98 | -10 | -92 | -2 | L | Occipital Pole |
| | -2.88 | -4 | -94 | 0 | L | Occipital Pole |
| 38 | 3.88 | 28 | -40 | -46 | R | Right VIIIb |
| | 2.44 | 32 | -46 | -48 | R | Right VIIIa |
| 37 | -6.17 | 2 | -48 | -44 | R | Right IX |
| | -2.45 | 10 | -46 | -44 | R | Right IX |
| | -2.05 | -4 | -46 | -46 | L | Brain-Stem |
| 37 | 4.32 | -44 | -4 | 8 | L | Central Opercular Cortex |
| | 2.56 | -46 | -8 | 2 | L | Planum Polare |
| 36 | 3.36 | 6 | -84 | -14 | R | Lingual Gyrus |
| | 2.78 | 12 | -78 | -14 | R | Lingual Gyrus |

| Important Clusters from the PLSDA Model Classifying Anger (movies) | | | | | | |
|---|---|---|---|---|---|---|
| 36 | -5.33 | 2 | -52 | -58 | R | Right IX |
| | -3.91 | 8 | -54 | -58 | R | Right IX |
| | -2.95 | 6 | -54 | -64 | R | Right IX |
| 35 | 3.06 | -40 | 22 | -18 | L | Frontal Orbital Cortex |
| | 2.61 | -36 | 14 | -18 | L | Frontal Orbital Cortex |
| 35 | -2.9 | 62 | -20 | 46 | R | Supramarginal Gyrus, anterior division |
| | -2.82 | 60 | -22 | 52 | R | Postcentral Gyrus |
| | -2.15 | 64 | -30 | 42 | R | Supramarginal Gyrus, anterior division |
| 35 | -3.3 | 54 | 8 | 46 | R | Middle Frontal Gyrus |
| | -2.53 | 52 | 6 | 52 | R | Middle Frontal Gyrus |
| 32 | 3.76 | -2 | 10 | -18 | L | Subcallosal Cortex |
| | 2.17 | 0 | 8 | -8 | C | Subcallosal Cortex |
| 32 | -2.64 | -24 | -28 | 60 | L | Precentral Gyrus |
| | -2.4 | -26 | -32 | 64 | L | Postcentral Gyrus |
| 30 | -3.01 | -2 | -60 | -34 | L | Vermis VIIIa |

*Anxiety*

| Important Clusters from the PLSDA Model Classifying Anxiety (movies) | | | | | | |
|---|---|---|---|---|---|---|
| Cluster Size (Voxels) | Max Coef | MNI | | | Hemisphere | Anatomical Label |
| | | x | y | z | | |
| 1155 | -5.87 | 14 | -96 | 24 | R | Occipital Pole |
| | -5.87 | -2 | -96 | 16 | L | Occipital Pole |
| | -5.86 | 8 | -98 | 14 | R | Occipital Pole |
| 591 | 7.53 | 10 | -76 | -4 | R | Lingual Gyrus |
| | 6.18 | 12 | -80 | 4 | R | Intracalcarine Cortex |
| | 4.25 | 2 | -76 | 2 | R | Lingual Gyrus |
| 124 | -3.74 | -32 | -60 | -18 | L | Temporal Occipital Fusiform Cortex |
| | -3.72 | -24 | -48 | -16 | L | Temporal Occipital Fusiform Cortex |
| | -3.71 | -30 | -54 | -18 | L | Temporal Occipital Fusiform Cortex |
| 111 | -4 | 0 | -60 | -36 | C | Vermis VIIIb |
| | -3.8 | 0 | -52 | -38 | C | Vermis IX |
| | -3.46 | 4 | -58 | -40 | R | Right IX |
| 93 | 5.36 | -24 | -46 | -8 | L | Lingual Gyrus |
| | 3.42 | -22 | -40 | -12 | L | Parahippocampal Gyrus, posterior division |
| | 3.27 | -16 | -42 | -14 | L | Parahippocampal Gyrus, posterior division |
| 93 | 3.36 | -2 | -36 | 48 | L | Cingulate Gyrus, posterior division |
| | 3.29 | 0 | -24 | 44 | C | Cingulate Gyrus, posterior division |
| | 2.85 | 0 | -30 | 46 | C | Cingulate Gyrus, posterior division |
| 76 | 3.71 | -52 | 18 | -4 | L | Inferior Frontal Gyrus, pars opercularis |
| | 3.58 | -38 | 22 | -2 | L | Insular Cortex |
| | 2.67 | -44 | 18 | -6 | L | Frontal Orbital Cortex |
| 70 | -4.39 | 58 | -16 | 12 | R | Central Opercular Cortex |
| | -3.54 | 48 | -18 | 14 | R | Central Opercular Cortex |
| | -2.73 | 40 | -18 | 16 | R | Central Opercular Cortex |
| 65 | -3.49 | 38 | 4 | 34 | R | Precentral Gyrus |
| | -3.37 | 42 | 10 | 30 | R | Precentral Gyrus |
| 63 | -4.76 | 18 | 2 | -22 | R | Parahippocampal Gyrus, anterior division |
| | -4.26 | 20 | 0 | -16 | R | Right Amygdala |
| 62 | -3.48 | 10 | -48 | -10 | R | Right I-IV |
| | -3.36 | -2 | -48 | -2 | L | Left I-IV |
| | -3.17 | -8 | -40 | -12 | L | Left I-IV |
| 62 | 5.58 | -4 | -14 | -26 | L | Brain-Stem |

| Important Clusters from the PLSDA Model Classifying Anxiety (movies) | | | | | | |
|---|---|---|---|---|---|---|
| | 5.18 | 2 | -14 | -28 | R | Brain-Stem |
| | 3.57 | -12 | -16 | -26 | L | Brain-Stem |
| 60 | -4.14 | 40 | -4 | 4 | R | Insular Cortex |
| | -3.34 | 40 | 2 | -6 | R | Insular Cortex |
| | -2.8 | 44 | -8 | 2 | R | Insular Cortex |
| 54 | 2.98 | -12 | -92 | -4 | L | Occipital Pole |
| | 2.75 | -14 | -98 | -4 | L | Occipital Pole |
| | 2.12 | -12 | -92 | -10 | L | Occipital Pole |
| 52 | 3.68 | -18 | -84 | -18 | L | Occipital Fusiform Gyrus |
| | 3.44 | -24 | -84 | -20 | L | Occipital Fusiform Gyrus |
| | 2.31 | -20 | -88 | -22 | L | Occipital Fusiform Gyrus |
| 52 | 4.09 | 20 | -20 | -22 | R | Parahippocampal Gyrus, anterior division |
| | 2.93 | 22 | -26 | -28 | R | Parahippocampal Gyrus, posterior division |
| | 2.81 | 16 | -14 | -20 | R | Right Hippocampus |
| 52 | -4.43 | -8 | -74 | -10 | L | Lingual Gyrus |
| | -2.8 | -14 | -76 | -14 | L | Lingual Gyrus |
| 51 | -3.59 | 44 | 38 | 12 | R | Frontal Pole |
| | -2.39 | 40 | 34 | 14 | R | Frontal Pole |
| | -2.23 | 38 | 38 | 10 | R | Frontal Pole |
| 49 | -3.51 | 26 | -82 | -18 | R | Occipital Fusiform Gyrus |
| | -3.46 | 30 | -74 | -18 | R | Occipital Fusiform Gyrus |
| | -2.66 | 28 | -68 | -12 | R | Occipital Fusiform Gyrus |
| 48 | -5.09 | -40 | 6 | -12 | L | Insular Cortex |
| | -3.8 | -46 | 2 | -14 | L | Planum Polare |
| 48 | -2.92 | 48 | -32 | 58 | R | Postcentral Gyrus |
| | -2.47 | 48 | -26 | 48 | R | Postcentral Gyrus |
| 48 | -3.54 | -26 | -82 | -10 | L | Occipital Fusiform Gyrus |
| | -3.49 | -32 | -74 | -18 | L | Occipital Fusiform Gyrus |
| | -2.5 | -30 | -72 | -10 | L | Occipital Fusiform Gyrus |
| 48 | -3.14 | -38 | 36 | 14 | L | Frontal Pole |
| | -2.76 | -40 | 30 | 16 | L | Inferior Frontal Gyrus, pars triangularis |
| | -2.37 | -48 | 38 | 14 | L | Frontal Pole |
| 41 | -3.77 | 30 | 10 | -22 | R | Temporal Pole |
| | -3.6 | 36 | 10 | -20 | R | Temporal Pole |
| | -3.57 | 34 | 8 | -14 | R | Insular Cortex |

| Important Clusters from the PLSDA Model Classifying Anxiety (movies) | | | | | | |
|---|---|---|---|---|---|---|
| 41 | -3.95 | 36 | -26 | -20 | R | Temporal Fusiform Cortex, posterior division |
| | -2.95 | 28 | -22 | -16 | R | Right Hippocampus |
| 40 | 3.14 | -60 | -26 | 50 | L | Postcentral Gyrus |
| | 2.71 | -60 | -24 | 44 | L | Postcentral Gyrus |
| 37 | -3 | 0 | -76 | 38 | C | Precuneous Cortex |
| | -2.93 | 8 | -78 | 46 | R | Precuneous Cortex |
| 35 | 4.09 | -4 | -24 | 4 | L | Left Thalamus |
| | 2.24 | -8 | -20 | 8 | L | Left Thalamus |
| 33 | 3.78 | 50 | 20 | -6 | R | Inferior Frontal Gyrus, pars triangularis |
| 32 | -3.04 | -30 | -70 | 50 | L | Lateral Occipital Cortex, superior division |
| 31 | 3.01 | 56 | -72 | 2 | R | Lateral Occipital Cortex, inferior division |
| | 2.91 | 52 | -68 | 6 | R | Lateral Occipital Cortex, inferior division |
| 30 | 2.73 | -58 | -38 | 22 | L | Parietal Operculum Cortex |
| | 2.07 | -64 | -36 | 22 | L | Parietal Operculum Cortex |
| 30 | -2.75 | 30 | -50 | -20 | R | Temporal Occipital Fusiform Cortex |
| | -2.71 | 24 | -52 | -18 | R | Right VI |

*Awe*

| Important Clusters from the PLSDA Model Classifying Awe (movies) | | | | | | |
|---|---|---|---|---|---|---|
| **Cluster Size (Voxels)** | **Max Coef** | **MNI** | | | **Hemisphere** | **Anatomical Label** |
| | | **x** | **y** | **z** | | |
| 448 | -5.86 | 54 | -70 | 6 | R | Lateral Occipital Cortex, inferior division |
| | -5.05 | 54 | -66 | 12 | R | Lateral Occipital Cortex, inferior division |
| | -4.41 | 52 | -74 | 10 | R | Lateral Occipital Cortex, inferior division |
| 381 | -4.14 | -6 | -88 | -20 | L | Lingual Gyrus |
| | -4.05 | 4 | -72 | 0 | R | Lingual Gyrus |
| | -4.03 | 0 | -74 | 6 | C | Lingual Gyrus |
| 227 | 4.77 | -40 | 18 | -16 | L | Frontal Orbital Cortex |
| | 4.33 | -50 | 20 | -6 | L | Inferior Frontal Gyrus, pars triangularis |
| | 3.56 | -56 | 14 | 0 | L | Inferior Frontal Gyrus, pars opercularis |
| 210 | -6.36 | 18 | -4 | -16 | R | Right Amygdala |
| | -4.61 | 24 | 4 | -18 | R | Parahippocampal Gyrus, anterior division |
| | -3.89 | 16 | -14 | -22 | R | Right Hippocampus |
| 201 | -4.13 | 42 | -74 | -20 | R | Lateral Occipital Cortex, inferior division |
| | -4.11 | 38 | -54 | -16 | R | Temporal Occipital Fusiform Cortex |
| | -3.61 | 42 | -50 | -22 | R | Temporal Occipital Fusiform Cortex |
| 200 | -5.07 | -52 | -76 | 10 | L | Lateral Occipital Cortex, inferior division |
| | -3.34 | -48 | -76 | 4 | L | Lateral Occipital Cortex, inferior division |
| | -3.19 | -46 | -82 | 6 | L | Lateral Occipital Cortex, inferior division |
| 160 | 4 | 40 | -86 | 12 | R | Lateral Occipital Cortex, superior division |
| | 3.2 | 38 | -78 | 6 | R | Lateral Occipital Cortex, inferior division |
| | 3 | 38 | -90 | 8 | R | Occipital Pole |
| 144 | -3.66 | 58 | -42 | 14 | R | Supramarginal Gyrus, posterior division |
| | -3.31 | 50 | -38 | 10 | R | Supramarginal Gyrus, posterior division |
| | -2.99 | 64 | -40 | 8 | R | Supramarginal Gyrus, posterior division |
| 142 | -3.5 | -26 | -56 | -8 | L | Temporal Occipital Fusiform Cortex |
| | -3.39 | -16 | -58 | -10 | L | Lingual Gyrus |
| | -3.23 | -20 | -64 | -12 | L | Lingual Gyrus |
| 139 | 8.55 | 14 | -80 | 6 | R | Intracalcarine Cortex |
| | 3.22 | 16 | -94 | 0 | R | Occipital Pole |
| | 2.57 | 10 | -74 | 12 | R | Intracalcarine Cortex |
| 126 | -4.49 | -46 | -68 | -22 | L | Occipital Fusiform Gyrus |
| | -3.16 | -42 | -44 | -20 | L | Temporal Fusiform Cortex, posterior division |

| Important Clusters from the PLSDA Model Classifying Awe (movies) | | | | | | |
|---|---|---|---|---|---|---|
| | -3.12 | -40 | -50 | -24 | L | Temporal Occipital Fusiform Cortex |
| 122 | 6.03 | -10 | -82 | 4 | L | Intracalcarine Cortex |
| | 2.65 | 0 | -88 | 8 | C | Supracalcarine Cortex |
| | 2.19 | -6 | -90 | -2 | L | Occipital Pole |
| 91 | 3.82 | 54 | -62 | -14 | R | Lateral Occipital Cortex, inferior division |
| | 3.53 | 52 | -58 | -10 | R | Inferior Temporal Gyrus, temporooccipital part |
| | 3.04 | 46 | -60 | -8 | R | Inferior Temporal Gyrus, temporooccipital part |
| 89 | 4.15 | 22 | -76 | 56 | R | Lateral Occipital Cortex, superior division |
| | 3.07 | 26 | -60 | 46 | R | Lateral Occipital Cortex, superior division |
| | 2.85 | 16 | -78 | 56 | R | Lateral Occipital Cortex, superior division |
| 84 | 4.78 | -14 | -42 | -14 | L | Left V |
| | 3.87 | -22 | -36 | -14 | L | Parahippocampal Gyrus, posterior division |
| | 3.8 | -8 | -38 | -16 | L | Left I-IV |
| 80 | 3.81 | -34 | -88 | 8 | L | Lateral Occipital Cortex, superior division |
| | 3.01 | -30 | -88 | 18 | L | Lateral Occipital Cortex, superior division |
| 77 | 3 | -12 | -100 | -14 | L | Occipital Pole |
| | 2.66 | -14 | -98 | -4 | L | Occipital Pole |
| | 2.27 | -6 | -96 | -6 | L | Occipital Pole |
| 64 | -4.34 | -10 | -30 | -4 | L | Brain-Stem |
| | -4.01 | -10 | -24 | 0 | L | Left Thalamus |
| | -3.15 | -4 | -26 | -6 | L | Brain-Stem |
| 64 | -2.77 | 44 | 0 | 46 | R | Precentral Gyrus |
| | -2.44 | 42 | 0 | 54 | R | Middle Frontal Gyrus |
| | -2.28 | 36 | -4 | 50 | R | Precentral Gyrus |
| 60 | 3.03 | 12 | -54 | 16 | R | Precuneous Cortex |
| | 2.91 | 12 | -58 | 22 | R | Precuneous Cortex |
| | 2.53 | 0 | -64 | 24 | C | Precuneous Cortex |
| 59 | 3.92 | 30 | -64 | 34 | R | Lateral Occipital Cortex, superior division |
| | 2.36 | 28 | -72 | 38 | R | Lateral Occipital Cortex, superior division |
| 49 | 4.07 | 12 | -66 | -6 | R | Lingual Gyrus |
| | 3.35 | 6 | -66 | -2 | R | Lingual Gyrus |
| 42 | -3.62 | -38 | 4 | -14 | L | Insular Cortex |
| | -3.34 | -38 | 10 | -18 | L | Temporal Pole |
| 41 | -3.62 | -24 | -26 | -30 | L | Parahippocampal Gyrus, posterior division |
| | -2.79 | -30 | -26 | -26 | L | Parahippocampal Gyrus, posterior division |

| Important Clusters from the PLSDA Model Classifying Awe (movies) | | | | | | |
|---|---|---|---|---|---|---|
| | -2.41 | -34 | -28 | -20 | L | Temporal Fusiform Cortex, posterior division |
| 40 | 3.76 | 10 | -50 | -6 | R | Lingual Gyrus |
| | 2.75 | 16 | -48 | -6 | R | Lingual Gyrus |
| 39 | 2.64 | -24 | -72 | -52 | L | Left VIIb |
| | 2.61 | -20 | -68 | -48 | L | Left VIIb |
| | 2.44 | -14 | -68 | -46 | L | Left VIIIa |
| 37 | -7.13 | -4 | -14 | -26 | L | Brain-Stem |
| | -5.33 | 4 | -14 | -26 | R | Brain-Stem |
| | -2.25 | -10 | -16 | -26 | L | Brain-Stem |
| 36 | -4.2 | -18 | -4 | -12 | L | Left Amygdala |
| | -3.04 | -22 | -12 | -12 | L | Left Amygdala |
| 34 | -3.4 | -2 | -56 | -36 | L | Vermis IX |
| 33 | 4.33 | 32 | 10 | -20 | R | Temporal Pole |
| | 3.23 | 38 | 18 | -20 | R | Temporal Pole |
| 33 | -3.89 | -38 | -76 | -20 | L | Occipital Fusiform Gyrus |
| | -3.19 | -34 | -82 | -20 | L | Occipital Fusiform Gyrus |
| 33 | -3.56 | -48 | -24 | 14 | L | Parietal Operculum Cortex |
| | -3.3 | -42 | -30 | 14 | L | Parietal Operculum Cortex |
| 33 | -4.83 | 4 | 8 | -20 | R | Subcallosal Cortex |
| | -2.74 | 12 | 8 | -20 | R | Frontal Orbital Cortex |
| 30 | -3.3 | -16 | 6 | -24 | L | Frontal Orbital Cortex |
| | -2.71 | -22 | 8 | -18 | L | Frontal Orbital Cortex |
| | -2.62 | -22 | 2 | -16 | L | Parahippocampal Gyrus, anterior division |

*Calmness*

| Important Clusters from the PLSDA Model Classifying Calmness (movies) | | | | | | |
|---|---|---|---|---|---|---|
| Cluster Size (Voxels) | Max Coef | MNI | | | Hemisphere | Anatomical Label |
| | | x | y | z | | |
| 220 | -5.01 | -42 | -78 | -20 | L | Lateral Occipital Cortex, inferior division |
| | -4.36 | -46 | -76 | -12 | L | Lateral Occipital Cortex, inferior division |
| | -4.05 | -40 | -84 | -20 | L | Lateral Occipital Cortex, inferior division |
| 127 | 5.01 | -20 | 0 | -20 | L | Left Amygdala |
| | 4.04 | -24 | 12 | -32 | L | Temporal Pole |
| | 3.9 | -20 | 6 | -30 | L | Temporal Pole |
| 111 | -4.86 | 10 | -30 | -14 | R | Brain-Stem |
| | -4.78 | 14 | -30 | -4 | R | Right Thalamus |
| | -4.54 | 0 | -28 | -2 | C | Brain-Stem |
| 89 | -3.6 | -54 | -32 | 22 | L | Parietal Operculum Cortex |
| | -3.11 | -62 | -40 | 28 | L | Supramarginal Gyrus, anterior division / Supramarginal Gyrus, posterior division |
| | -2.8 | -48 | -40 | 26 | L | Parietal Operculum Cortex |
| 88 | -3.6 | 6 | -88 | -10 | R | Lingual Gyrus |
| | -3.35 | 2 | -84 | -12 | R | Lingual Gyrus |
| | -2.93 | 6 | -76 | -14 | R | Lingual Gyrus |
| 83 | 4.52 | 26 | -8 | -20 | R | Right Hippocampus |
| | 3.31 | 20 | -16 | -18 | R | Right Hippocampus |
| | 3.24 | 22 | -14 | -12 | R | Right Amygdala |
| 81 | -4.56 | -12 | -24 | 40 | L | Cingulate Gyrus, posterior division |
| | -3.26 | -8 | -20 | 44 | L | Cingulate Gyrus, posterior division |
| | -2.97 | 0 | -22 | 46 | C | Cingulate Gyrus, posterior division |
| 81 | 4.25 | 0 | -82 | 0 | C | Lingual Gyrus |
| | 3.19 | -4 | -92 | -2 | L | Occipital Pole |
| | 2.68 | 2 | -76 | -4 | R | Lingual Gyrus |
| 76 | 3.32 | 22 | -102 | -4 | R | Occipital Pole |
| | 3.19 | 26 | -98 | -10 | R | Occipital Pole |
| | 2.61 | 24 | -96 | 0 | R | Occipital Pole |
| 70 | -4.11 | 48 | -64 | 6 | R | Lateral Occipital Cortex, inferior division |
| | -3.24 | 50 | -72 | 2 | R | Lateral Occipital Cortex, inferior division |
| | -2.56 | 56 | -62 | 4 | R | Lateral Occipital Cortex, inferior division |
| 64 | -5.82 | -6 | -36 | -58 | L | Brain-Stem |
| | -3.77 | 6 | -38 | -62 | R | Brain-Stem |

| Important Clusters from the PLSDA Model Classifying Calmness (movies) | | | | | | |
|---|---|---|---|---|---|---|
| | -2.82 | -4 | -38 | -46 | L | Brain-Stem |
| 54 | 4.21 | -66 | -26 | 12 | L | Superior Temporal Gyrus, posterior division |
| | 3.3 | -66 | -26 | 2 | L | Superior Temporal Gyrus, posterior division |
| 54 | -2.85 | 12 | -90 | 30 | R | Occipital Pole |
| | -2.73 | 24 | -84 | 28 | R | Lateral Occipital Cortex, superior division |
| | -2.62 | 20 | -90 | 24 | R | Occipital Pole |
| 52 | 3.86 | 10 | -42 | -4 | R | Lingual Gyrus |
| | 3.34 | 14 | -52 | 2 | R | Lingual Gyrus |
| | 3.3 | 14 | -42 | -10 | R | Lingual Gyrus |
| 49 | -3.53 | 52 | 10 | 48 | R | Middle Frontal Gyrus |
| | -2.73 | 50 | 8 | 42 | R | Middle Frontal Gyrus |
| | -2.66 | 50 | 6 | 54 | R | Middle Frontal Gyrus |
| 47 | -3.79 | 38 | -68 | -18 | R | Occipital Fusiform Gyrus |
| | -3.73 | 38 | -64 | -12 | R | Occipital Fusiform Gyrus |
| 46 | -4.03 | -10 | -22 | 10 | L | Left Thalamus |
| | -3.78 | -12 | -16 | 0 | L | Left Thalamus |
| | -3.26 | -12 | -16 | 6 | L | Left Thalamus |
| 45 | -4.64 | 14 | -82 | 50 | R | Lateral Occipital Cortex, superior division |
| | -2.47 | 10 | -88 | 44 | R | Occipital Pole |
| 45 | -3.24 | -6 | -50 | -36 | L | Left IX |
| | -2.74 | 0 | -54 | -36 | C | Vermis IX |
| | -2.61 | -2 | -60 | -36 | L | Vermis VIIIb |
| 42 | 3.48 | 6 | 2 | -14 | R | Right Cerebral Cortex |
| | 3.25 | 16 | 4 | -22 | R | Parahippocampal Gyrus, anterior division |
| | 2.89 | 10 | 6 | -18 | R | Frontal Orbital Cortex |
| 42 | -3.22 | -46 | 6 | 36 | L | Middle Frontal Gyrus |
| | -3.15 | -52 | 10 | 38 | L | Middle Frontal Gyrus |
| | -2.25 | -48 | 0 | 38 | L | Precentral Gyrus |
| 41 | 3.2 | -2 | -78 | 32 | L | Cuneal Cortex |
| | 2.97 | 2 | -72 | 34 | R | Precuneous Cortex |
| 39 | -5.91 | 0 | 10 | -14 | C | Subcallosal Cortex |
| 39 | 3.23 | 32 | -88 | 4 | R | Lateral Occipital Cortex, inferior division |
| | 2.47 | 32 | -94 | 4 | R | Occipital Pole |
| | 2.45 | 32 | -88 | -4 | R | Lateral Occipital Cortex, inferior division |
| 38 | -3.28 | 28 | -82 | 42 | R | Lateral Occipital Cortex, superior division |

| Important Clusters from the PLSDA Model Classifying Calmness (movies) | | | | | | |
|---|---|---|---|---|---|---|
| | -2.65 | 22 | -80 | 46 | R | Lateral Occipital Cortex, superior division |
| 37 | 2.85 | -10 | -78 | -4 | L | Lingual Gyrus |
| | 2.76 | -14 | -76 | -8 | L | Lingual Gyrus |
| 36 | 3.05 | 44 | -78 | 28 | R | Lateral Occipital Cortex, superior division |
| | 2.89 | 48 | -70 | 26 | R | Lateral Occipital Cortex, superior division |
| 35 | -3.09 | 2 | -72 | -26 | R | Vermis VI |
| | -2.74 | 0 | -76 | -30 | C | Vermis Crus II |
| | -2.07 | -4 | -72 | -36 | L | Left Crus II |
| 33 | 3.03 | -30 | -96 | -2 | L | Occipital Pole |
| | 2.34 | -34 | -96 | -10 | L | Occipital Pole |
| 33 | -3.2 | 2 | 52 | 36 | R | Superior Frontal Gyrus |
| | -3.05 | 4 | 58 | 40 | R | Frontal Pole |
| | -2.38 | -2 | 56 | 32 | L | Superior Frontal Gyrus |
| 32 | -3.17 | -2 | 36 | 38 | L | Paracingulate Gyrus |
| | -3.03 | 2 | 32 | 36 | R | Paracingulate Gyrus |
| 32 | 4.04 | 0 | 8 | -8 | C | Subcallosal Cortex |
| | 3.33 | 0 | 14 | 0 | C | Subcallosal Cortex |
| 32 | 3.53 | -12 | -58 | 16 | L | Precuneous Cortex |
| 30 | 4.28 | -24 | -14 | -12 | L | Left Amygdala |
| | 3.67 | -18 | -18 | -14 | L | Left Hippocampus |
| | 2.79 | -28 | -18 | -14 | L | Left Hippocampus |
| 30 | 2.63 | -42 | -22 | 64 | L | Postcentral Gyrus |
| | 2.39 | -40 | -24 | 56 | L | Postcentral Gyrus |
| | 2.25 | -46 | -14 | 60 | L | Precentral Gyrus |
| 30 | -4.33 | 2 | -68 | -4 | R | Lingual Gyrus |
| | -3.87 | 4 | -62 | 0 | R | Lingual Gyrus |
| 30 | -2.91 | 54 | -4 | -16 | R | Superior Temporal Gyrus, anterior division |

*Craving*

| Important Clusters from the PLSDA Model Classifying Craving (movies) | | | | | | |
|---|---|---|---|---|---|---|
| Cluster Size (Voxels) | Max Coef | MNI | | | Hemisphere | Anatomical Label |
| | | x | y | z | | |
| 552 | 4.64 | 62 | -14 | 28 | R | Postcentral Gyrus |
| | 4.08 | 64 | -14 | 38 | R | Postcentral Gyrus |
| | 4 | 50 | -28 | 44 | R | Supramarginal Gyrus, anterior division |
| 497 | 5.03 | -26 | -66 | -8 | L | Occipital Fusiform Gyrus |
| | 4.71 | -24 | -74 | -18 | L | Occipital Fusiform Gyrus |
| | 4.62 | -28 | -60 | -18 | L | Temporal Occipital Fusiform Cortex |
| 430 | 5.2 | -60 | -28 | 50 | L | Supramarginal Gyrus, anterior division |
| | 4.57 | -62 | -22 | 44 | L | Postcentral Gyrus |
| | 3.77 | -52 | -36 | 56 | L | Supramarginal Gyrus, anterior division |
| 272 | 4.73 | 30 | -54 | -8 | R | Temporal Occipital Fusiform Cortex |
| | 3.77 | 26 | -72 | -6 | R | Occipital Fusiform Gyrus |
| | 3.4 | 30 | -62 | -8 | R | Occipital Fusiform Gyrus |
| 236 | 6.84 | 40 | -2 | 0 | R | Insular Cortex |
| | 5.91 | 40 | 6 | -12 | R | Insular Cortex |
| | 5.84 | 40 | 2 | -6 | R | Insular Cortex |
| 116 | -3.58 | 44 | -78 | 28 | R | Lateral Occipital Cortex, superior division |
| | -2.97 | 50 | -76 | 26 | R | Lateral Occipital Cortex, superior division |
| | -2.91 | 44 | -66 | 18 | R | Lateral Occipital Cortex, superior division |
| 113 | 4.82 | -40 | 6 | -12 | L | Insular Cortex |
| | 4.68 | -38 | -6 | 4 | L | Insular Cortex |
| | 4.15 | -38 | -2 | -2 | L | Insular Cortex |
| 102 | 3.53 | -18 | -62 | 66 | L | Lateral Occipital Cortex, superior division |
| | 3.12 | -30 | -66 | 60 | L | Lateral Occipital Cortex, superior division |
| | 2.85 | -18 | -68 | 64 | L | Lateral Occipital Cortex, superior division |
| 98 | -4.59 | -18 | -44 | -12 | L | Lingual Gyrus |
| | -2.72 | -26 | -48 | -6 | L | Lingual Gyrus |
| | -2.2 | -22 | -52 | -8 | L | Lingual Gyrus |
| 94 | -3.69 | 20 | -56 | 12 | R | Precuneous Cortex |
| | -3.49 | 20 | -68 | 24 | R | Cuneal Cortex |
| | -3.22 | 22 | -58 | 20 | R | Precuneous Cortex |
| 91 | -3.7 | -38 | -12 | -6 | L | Insular Cortex |
| | -3.58 | -38 | -22 | 0 | L | Insular Cortex |

| Important Clusters from the PLSDA Model Classifying Craving (movies) | | | | | | |
|---|---|---|---|---|---|---|
| | -3.06 | -46 | -8 | -4 | L | Planum Polare |
| 91 | 4.94 | 34 | -26 | -22 | R | Temporal Fusiform Cortex, posterior division |
| | 4.05 | 32 | -32 | -22 | R | Temporal Fusiform Cortex, posterior division |
| | 3.15 | 34 | -20 | -24 | R | Parahippocampal Gyrus, anterior division |
| 81 | -3.64 | -4 | -30 | 4 | L | Left Thalamus |
| | -3.33 | -6 | -24 | 0 | L | Left Thalamus |
| | -3.32 | -2 | -30 | -2 | L | Brain-Stem |
| 79 | -3.85 | -62 | -58 | -2 | L | Middle Temporal Gyrus, temporooccipital part |
| 76 | -3.35 | 40 | -66 | -20 | R | Occipital Fusiform Gyrus |
| | -2.72 | 42 | -50 | -24 | R | Temporal Occipital Fusiform Cortex |
| | -2.64 | 40 | -60 | -14 | R | Temporal Occipital Fusiform Cortex |
| 75 | 4.56 | -32 | -26 | -22 | L | Parahippocampal Gyrus, posterior division |
| | 2.88 | -28 | -34 | -22 | L | Temporal Fusiform Cortex, posterior division |
| | 2.3 | -32 | -22 | -28 | L | Parahippocampal Gyrus, anterior division |
| 70 | 3.95 | -44 | -70 | 4 | L | Lateral Occipital Cortex, inferior division |
| | 2.77 | -50 | -72 | 10 | L | Lateral Occipital Cortex, inferior division |
| | 2.03 | -56 | -68 | 14 | L | Lateral Occipital Cortex, superior division |
| 63 | 3.19 | 48 | -64 | 2 | R | Lateral Occipital Cortex, inferior division |
| | 2.86 | 46 | -62 | 10 | R | Lateral Occipital Cortex, inferior division |
| | 2.76 | 54 | -58 | 18 | R | Angular Gyrus |
| 63 | -3.04 | -8 | -90 | 4 | L | Occipital Pole |
| | -2.24 | -10 | -92 | -2 | L | Occipital Pole |
| 61 | 4.55 | 12 | -6 | -20 | R | Right Hippocampus |
| | 3.62 | 18 | -2 | -22 | R | Right Amygdala |
| | 2.69 | 14 | -8 | -26 | R | Parahippocampal Gyrus, anterior division |
| 60 | -4.72 | 2 | -8 | -2 | R | Right Thalamus |
| | -2.55 | 0 | -6 | 4 | C | Left Thalamus |
| | -2.36 | -4 | 0 | 0 | L | Left Thalamus |
| 53 | -3.37 | 50 | 0 | 56 | R | Precentral Gyrus |
| | -2.67 | 48 | -8 | 58 | R | Precentral Gyrus |
| | -2.23 | 40 | 0 | 64 | R | Middle Frontal Gyrus |
| 52 | -4.15 | 20 | -22 | -18 | R | Parahippocampal Gyrus, anterior division / Parahippocampal Gyrus, posterior division |
| | -2.69 | 26 | -12 | -18 | R | Right Hippocampus |
| 51 | -2.96 | 8 | -16 | 44 | R | Cingulate Gyrus, posterior division |

| Important Clusters from the PLSDA Model Classifying Craving (movies) | | | | | | |
|---|---|---|---|---|---|---|
| | -2.76 | 0 | -22 | 38 | C | Cingulate Gyrus, posterior division |
| | -2.69 | 6 | -22 | 44 | R | Cingulate Gyrus, posterior division |
| 49 | 3.46 | 0 | -84 | -4 | C | Lingual Gyrus |
| | 2.7 | -2 | -84 | -10 | L | Lingual Gyrus |
| | 2.56 | -10 | -86 | -16 | L | Occipital Fusiform Gyrus |
| 48 | -4.06 | 14 | -6 | 78 | R | Superior Frontal Gyrus |
| | -2.71 | 20 | -4 | 74 | R | Superior Frontal Gyrus |
| | -2.04 | 8 | -4 | 74 | R | Superior Frontal Gyrus |
| 43 | 3.78 | -32 | -58 | 68 | L | Superior Parietal Lobule |
| | 2.79 | -36 | -54 | 66 | L | Superior Parietal Lobule |
| 41 | -3.61 | -30 | 18 | -16 | L | Frontal Orbital Cortex |
| | -2.93 | -36 | 14 | -16 | L | Insular Cortex |
| | -2.43 | -42 | 18 | -16 | L | Frontal Orbital Cortex |
| 39 | 4.4 | -14 | -4 | -20 | L | Left Amygdala |
| 38 | -3.08 | 2 | 12 | 54 | R | Paracingulate Gyrus |
| | -2.4 | 0 | 18 | 54 | C | Superior Frontal Gyrus |
| 38 | -3.5 | -12 | -46 | -4 | L | Lingual Gyrus |
| | -3.5 | 0 | -48 | -4 | C | Left I-IV |
| | -2.72 | -6 | -46 | -4 | L | Left I-IV |
| 37 | 3.07 | 8 | -70 | 12 | R | Intracalcarine Cortex |
| | 2.95 | 12 | -74 | 14 | R | Intracalcarine Cortex |
| | 2.36 | 10 | -64 | 10 | R | Intracalcarine Cortex |
| 36 | -3.16 | 2 | 46 | 36 | R | Superior Frontal Gyrus |
| 33 | -3.33 | 40 | -12 | -6 | R | Insular Cortex |
| | -2.13 | 44 | -6 | -4 | R | Insular Cortex |
| 32 | 3.54 | 22 | -8 | -40 | R | Parahippocampal Gyrus, anterior division |
| | 3.32 | 18 | -6 | -36 | R | Parahippocampal Gyrus, anterior division |
| 32 | -3.94 | -30 | -86 | 38 | L | Lateral Occipital Cortex, superior division |
| | -2.09 | -30 | -90 | 30 | L | Occipital Pole |
| 32 | -3.44 | 60 | -26 | 18 | R | Parietal Operculum Cortex |
| 30 | -4.36 | -16 | -18 | -22 | L | Parahippocampal Gyrus, anterior division |
| | -2.99 | -18 | -24 | -22 | L | Parahippocampal Gyrus, posterior division |

*Disgust*

| Important Clusters from the PLSDA Model Classifying Disgust (movies) | | | | | | |
|---|---|---|---|---|---|---|
| Cluster Size (Voxels) | Max Coef | MNI | | | Hemisphere | Anatomical Label |
| | | x | y | z | | |
| 620 | 6.34 | -18 | -6 | -14 | L | Left Amygdala |
| | 6.04 | -26 | 2 | -18 | L | Parahippocampal Gyrus, anterior division |
| | 5.78 | -38 | 4 | -14 | L | Insular Cortex |
| 421 | 6.04 | 22 | 0 | -16 | R | Right Amygdala |
| | 5.37 | 18 | -6 | -14 | R | Right Amygdala |
| | 4.56 | 12 | -8 | -16 | R | Right Amygdala |
| 412 | 5.04 | 62 | -16 | 14 | R | Central Opercular Cortex |
| | 4.37 | 58 | -14 | 18 | R | Central Opercular Cortex |
| | 4.31 | 64 | -18 | 26 | R | Postcentral Gyrus |
| 244 | 4.81 | -60 | -20 | 30 | L | Postcentral Gyrus |
| | 3.66 | -66 | -22 | 30 | L | Postcentral Gyrus |
| | 3.6 | -64 | -18 | 20 | L | Postcentral Gyrus |
| 233 | -6.14 | -8 | -84 | 2 | L | Intracalcarine Cortex |
| | -4.9 | -6 | -90 | -2 | L | Occipital Pole |
| | -4.56 | -6 | -78 | 6 | L | Intracalcarine Cortex |
| 170 | -7.74 | 14 | -80 | 6 | R | Intracalcarine Cortex |
| | -3.87 | 16 | -90 | 4 | R | Occipital Pole |
| 112 | 3.27 | -28 | -86 | 38 | L | Lateral Occipital Cortex, superior division |
| | 3.24 | -28 | -70 | 28 | L | Lateral Occipital Cortex, superior division |
| | 3.05 | -26 | -80 | 42 | L | Lateral Occipital Cortex, superior division |
| 106 | -3.87 | 12 | -76 | -6 | R | Lingual Gyrus |
| | -3.38 | 12 | -74 | -12 | R | Lingual Gyrus |
| 104 | -3.69 | 38 | -30 | 18 | R | Parietal Operculum Cortex |
| | -3.56 | 44 | -30 | 22 | R | Parietal Operculum Cortex |
| | -3.11 | 38 | -18 | 20 | R | Central Opercular Cortex |
| 93 | -4.32 | -12 | -76 | -14 | L | Lingual Gyrus |
| | -3.2 | -14 | -78 | -8 | L | Lingual Gyrus |
| 87 | 5.08 | 0 | -6 | 6 | C | Left Thalamus |
| | 4.55 | 2 | -20 | 4 | R | Right Thalamus |
| | 2.79 | 0 | -14 | 8 | C | Left Thalamus |
| 86 | 3.64 | 2 | 24 | 34 | R | Paracingulate Gyrus / Cingulate Gyrus, anterior division |
| | 3.34 | 0 | 14 | 36 | C | Cingulate Gyrus, anterior division |

| Important Clusters from the PLSDA Model Classifying Disgust (movies) | | | | | | |
|---|---|---|---|---|---|---|
| | 3.08 | 0 | 22 | 26 | C | Cingulate Gyrus, anterior division |
| 82 | -3.79 | -44 | -24 | 20 | L | Parietal Operculum Cortex |
| | -3.45 | -48 | -28 | 14 | L | Parietal Operculum Cortex |
| | -3.04 | -40 | -22 | 14 | L | Central Opercular Cortex |
| 70 | 2.91 | -56 | -62 | -16 | L | Inferior Temporal Gyrus, temporooccipital part |
| | 2.9 | -50 | -56 | -14 | L | Inferior Temporal Gyrus, temporooccipital part |
| | 2.74 | -60 | -56 | -16 | L | Inferior Temporal Gyrus, temporooccipital part |
| 61 | -4.64 | 10 | -46 | 0 | R | Lingual Gyrus |
| | -2.98 | 6 | -42 | 8 | R | Cingulate Gyrus, posterior division |
| | -2.89 | 16 | -50 | 4 | R | Cingulate Gyrus, posterior division |
| 58 | -3.41 | -58 | -38 | 34 | L | Supramarginal Gyrus, anterior division |
| | -3.2 | -52 | -40 | 30 | L | Supramarginal Gyrus, posterior division |
| | -2.88 | -46 | -36 | 24 | L | Parietal Operculum Cortex |
| 48 | -5.22 | 18 | -38 | -14 | R | Lingual Gyrus |
| | -3.77 | 12 | -34 | -8 | R | Parahippocampal Gyrus, posterior division |
| 48 | 2.98 | 24 | 30 | -18 | R | Frontal Orbital Cortex |
| | 2.87 | 24 | 20 | -24 | R | Frontal Orbital Cortex |
| 45 | 2.88 | 46 | -10 | 50 | R | Precentral Gyrus |
| | 2.51 | 38 | -14 | 40 | R | Precentral Gyrus |
| | 2.35 | 42 | -12 | 44 | R | Precentral Gyrus |
| 44 | -6.33 | -12 | -16 | -30 | L | Brain-Stem |
| 43 | 3.23 | 6 | 6 | 56 | R | Juxtapositional Lobule Cortex (formerly Supplementary Motor Cortex) |
| | 2.96 | 2 | 6 | 62 | R | Juxtapositional Lobule Cortex (formerly Supplementary Motor Cortex) |
| 43 | -2.7 | 52 | -66 | 4 | R | Lateral Occipital Cortex, inferior division |
| | -2.64 | 46 | -68 | 2 | R | Lateral Occipital Cortex, inferior division |
| | -2.53 | 50 | -72 | -2 | R | Lateral Occipital Cortex, inferior division |
| 42 | -3.27 | 4 | -12 | 58 | R | Juxtapositional Lobule Cortex (formerly Supplementary Motor Cortex) |
| 42 | -3.4 | 34 | -4 | 60 | R | Precentral Gyrus |
| | -2.73 | 32 | -8 | 70 | R | Precentral Gyrus |
| 42 | -3.75 | -48 | -10 | 28 | L | Postcentral Gyrus |
| | -2.61 | -58 | -8 | 22 | L | Postcentral Gyrus |
| | -2.42 | -54 | -6 | 26 | L | Precentral Gyrus |
| 37 | -3.56 | 18 | -16 | -18 | R | Right Hippocampus |

| Important Clusters from the PLSDA Model Classifying Disgust (movies) | | | | | | |
|---|---|---|---|---|---|---|
| | -2.23 | 20 | -10 | -22 | R | Right Hippocampus |
| | -2.17 | 24 | -18 | -16 | R | Right Hippocampus |
| 37 | 3.64 | 32 | -34 | -16 | R | Parahippocampal Gyrus, posterior division |
| | 3.24 | 28 | -40 | -20 | R | Temporal Occipital Fusiform Cortex |
| 36 | 3.28 | -24 | 34 | -14 | L | Frontal Orbital Cortex |
| | 2.71 | -28 | 36 | -20 | L | Frontal Pole |
| 36 | -3.04 | -46 | -70 | 10 | L | Lateral Occipital Cortex, inferior division |
| | -2.59 | -50 | -72 | 6 | L | Lateral Occipital Cortex, inferior division |
| 36 | -3.17 | 64 | -2 | 22 | R | Precentral Gyrus |
| 35 | -3 | 18 | -44 | -50 | R | Right VIIIb |
| | -2.64 | 26 | -46 | -52 | R | Right VIIIb |
| 32 | 2.55 | 0 | -68 | 16 | C | Supracalcarine Cortex |
| | 2.45 | -4 | -72 | 14 | L | Intracalcarine Cortex |
| | 2.4 | 4 | -78 | 14 | R | Supracalcarine Cortex |
| 31 | -3.59 | 0 | 16 | -10 | C | Subcallosal Cortex |
| 30 | 3.1 | -38 | -64 | -18 | L | Temporal Occipital Fusiform Cortex |
| 30 | -2.62 | 26 | -10 | 56 | R | Precentral Gyrus |
| | -2.6 | 28 | -8 | 48 | R | Precentral Gyrus |
| 30 | -2.84 | -4 | 40 | -12 | L | Paracingulate Gyrus |
| | -2.17 | 0 | 48 | -10 | C | Frontal Medial Cortex |

*Excitement*

| Important Clusters from the PLSDA Model Classifying Excitement (movies) | | | | | | |
|---|---|---|---|---|---|---|
| Cluster Size (Voxels) | Max Coef | MNI | | | Hemisphere | Anatomical Label |
| | | x | y | z | | |
| 213 | 7.28 | 10 | -84 | 2 | R | Intracalcarine Cortex |
| | 3.96 | 14 | -92 | 0 | R | Occipital Pole |
| 163 | -5.82 | -2 | -22 | 12 | L | Left Thalamus |
| | -5.19 | -2 | -4 | -2 | L | Left Thalamus |
| | -5.16 | -4 | -28 | 8 | L | Left Thalamus |
| 145 | 4.09 | 30 | -50 | -8 | R | Temporal Occipital Fusiform Cortex |
| | 3.09 | 24 | -56 | -8 | R | Lingual Gyrus |
| | 3.08 | 18 | -46 | -12 | R | Lingual Gyrus |
| 142 | 6.61 | -8 | -84 | 2 | L | Intracalcarine Cortex |
| 141 | 4.44 | -54 | -64 | -18 | L | Lateral Occipital Cortex, inferior division |
| | 4.19 | -36 | -78 | -20 | L | Occipital Fusiform Gyrus |
| | 4.02 | -42 | -76 | -20 | L | Lateral Occipital Cortex, inferior division |
| 136 | 3.98 | -30 | -54 | -18 | L | Temporal Occipital Fusiform Cortex |
| | 3.5 | -28 | -48 | -16 | L | Temporal Occipital Fusiform Cortex |
| | 2.94 | -24 | -56 | -10 | L | Temporal Occipital Fusiform Cortex |
| 99 | -3.83 | 2 | -72 | 2 | R | Lingual Gyrus |
| | -3.41 | -6 | -72 | -2 | L | Lingual Gyrus |
| | -2.94 | 0 | -76 | -4 | C | Lingual Gyrus |
| 86 | -6.19 | -38 | -24 | 12 | L | Heschl's Gyrus (includes H1 and H2) |
| | -3.59 | -42 | -16 | 4 | L | Heschl's Gyrus (includes H1 and H2) |
| | -2.86 | -46 | -10 | 2 | L | Heschl's Gyrus (includes H1 and H2) |
| 76 | 4.87 | -12 | -22 | 40 | L | Cingulate Gyrus, posterior division |
| | 3.31 | -10 | -16 | 44 | L | Precentral Gyrus |
| 68 | 3.21 | -56 | -44 | -10 | L | Middle Temporal Gyrus, temporooccipital part |
| | 3.1 | -60 | -52 | -10 | L | Middle Temporal Gyrus, temporooccipital part |
| | 2.65 | -62 | -56 | -14 | L | Inferior Temporal Gyrus, temporooccipital part |
| 66 | 3.23 | -48 | -38 | 28 | L | Parietal Operculum Cortex |
| | 3.05 | -42 | -36 | 18 | L | Parietal Operculum Cortex |
| | 3.01 | -50 | -38 | 20 | L | Planum Temporale |
| 65 | -4.59 | -34 | 10 | -24 | L | Temporal Pole |
| | -3.81 | -30 | 14 | -22 | L | Frontal Orbital Cortex |
| | -3.54 | -34 | 18 | -26 | L | Frontal Orbital Cortex |

| Important Clusters from the PLSDA Model Classifying Excitement (movies) | | | | | | |
|---|---|---|---|---|---|---|
| 61 | 4.36 | 6 | -26 | 0 | R | Right Thalamus |
| | 4.35 | 14 | -30 | -4 | R | Right Thalamus |
| | 2.71 | 10 | -34 | 0 | R | Right Thalamus |
| 61 | 3.95 | 14 | -56 | 20 | R | Precuneous Cortex |
| | 2.8 | 22 | -58 | 20 | R | Precuneous Cortex |
| 54 | 3.6 | 26 | -38 | -20 | R | Temporal Fusiform Cortex, posterior division |
| | 3.28 | 30 | -44 | -20 | R | Temporal Occipital Fusiform Cortex |
| | 2.7 | 34 | -40 | -24 | R | Temporal Occipital Fusiform Cortex |
| 53 | -2.94 | 46 | -58 | -6 | R | Inferior Temporal Gyrus, temporooccipital part |
| | -2.86 | 50 | -64 | -10 | R | Lateral Occipital Cortex, inferior division |
| | -2.79 | 46 | -64 | -4 | R | Lateral Occipital Cortex, inferior division |
| 51 | 4.19 | -2 | 36 | -30 | L | Frontal Medial Cortex |
| | 2.81 | 0 | 48 | -26 | C | Frontal Medial Cortex |
| 50 | 3.49 | 14 | -52 | 76 | R | Superior Parietal Lobule |
| | 3.2 | 10 | -56 | 72 | R | Lateral Occipital Cortex, superior division |
| | 2.51 | 8 | -50 | 76 | R | Postcentral Gyrus / Superior Parietal Lobule / Precuneous Cortex |
| 49 | -3.22 | 30 | -74 | 32 | R | Lateral Occipital Cortex, superior division |
| | -2.79 | 30 | -76 | 40 | R | Lateral Occipital Cortex, superior division |
| 49 | -4.34 | -24 | 12 | -30 | L | Temporal Pole |
| | -3.18 | -20 | 8 | -24 | L | Frontal Orbital Cortex |
| | -2.83 | -26 | 4 | -26 | L | Temporal Pole |
| 48 | -2.84 | 38 | -36 | 52 | R | Postcentral Gyrus |
| | -2.27 | 38 | -44 | 56 | R | Superior Parietal Lobule |
| | -2.02 | 42 | -38 | 58 | R | Superior Parietal Lobule |
| 46 | 3.45 | 34 | -72 | -20 | R | Right Crus I |
| | 3.08 | 16 | -82 | -18 | R | Occipital Fusiform Gyrus |
| | 2.96 | 28 | -78 | -18 | R | Occipital Fusiform Gyrus |
| 43 | 3.02 | 22 | 8 | 2 | R | Right Putamen |
| | 2.8 | 26 | 14 | 2 | R | Right Putamen |
| 39 | 3.47 | 0 | 32 | -18 | C | Frontal Medial Cortex |
| | 2.77 | -4 | 28 | -22 | L | Subcallosal Cortex |
| 39 | 3.3 | -22 | 52 | 40 | L | Frontal Pole |
| | 3.02 | -16 | 50 | 44 | L | Frontal Pole |
| 38 | -3.17 | 42 | -50 | -44 | R | Right Crus II |

| Important Clusters from the PLSDA Model Classifying Excitement (movies) | | | | | | |
|---|---|---|---|---|---|---|
| | -2.95 | 36 | -44 | -46 | R | Right VIIb |
| | -2.85 | 40 | -40 | -44 | R | Right VIIb |
| 38 | 3.42 | 62 | -18 | 14 | R | Central Opercular Cortex |
| | 3.15 | 58 | -28 | 12 | R | Planum Temporale |
| | 2.82 | 68 | -20 | 16 | R | Superior Temporal Gyrus, posterior division |
| 37 | 3.67 | -28 | -84 | -20 | L | Occipital Fusiform Gyrus |
| | 2.9 | -22 | -84 | -20 | L | Occipital Fusiform Gyrus |
| | 2.17 | -30 | -82 | -14 | L | Occipital Fusiform Gyrus |
| 37 | -3.54 | 40 | -14 | 0 | R | Insular Cortex |
| | -3.1 | 46 | -14 | -2 | R | Planum Polare |
| | -2.93 | 46 | -16 | 6 | R | Heschl's Gyrus (includes H1 and H2) |
| 37 | 2.72 | 2 | -74 | -24 | R | Vermis VI |
| | 2.68 | 4 | -80 | -32 | R | Right Crus II |
| | 2.33 | 2 | -68 | -28 | R | Vermis VI |
| 35 | 2.9 | 6 | -72 | -4 | R | Lingual Gyrus |
| | 2.86 | 12 | -74 | -6 | R | Lingual Gyrus |
| 34 | -3.61 | -12 | -70 | -44 | L | Left VIIb |
| | -2.46 | -20 | -70 | -46 | L | Left VIIb |
| 34 | 3.55 | 36 | -26 | -20 | R | Temporal Fusiform Cortex, posterior division |
| | 2.82 | 34 | -34 | -14 | R | Parahippocampal Gyrus, posterior division |
| | 2.31 | 30 | -24 | -18 | R | Right Hippocampus |
| 33 | -3.31 | 20 | 28 | -22 | R | Frontal Orbital Cortex |
| | -2.99 | 24 | 32 | -16 | R | Frontal Orbital Cortex |
| 32 | 3.26 | -54 | -50 | 10 | L | Middle Temporal Gyrus, temporooccipital part |
| | 2.31 | -48 | -46 | 8 | L | Middle Temporal Gyrus, temporooccipital part |
| 32 | 4.28 | 18 | -22 | -22 | R | Parahippocampal Gyrus, anterior division |
| | 4.2 | 16 | -32 | -18 | R | Parahippocampal Gyrus, posterior division |
| | 2.81 | 20 | -30 | -22 | R | Parahippocampal Gyrus, posterior division |
| 31 | -3.73 | -40 | 10 | -14 | L | Insular Cortex |
| | -3.22 | -38 | 6 | -18 | L | Temporal Pole |
| 30 | 2.96 | -62 | -30 | 16 | L | Planum Temporale |
| | 2.82 | -64 | -24 | 16 | L | Supramarginal Gyrus, anterior division / Planum Temporale |
| | 2.14 | -58 | -20 | 14 | L | Central Opercular Cortex |

*Fear*

| Important Clusters from the PLSDA Model Classifying Fear (movies) | | | | | | |
|---|---|---|---|---|---|---|
| Cluster Size (Voxels) | Max Coef | MNI | | | Hemisphere | Anatomical Label |
| | | x | y | z | | |
| 470 | 6.7 | -10 | -84 | 4 | L | Intracalcarine Cortex |
| | 5.08 | -10 | -74 | -4 | L | Lingual Gyrus |
| | 4.41 | -6 | -74 | 8 | L | Intracalcarine Cortex |
| 430 | -3.8 | -4 | -92 | -12 | L | Occipital Pole |
| | -3.8 | -10 | -98 | -12 | L | Occipital Pole |
| | -3.61 | -14 | -92 | -14 | L | Occipital Pole |
| 244 | -5.34 | 24 | -90 | -18 | R | Occipital Fusiform Gyrus |
| | -3.94 | 14 | -94 | -6 | R | Occipital Pole |
| | -3.88 | 24 | -78 | -18 | R | Occipital Fusiform Gyrus |
| 241 | 4.03 | 10 | -42 | 56 | R | Precuneous Cortex |
| | 3.87 | 12 | -22 | 44 | R | Precentral Gyrus / Cingulate Gyrus, posterior division |
| | 3.28 | 4 | -44 | 58 | R | Precuneous Cortex |
| 172 | 5.01 | -16 | -48 | -50 | L | Left VIIIb |
| | 4.74 | -30 | -40 | -44 | L | Left VIIIa |
| | 4.58 | -12 | -42 | -52 | L | Left VIIIb |
| 151 | 4.95 | -10 | -20 | 44 | L | Precentral Gyrus |
| | 3.84 | -2 | -22 | 46 | L | Cingulate Gyrus, posterior division |
| | 2.8 | -2 | -28 | 44 | L | Cingulate Gyrus, posterior division |
| 145 | 4.71 | -2 | -60 | -36 | L | Vermis VIIIb |
| | 4.48 | 4 | -60 | -36 | R | Vermis VIIIb |
| | 3.97 | 0 | -62 | -48 | C | Vermis IX |
| 141 | 4.57 | 22 | -36 | -16 | R | Parahippocampal Gyrus, posterior division |
| | 4.12 | 26 | -48 | -8 | R | Lingual Gyrus |
| | 3.78 | 26 | -42 | -10 | R | Lingual Gyrus |
| 140 | -3.58 | -54 | -76 | 2 | L | Lateral Occipital Cortex, inferior division |
| | -3.5 | -52 | -80 | 8 | L | Lateral Occipital Cortex, inferior division |
| | -3.45 | -54 | -68 | 12 | L | Lateral Occipital Cortex, inferior division |
| 104 | -5.04 | 12 | -52 | -2 | R | Lingual Gyrus |
| | -4.01 | 14 | -56 | -6 | R | Lingual Gyrus |
| | -2.86 | 14 | -56 | -12 | R | Right V |
| 60 | 3.6 | -44 | 2 | -16 | L | Planum Polare |
| | 3.56 | -40 | 10 | -14 | L | Insular Cortex |

| Important Clusters from the PLSDA Model Classifying Fear (movies) | | | | | | |
|---|---|---|---|---|---|---|
| | 2.8 | -38 | 2 | -10 | L | Insular Cortex |
| 55 | 3.68 | 0 | -20 | 6 | C | Left Thalamus |
| | 3.38 | 0 | -18 | 0 | C | Left Thalamus |
| | 3.36 | 4 | -26 | 8 | R | Right Thalamus |
| 53 | -4.02 | 8 | -82 | 46 | R | Cuneal Cortex |
| | -3.46 | 10 | -86 | 42 | R | Lateral Occipital Cortex, superior division |
| 52 | -3.2 | 2 | -82 | 36 | R | Cuneal Cortex |
| | -2.7 | -6 | -74 | 28 | L | Precuneous Cortex |
| | -2.45 | 0 | -76 | 28 | C | Cuneal Cortex |
| 51 | -3.91 | 20 | -62 | -6 | R | Lingual Gyrus |
| | -3.39 | 24 | -72 | -6 | R | Occipital Fusiform Gyrus |
| 51 | -4.83 | 0 | 30 | 12 | C | Cingulate Gyrus, anterior division |
| | -2.75 | 2 | 36 | 12 | R | Cingulate Gyrus, anterior division |
| | -2.72 | 2 | 30 | 20 | R | Cingulate Gyrus, anterior division |
| 49 | 3.25 | 16 | -52 | -52 | R | Right VIIIb |
| | 2.82 | 10 | -54 | -58 | R | Right IX |
| | 2.79 | 20 | -46 | -52 | R | Right VIIIb |
| 45 | 4.26 | 44 | -62 | 4 | R | Lateral Occipital Cortex, inferior division |
| | 2.56 | 44 | -68 | -4 | R | Lateral Occipital Cortex, inferior division |
| | 2.26 | 44 | -62 | -2 | R | Lateral Occipital Cortex, inferior division |
| 45 | 5.13 | -32 | 10 | -24 | L | Temporal Pole |
| 43 | 3.34 | -18 | -88 | 24 | L | Lateral Occipital Cortex, superior division |
| | 2.54 | -22 | -84 | 20 | L | Lateral Occipital Cortex, superior division |
| 43 | 3.53 | -12 | -46 | 56 | L | Precuneous Cortex |
| | 3.1 | -14 | -44 | 50 | L | Precuneous Cortex |
| | 2.51 | -16 | -38 | 44 | L | Precentral Gyrus / Precuneous Cortex |
| 42 | -3.43 | -20 | -56 | 2 | L | Lingual Gyrus |
| | -2.27 | -24 | -60 | 6 | L | Precuneous Cortex |
| 42 | 3.38 | -8 | -56 | 74 | L | Superior Parietal Lobule |
| | 3.35 | -8 | -62 | 72 | L | Lateral Occipital Cortex, superior division |
| | 2.37 | -14 | -58 | 74 | L | Lateral Occipital Cortex, superior division |
| 38 | -4.27 | 2 | -72 | 18 | R | Supracalcarine Cortex |
| 37 | 3.39 | 36 | -30 | 20 | R | Parietal Operculum Cortex |
| | 2.31 | 50 | -28 | 18 | R | Parietal Operculum Cortex |
| | 2.23 | 46 | -30 | 22 | R | Parietal Operculum Cortex |

| Important Clusters from the PLSDA Model Classifying Fear (movies) | | | | | | |
|---|---|---|---|---|---|---|
| 37 | 3.09 | 8 | -56 | 68 | R | Precuneous Cortex |
| 37 | -2.82 | 68 | -40 | 6 | R | Middle Temporal Gyrus, temporooccipital part |
| | -2.77 | 62 | -38 | 8 | R | Supramarginal Gyrus, posterior division |
| | -2.36 | 66 | -44 | 10 | R | Middle Temporal Gyrus, temporooccipital part |
| 35 | -2.91 | -52 | -40 | 22 | L | Parietal Operculum Cortex |
| | -2.33 | -58 | -36 | 20 | L | Parietal Operculum Cortex |
| | -2.27 | -62 | -30 | 22 | L | Supramarginal Gyrus, anterior division |
| 34 | 3.79 | -6 | -14 | 10 | L | Left Thalamus |
| | 2.79 | 0 | -12 | -2 | C | Left Thalamus |
| | 2.57 | -4 | -12 | 4 | L | Left Thalamus |
| 32 | -4.34 | -40 | -14 | -8 | L | Planum Polare |
| | -2.3 | -36 | -10 | -2 | L | Insular Cortex |
| 30 | 3.01 | 2 | 62 | 2 | R | Frontal Pole |
| | 2.28 | 4 | 58 | -2 | R | Frontal Pole |
| | 2.28 | 2 | 68 | 2 | R | Frontal Pole |
| 30 | 3.25 | -14 | -80 | 42 | L | Lateral Occipital Cortex, superior division |
| | 2.43 | -14 | -76 | 34 | L | Precuneous Cortex |

*Horror*

| Important Clusters from the PLSDA Model Classifying Horror (movies) | | | | | | |
|---|---|---|---|---|---|---|
| Cluster Size (Voxels) | Max Coef | MNI | | | Hemisphere | Anatomical Label |
| | | x | y | z | | |
| 1002 | -8.75 | 10 | -86 | 6 | R | Intracalcarine Cortex |
| | -7.41 | -8 | -84 | 2 | L | Intracalcarine Cortex |
| | -5.23 | -6 | -90 | 0 | L | Intracalcarine Cortex |
| 187 | 5.5 | 16 | -42 | -52 | R | Right VIIIb |
| | 4.55 | 24 | -38 | -48 | R | Right X |
| | 3.35 | 16 | -36 | -46 | R | Right X |
| 121 | -3.3 | 28 | -32 | -18 | R | Parahippocampal Gyrus, posterior division |
| | -3.25 | 24 | -42 | -18 | R | Temporal Occipital Fusiform Cortex |
| | -3.18 | 26 | -44 | -10 | R | Lingual Gyrus |
| 89 | 3.33 | 4 | -58 | 70 | R | Precuneous Cortex |
| | 3.07 | 2 | -52 | 68 | R | Precuneous Cortex |
| | 3.02 | 0 | -54 | 56 | C | Precuneous Cortex |
| 80 | 4.97 | 14 | -36 | -14 | R | Parahippocampal Gyrus, posterior division |
| | 3.98 | 8 | -40 | 0 | R | Cingulate Gyrus, posterior division |
| | 3.73 | 12 | -46 | -2 | R | Lingual Gyrus |
| 80 | 3.09 | 2 | -78 | 14 | R | Supracalcarine Cortex |
| | 3.02 | 2 | -76 | 22 | R | Cuneal Cortex |
| | 2.53 | 0 | -72 | 14 | C | Supracalcarine Cortex |
| 70 | 3.64 | -12 | -44 | -8 | L | Lingual Gyrus |
| | 3.41 | -18 | -56 | 4 | L | Precuneous Cortex |
| | 3.33 | -18 | -46 | -2 | L | Lingual Gyrus |
| 67 | 3.81 | -12 | -86 | 46 | L | Lateral Occipital Cortex, superior division |
| | 3.48 | -6 | -86 | 42 | L | Lateral Occipital Cortex, superior division |
| | 3.46 | -4 | -82 | 50 | L | Lateral Occipital Cortex, superior division |
| 67 | 4.61 | 12 | -58 | -6 | R | Lingual Gyrus |
| | 3.12 | 6 | -60 | 4 | R | Lingual Gyrus |
| | 2.91 | 20 | -60 | -8 | R | Lingual Gyrus |
| 66 | 3.38 | -52 | 10 | 18 | L | Inferior Frontal Gyrus, pars opercularis |
| | 2.56 | -58 | 8 | 18 | L | Precentral Gyrus |
| | 2.47 | -60 | 12 | 12 | L | Inferior Frontal Gyrus, pars opercularis |
| 64 | -3.65 | 52 | -70 | -18 | R | Lateral Occipital Cortex, inferior division |
| | -3.41 | 46 | -64 | -20 | R | Occipital Fusiform Gyrus |

| Important Clusters from the PLSDA Model Classifying Horror (movies) | | | | | | |
|---|---|---|---|---|---|---|
| | -3.18 | 40 | -56 | -20 | R | Temporal Occipital Fusiform Cortex |
| 60 | 4.59 | -14 | -44 | -58 | L | Left VIIIb |
| | 3.41 | -18 | -42 | -52 | L | Left VIIIb |
| | 2.65 | -12 | -42 | -52 | L | Left VIIIb |
| 59 | -3.4 | 42 | -10 | -2 | R | Insular Cortex |
| | -3.39 | 40 | -8 | -8 | R | Insular Cortex |
| | -2.72 | 40 | -4 | 0 | R | Insular Cortex |
| 59 | 5.31 | -28 | -86 | -20 | L | Occipital Fusiform Gyrus |
| | 3.24 | -20 | -88 | -20 | L | Occipital Fusiform Gyrus |
| | 2.56 | -34 | -90 | -20 | L | Lateral Occipital Cortex, inferior division |
| 59 | 3.26 | 34 | -74 | -20 | R | Right Crus I |
| | 3.11 | 22 | -82 | -16 | R | Occipital Fusiform Gyrus |
| | 2.02 | 24 | -86 | -12 | R | Occipital Fusiform Gyrus |
| 57 | 3.7 | 22 | -56 | 2 | R | Lingual Gyrus |
| | 3.32 | 18 | -60 | 6 | R | Intracalcarine Cortex |
| | 2.95 | 18 | -52 | 0 | R | Lingual Gyrus |
| 56 | 2.88 | 54 | -72 | 2 | R | Lateral Occipital Cortex, inferior division |
| | 2.81 | 48 | -80 | -2 | R | Lateral Occipital Cortex, inferior division |
| | 2.61 | 50 | -74 | -2 | R | Lateral Occipital Cortex, inferior division |
| 53 | -3.54 | 60 | -38 | 2 | R | Middle Temporal Gyrus, temporooccipital part |
| | -3.31 | 50 | -38 | 4 | R | Middle Temporal Gyrus, temporooccipital part |
| 46 | 3.13 | 2 | 50 | 36 | R | Superior Frontal Gyrus |
| 46 | 3.36 | 0 | -66 | -48 | C | Vermis VIIIb |
| | 2.77 | -2 | -72 | -42 | L | Vermis VIIIa |
| | 2.52 | 6 | -70 | -48 | R | Right VIIIa |
| 44 | 2.98 | -66 | -26 | 20 | L | Supramarginal Gyrus, anterior division |
| | 2.74 | -62 | -40 | 20 | L | Supramarginal Gyrus, posterior division |
| | 2.17 | -62 | -34 | 22 | L | Parietal Operculum Cortex |
| 44 | -3.26 | -44 | 26 | -22 | L | Temporal Pole |
| 44 | -3.93 | 54 | 28 | -4 | R | Inferior Frontal Gyrus, pars triangularis |
| 43 | 3.32 | 0 | 34 | -24 | C | Frontal Medial Cortex |
| | 3.01 | 0 | 40 | -22 | C | Frontal Medial Cortex |
| 42 | -3.27 | -36 | -22 | 20 | L | Central Opercular Cortex |
| | -2.77 | -36 | -30 | 16 | L | Planum Temporale |
| 40 | -4.67 | -34 | 12 | -20 | L | Frontal Orbital Cortex |

| Important Clusters from the PLSDA Model Classifying Horror (movies) | | | | | | |
|---|---|---|---|---|---|---|
| | -2.81 | -32 | 10 | -28 | L | Temporal Pole |
| 38 | 3.44 | -4 | -8 | 4 | L | Left Thalamus |
| | 2.93 | -4 | 0 | 2 | L | Left Thalamus |
| | 2.67 | 2 | -6 | 4 | R | Right Thalamus |
| 35 | 3.49 | -4 | -26 | -12 | L | Brain-Stem |
| | 3.09 | -8 | -26 | -4 | L | Left Thalamus |
| | 2.71 | -12 | -26 | -10 | L | Brain-Stem |
| 35 | -3.19 | -50 | 24 | -10 | L | Frontal Orbital Cortex |
| | -2.31 | -48 | 20 | -6 | L | Frontal Orbital Cortex |
| 34 | 2.8 | 42 | -66 | 16 | R | Lateral Occipital Cortex, superior division |
| | 2.77 | 50 | -70 | 18 | R | Lateral Occipital Cortex, superior division |
| 34 | 3.49 | 24 | -26 | -10 | R | Right Hippocampus |
| | 2.82 | 30 | -28 | -10 | R | Right Hippocampus |
| 33 | 3.89 | 0 | -42 | -16 | C | Left I-IV |
| | 2.72 | -4 | -44 | -20 | L | Left I-IV |
| | 2.68 | -4 | -38 | -24 | L | Brain-Stem |
| 33 | -2.99 | 40 | -86 | -16 | R | Lateral Occipital Cortex, inferior division |
| | -2.7 | 46 | -84 | -14 | R | Lateral Occipital Cortex, inferior division |
| 32 | -4.22 | -8 | -36 | 2 | L | Left Thalamus |
| | -3.24 | -8 | -30 | 0 | L | Left Thalamus |
| | -2.6 | -14 | -36 | 2 | L | Left Thalamus |
| 31 | -4.69 | -16 | -2 | -26 | L | Parahippocampal Gyrus, anterior division |
| 30 | -3.47 | 46 | -54 | -8 | R | Inferior Temporal Gyrus, temporooccipital part |
| | -2.49 | 48 | -62 | -10 | R | Lateral Occipital Cortex, inferior division |

*Joy*

| Important Clusters from the PLSDA Model Classifying Joy (movies) | | | | | | |
|---|---|---|---|---|---|---|
| Cluster Size (Voxels) | Max Coef | MNI | | | Hemisphere | Anatomical Label |
| | | x | y | z | | |
| 585 | -5.69 | 28 | -48 | -8 | R | Temporal Occipital Fusiform Cortex |
| | -5.06 | 28 | -56 | -6 | R | Lingual Gyrus |
| | -4.75 | 20 | -36 | -14 | R | Parahippocampal Gyrus, posterior division |
| 534 | 5.57 | -8 | -84 | 2 | L | Intracalcarine Cortex |
| | 4.65 | -14 | -74 | -14 | L | Lingual Gyrus |
| | 4.15 | -14 | -62 | -10 | L | Lingual Gyrus |
| 340 | -4.52 | -22 | -46 | -8 | L | Lingual Gyrus |
| | -4.43 | -22 | -44 | -16 | L | Temporal Occipital Fusiform Cortex |
| | -4.34 | -30 | -48 | -6 | L | Lingual Gyrus |
| 201 | -3.39 | 38 | -80 | 14 | R | Lateral Occipital Cortex, superior division |
| | -3.23 | 38 | -80 | 4 | R | Lateral Occipital Cortex, inferior division |
| | -3.18 | 32 | -76 | 20 | R | Lateral Occipital Cortex, superior division |
| 188 | 4.4 | -52 | -76 | 8 | L | Lateral Occipital Cortex, inferior division |
| | 3.36 | -44 | -78 | 6 | L | Lateral Occipital Cortex, inferior division |
| | 3.08 | -50 | -82 | 4 | L | Lateral Occipital Cortex, inferior division |
| 175 | -4.34 | -24 | -70 | 62 | L | Lateral Occipital Cortex, superior division |
| | -3.47 | -10 | -58 | 74 | L | Lateral Occipital Cortex, superior division |
| | -3.41 | -18 | -64 | 66 | L | Lateral Occipital Cortex, superior division |
| 156 | -4.26 | -32 | -92 | 22 | L | Occipital Pole |
| | -3.12 | -38 | -82 | 16 | L | Lateral Occipital Cortex, superior division |
| | -2.67 | -32 | -92 | 12 | L | Occipital Pole |
| 155 | 4.44 | 56 | -36 | 4 | R | Superior Temporal Gyrus, posterior division |
| | 3.65 | 48 | -36 | 4 | R | Superior Temporal Gyrus, posterior division |
| | 2.84 | 66 | -46 | 6 | R | Middle Temporal Gyrus, temporooccipital part |
| 121 | -4.23 | 0 | -28 | 28 | C | Cingulate Gyrus, posterior division |
| | -4.08 | 0 | -18 | 30 | C | Cingulate Gyrus, posterior division |
| | -3.64 | 0 | -36 | 26 | C | Cingulate Gyrus, posterior division |
| 121 | 4.32 | 52 | 12 | -22 | R | Temporal Pole |
| | 3.21 | 60 | 6 | -10 | R | Superior Temporal Gyrus, anterior division |
| | 2.21 | 56 | 14 | -14 | R | Temporal Pole |
| 92 | 3.92 | 52 | -58 | -20 | R | Inferior Temporal Gyrus, temporooccipital part |
| | 3.08 | 40 | -60 | -14 | R | Temporal Occipital Fusiform Cortex |

| Important Clusters from the PLSDA Model Classifying Joy (movies) | | | | | | |
|---|---|---|---|---|---|---|
| | 3.08 | 48 | -52 | -12 | R | Inferior Temporal Gyrus, temporooccipital part |
| 89 | 3.56 | 40 | -52 | 64 | R | Superior Parietal Lobule |
| | 3.38 | 34 | -58 | 66 | R | Lateral Occipital Cortex, superior division |
| | 2.2 | 38 | -54 | 52 | R | Angular Gyrus |
| 88 | -3.8 | 18 | -44 | -52 | R | Right VIIIb |
| | -3.42 | 24 | -40 | -50 | R | Right VIIIb |
| | -3.36 | 14 | -48 | -54 | R | Right VIIIb |
| 75 | 5.59 | 2 | -50 | -46 | R | Right IX |
| | 5.34 | 2 | -46 | -38 | R | Vermis X |
| | 3.64 | 4 | -52 | -54 | R | Right IX |
| 69 | -3.19 | 30 | -76 | 52 | R | Lateral Occipital Cortex, superior division |
| | -2.93 | 22 | -78 | 50 | R | Lateral Occipital Cortex, superior division |
| | -2.74 | 16 | -80 | 48 | R | Lateral Occipital Cortex, superior division |
| 68 | -3.83 | 10 | -98 | 14 | R | Occipital Pole |
| | -2.15 | 6 | -94 | 10 | R | Occipital Pole |
| 62 | 3.38 | 58 | -30 | 28 | R | Parietal Operculum Cortex |
| | 3 | 64 | -24 | 38 | R | Supramarginal Gyrus, anterior division |
| | 2.35 | 60 | -30 | 36 | R | Supramarginal Gyrus, anterior division |
| 62 | 3.13 | 56 | -60 | 10 | R | Middle Temporal Gyrus, temporooccipital part |
| | 2.88 | 54 | -54 | 0 | R | Middle Temporal Gyrus, temporooccipital part |
| | 2.59 | 60 | -58 | 0 | R | Middle Temporal Gyrus, temporooccipital part |
| 61 | 3.46 | 14 | -80 | 8 | R | Intracalcarine Cortex |
| | 2.57 | 10 | -84 | 10 | R | Intracalcarine Cortex |
| | 2.56 | 12 | -74 | 10 | R | Intracalcarine Cortex |
| 59 | 3.3 | 46 | -42 | 60 | R | Superior Parietal Lobule |
| | 2.96 | 50 | -36 | 60 | R | Supramarginal Gyrus, posterior division |
| | 2.47 | 56 | -30 | 56 | R | Supramarginal Gyrus, anterior division |
| 56 | 5 | -22 | 0 | -20 | L | Left Amygdala |
| | 3.13 | -20 | 6 | -20 | L | Frontal Orbital Cortex |
| | 2.94 | -24 | -6 | -22 | L | Left Amygdala |
| 53 | 3.31 | 36 | 18 | -38 | R | Temporal Pole |
| | 2.81 | 44 | 4 | -44 | R | Inferior Temporal Gyrus, anterior division |
| | 2.5 | 38 | 6 | -44 | R | Temporal Pole |
| 52 | -3.16 | -58 | -18 | 30 | L | Postcentral Gyrus |
| | -2.57 | -58 | -18 | 40 | L | Postcentral Gyrus |

| Important Clusters from the PLSDA Model Classifying Joy (movies) | | | | | | |
|---|---|---|---|---|---|---|
| 52 | -3.84 | 16 | -50 | 6 | R | Precuneous Cortex |
| | -2.94 | 8 | -48 | 4 | R | Cingulate Gyrus, posterior division |
| 50 | -3.4 | 16 | -54 | 16 | R | Precuneous Cortex |
| | -3 | 22 | -58 | 16 | R | Precuneous Cortex |
| 48 | 3.45 | 64 | -50 | 14 | R | Angular Gyrus |
| | 2.79 | 58 | -50 | 12 | R | Middle Temporal Gyrus, temporooccipital part |
| | 2.58 | 64 | -58 | 14 | R | Angular Gyrus |
| 48 | -4.02 | -4 | -102 | 10 | L | Occipital Pole |
| | -2.93 | -2 | -102 | 0 | L | Occipital Pole |
| 43 | 3.82 | -44 | -56 | -22 | L | Temporal Occipital Fusiform Cortex |
| | 2.9 | -44 | -56 | -16 | L | Temporal Occipital Fusiform Cortex |
| | 2.47 | -42 | -52 | -10 | L | Inferior Temporal Gyrus, temporooccipital part |
| 42 | -3.6 | -48 | 18 | -10 | L | Frontal Orbital Cortex |
| | -2.19 | -54 | 22 | -6 | L | Inferior Frontal Gyrus, pars triangularis |
| 41 | 3.72 | -24 | -36 | -44 | L | Left X |
| | 2.66 | -16 | -38 | -44 | L | Left X |
| 36 | 2.89 | 2 | -86 | 20 | R | Cuneal Cortex |
| | 2.64 | 6 | -80 | 20 | R | Cuneal Cortex |
| 36 | -4.19 | 8 | -98 | -6 | R | Occipital Pole |
| | -3.4 | 10 | -96 | -12 | R | Occipital Pole |
| | -2.13 | 4 | -94 | -10 | R | Occipital Pole |
| 35 | -3.88 | 34 | 20 | -22 | R | Frontal Orbital Cortex |
| | -2.56 | 34 | 22 | -16 | R | Frontal Orbital Cortex |
| 34 | 5.12 | 2 | -4 | 2 | R | Right Thalamus |
| | 2.16 | 4 | -4 | 12 | R | Right Thalamus |
| 33 | 4.15 | 2 | 36 | 0 | R | Cingulate Gyrus, anterior division |
| | 2.92 | -2 | 42 | -2 | L | Cingulate Gyrus, anterior division |
| 32 | 3.49 | 46 | -84 | -6 | R | Lateral Occipital Cortex, inferior division |
| | 3.11 | 42 | -88 | -8 | R | Lateral Occipital Cortex, inferior division |
| 32 | 3.44 | 42 | -74 | -18 | R | Lateral Occipital Cortex, inferior division |
| | 2.41 | 44 | -70 | -12 | R | Lateral Occipital Cortex, inferior division |
| 31 | -2.81 | -56 | 12 | 16 | L | Inferior Frontal Gyrus, pars opercularis |
| | -2.58 | -48 | 6 | 20 | L | Precentral Gyrus |
| 30 | 4.46 | 46 | -4 | -2 | R | Insular Cortex |
| | 2.92 | 50 | 0 | 0 | R | Central Opercular Cortex |

*Neutral*

| Important Clusters from the PLSDA Model Classifying Neutral (movies) | | | | | | |
|---|---|---|---|---|---|---|
| Cluster Size (Voxels) | Max Coef | MNI | | | Hemisphere | Anatomical Label |
| | | x | y | z | | |
| 493 | -3.55 | -50 | -80 | 6 | L | Lateral Occipital Cortex, inferior division |
| | -3.31 | -52 | -74 | 10 | L | Lateral Occipital Cortex, inferior division |
| | -3.16 | -56 | -68 | 12 | L | Lateral Occipital Cortex, inferior division |
| 486 | -4.4 | -2 | -92 | -2 | L | Occipital Pole |
| | -4.27 | -20 | -84 | -18 | L | Occipital Fusiform Gyrus |
| | -3.08 | 14 | -94 | 0 | R | Occipital Pole |
| 332 | -3.4 | 54 | -62 | 12 | R | Lateral Occipital Cortex, inferior division |
| | -3.27 | 44 | -60 | 10 | R | Lateral Occipital Cortex, inferior division |
| | -3.23 | 52 | -58 | 4 | R | Middle Temporal Gyrus, temporooccipital part |
| 144 | -3.46 | -10 | -18 | 10 | L | Left Thalamus |
| | -2.95 | 0 | -14 | 10 | C | Left Thalamus |
| | -2.93 | 4 | -18 | 8 | R | Right Thalamus |
| 138 | -4.08 | 22 | -54 | 4 | R | Lingual Gyrus |
| | -3.76 | 12 | -40 | -4 | R | Lingual Gyrus |
| | -2.94 | 18 | -50 | 0 | R | Lingual Gyrus |
| 129 | -3.19 | -24 | -70 | -6 | L | Occipital Fusiform Gyrus |
| | -3.01 | -20 | -64 | -12 | L | Lingual Gyrus |
| | -2.5 | -24 | -72 | -16 | L | Occipital Fusiform Gyrus |
| 119 | -4.04 | -2 | -32 | -2 | L | Brain-Stem |
| | -3.98 | 0 | -26 | -2 | C | Brain-Stem |
| | -3.86 | 14 | -26 | -10 | R | Brain-Stem |
| 105 | 4.8 | 0 | -32 | 26 | C | Cingulate Gyrus, posterior division |
| | 4.25 | 0 | -38 | 24 | C | Cingulate Gyrus, posterior division |
| | 3.88 | 0 | -24 | 28 | C | Cingulate Gyrus, posterior division |
| 89 | 2.88 | 28 | 66 | -4 | R | Frontal Pole |
| | 2.47 | 38 | 62 | 0 | R | Frontal Pole |
| | 2.24 | 22 | 68 | -2 | R | Frontal Pole |
| 87 | -3.31 | 8 | -54 | 74 | R | Precuneous Cortex |
| | -3.15 | 6 | -58 | 70 | R | Precuneous Cortex |
| | -2.69 | 16 | -62 | 70 | R | Lateral Occipital Cortex, superior division |
| 77 | -3.57 | -16 | -58 | 4 | L | Precuneous Cortex |
| | -2.43 | -16 | -50 | -2 | L | Lingual Gyrus |

| Important Clusters from the PLSDA Model Classifying Neutral (movies) | | | | | | |
|---|---|---|---|---|---|---|
| 77 | -4.24 | 20 | -42 | -48 | R | Right VIIIb |
| | -3.54 | 26 | -38 | -42 | R | Right X |
| | -2.89 | 16 | -54 | -50 | R | Right VIIIb |
| 76 | -3.31 | 50 | 8 | 38 | R | Precentral Gyrus |
| | -2.79 | 56 | 12 | 40 | R | Middle Frontal Gyrus |
| | -2.41 | 44 | 12 | 30 | R | Middle Frontal Gyrus |
| 75 | -3.53 | 30 | -76 | -18 | R | Occipital Fusiform Gyrus |
| | -2.35 | 38 | -64 | -18 | R | Occipital Fusiform Gyrus |
| 71 | -2.49 | 2 | -58 | 44 | R | Precuneous Cortex |
| | -2.47 | 2 | -58 | 60 | R | Precuneous Cortex |
| | -2.33 | 0 | -60 | 52 | C | Precuneous Cortex |
| 65 | -3.43 | 44 | 0 | 60 | R | Middle Frontal Gyrus |
| | -2.96 | 50 | 2 | 54 | R | Precentral Gyrus |
| | -2.71 | 40 | -2 | 64 | R | Precentral Gyrus |
| 53 | 3.19 | 60 | -2 | 6 | R | Central Opercular Cortex |
| | 2.67 | 66 | -4 | 4 | R | Superior Temporal Gyrus, anterior division |
| 51 | -3.21 | -16 | -44 | -46 | L | Left X |
| | -2.58 | -24 | -42 | -48 | L | Left VIIIb |
| | -2.52 | -22 | -36 | -44 | L | Left X |
| 49 | -3.62 | 18 | -4 | -14 | R | Right Amygdala |
| | -2.92 | 22 | 2 | -14 | R | Right Cerebral Cortex |
| 47 | 3.02 | 46 | -60 | 54 | R | Lateral Occipital Cortex, superior division |
| | 2.73 | 50 | -62 | 48 | R | Lateral Occipital Cortex, superior division |
| 45 | -2.66 | -44 | -52 | -18 | L | Inferior Temporal Gyrus, temporooccipital part |
| | -2.5 | -42 | -52 | -10 | L | Inferior Temporal Gyrus, temporooccipital part |
| | -2.27 | -40 | -50 | -24 | L | Temporal Occipital Fusiform Cortex |
| 44 | -3.05 | 56 | -52 | 14 | R | Angular Gyrus |
| | -2.36 | 64 | -52 | 16 | R | Angular Gyrus |
| 42 | -2.97 | -8 | -46 | 58 | L | Precuneous Cortex |
| | -2.73 | -12 | -44 | 52 | L | Precuneous Cortex |
| | -2.62 | -14 | -40 | 46 | L | Precuneous Cortex |
| 39 | -3.53 | -20 | -2 | -14 | L | Left Amygdala |
| 38 | -3.14 | 2 | -52 | 20 | R | Cingulate Gyrus, posterior division |
| | -2.24 | -4 | -54 | 28 | L | Cingulate Gyrus, posterior division |
| 36 | -2.88 | 0 | -54 | -34 | C | Vermis IX |

| Important Clusters from the PLSDA Model Classifying Neutral (movies) | | | | | | |
|---|---|---|---|---|---|---|
| | -2.6 | 0 | -54 | -40 | C | Vermis IX |
| 34 | 2.93 | 46 | -4 | -6 | R | Planum Polare |
| | 2.51 | 44 | 4 | -12 | R | Planum Polare |
| | 2.37 | 48 | -8 | 0 | R | Heschl's Gyrus (includes H1 and H2) |
| 33 | 3.59 | 12 | -66 | 38 | R | Precuneous Cortex |
| 33 | -3.2 | -10 | -62 | 70 | L | Lateral Occipital Cortex, superior division |
| | -2.25 | -8 | -54 | 74 | L | Superior Parietal Lobule |
| 32 | 2.7 | -8 | 6 | -20 | L | Subcallosal Cortex |
| | 2.61 | -12 | 12 | -24 | L | Frontal Orbital Cortex |
| | 2.19 | -16 | 12 | -18 | L | Frontal Orbital Cortex |
| 31 | 2.94 | -40 | -66 | 56 | L | Lateral Occipital Cortex, superior division |
| | 2.7 | -42 | -70 | 50 | L | Lateral Occipital Cortex, superior division |
| | 2.22 | -46 | -64 | 52 | L | Lateral Occipital Cortex, superior division |
| 31 | 2.95 | -10 | -68 | 38 | L | Precuneous Cortex |
| | 2.57 | -8 | -70 | 44 | L | Precuneous Cortex |
| | 2.54 | -6 | -72 | 36 | L | Precuneous Cortex |

*Romance*

| Important Clusters from the PLSDA Model Classifying Romance (movies) | | | | | | |
|---|---|---|---|---|---|---|
| **Cluster Size (Voxels)** | **Max Coef** | **MNI** | | | **Hemisphere** | **Anatomical Label** |
| | | **x** | **y** | **z** | | |
| 359 | 5.07 | 38 | -56 | -20 | R | Temporal Occipital Fusiform Cortex |
| | 4.67 | 36 | -50 | -22 | R | Temporal Occipital Fusiform Cortex |
| | 4.23 | 42 | -44 | -22 | R | Temporal Occipital Fusiform Cortex |
| 247 | 4.99 | 0 | -26 | 28 | C | Cingulate Gyrus, posterior division |
| | 4.7 | 0 | -36 | 26 | C | Cingulate Gyrus, posterior division |
| | 3.53 | 0 | -38 | 32 | C | Cingulate Gyrus, posterior division |
| 225 | 3.92 | 10 | -94 | 20 | R | Occipital Pole |
| | 3.78 | 8 | -96 | 12 | R | Occipital Pole |
| | 3.56 | 16 | -94 | 16 | R | Occipital Pole |
| 218 | -5.3 | -14 | -76 | -14 | L | Lingual Gyrus |
| | -4.8 | -28 | -86 | -20 | L | Occipital Fusiform Gyrus |
| | -3.42 | -18 | -86 | -20 | L | Occipital Fusiform Gyrus |
| 204 | -3.78 | 26 | -96 | 2 | R | Occipital Pole |
| | -3.48 | 14 | -96 | -8 | R | Occipital Pole |
| | -3.19 | 14 | -102 | 0 | R | Occipital Pole |
| 187 | -4.46 | -8 | -86 | 44 | L | Lateral Occipital Cortex, superior division |
| | -3.65 | 2 | -78 | 20 | R | Cuneal Cortex |
| | -2.94 | -2 | -86 | 40 | L | Cuneal Cortex |
| 168 | -4.24 | -54 | -72 | -6 | L | Lateral Occipital Cortex, inferior division |
| | -3.48 | -46 | -70 | -20 | L | Lateral Occipital Cortex, inferior division |
| | -3.4 | -58 | -66 | -6 | L | Lateral Occipital Cortex, inferior division |
| 167 | 3.91 | 54 | -62 | 12 | R | Lateral Occipital Cortex, inferior division |
| | 3.32 | 54 | -64 | 4 | R | Lateral Occipital Cortex, inferior division |
| | 2.92 | 54 | -74 | 6 | R | Lateral Occipital Cortex, inferior division |
| 144 | -4.56 | 66 | -18 | 14 | R | Planum Temporale |
| | -3.48 | 66 | -2 | 6 | R | Superior Temporal Gyrus, anterior division / Central Opercular Cortex / Planum Polare |
| | -3.32 | 66 | -12 | 8 | R | Planum Temporale |
| 142 | 6.46 | 2 | 6 | -16 | R | Subcallosal Cortex |
| | 5.06 | 2 | 8 | -8 | R | Subcallosal Cortex |
| | 3.9 | -2 | 2 | -12 | L | Left Cerebral Cortex |
| 141 | 4.36 | -42 | -50 | -20 | L | Temporal Occipital Fusiform Cortex |
| | 3.31 | -38 | -44 | -24 | L | Temporal Fusiform Cortex, posterior division |

| Important Clusters from the PLSDA Model Classifying Romance (movies) | | | | | | |
|---|---|---|---|---|---|---|
| | 2.75 | -38 | -36 | -22 | L | Temporal Fusiform Cortex, posterior division |
| 137 | 4.29 | -4 | -100 | 6 | L | Occipital Pole |
| | 3.23 | -4 | -102 | 14 | L | Occipital Pole |
| 134 | 3.67 | 48 | 14 | 30 | R | Inferior Frontal Gyrus, pars opercularis |
| | 3.44 | 38 | 16 | 26 | R | Inferior Frontal Gyrus, pars opercularis |
| | 3.4 | 40 | 22 | 24 | R | Middle Frontal Gyrus |
| 132 | 3.46 | 38 | -56 | 64 | R | Superior Parietal Lobule |
| | 3.28 | 36 | -52 | 58 | R | Superior Parietal Lobule |
| | 3.23 | 28 | -52 | 54 | R | Superior Parietal Lobule |
| 114 | 3.69 | 58 | -30 | 32 | R | Supramarginal Gyrus, anterior division |
| | 2.89 | 60 | -34 | 26 | R | Parietal Operculum Cortex |
| | 2.66 | 66 | -38 | 28 | R | Supramarginal Gyrus, posterior division |
| 86 | 3.7 | 48 | 4 | 38 | R | Precentral Gyrus |
| | 2.51 | 38 | 4 | 34 | R | Precentral Gyrus |
| | 2.44 | 58 | 4 | 42 | R | Precentral Gyrus |
| 85 | -3.41 | -52 | -28 | 14 | L | Parietal Operculum Cortex |
| | -3.23 | -58 | -28 | 14 | L | Parietal Operculum Cortex |
| | -2.76 | -66 | -22 | 14 | L | Superior Temporal Gyrus, posterior division |
| 82 | 4.96 | 2 | -26 | -4 | R | Brain-Stem |
| | 3.86 | 8 | -28 | -4 | R | Brain-Stem |
| | 2.9 | 22 | -30 | -4 | R | Right Thalamus |
| 72 | 3.58 | -28 | -72 | 60 | L | Lateral Occipital Cortex, superior division |
| | 3.13 | -36 | -60 | 60 | L | Lateral Occipital Cortex, superior division |
| | 2.68 | -36 | -68 | 58 | L | Lateral Occipital Cortex, superior division |
| 66 | -3.32 | -18 | -94 | -8 | L | Occipital Pole |
| | -2.7 | -10 | -98 | -14 | L | Occipital Pole |
| 64 | -4.97 | 16 | -40 | -54 | R | Right VIIIb |
| | -3.1 | 16 | -52 | -52 | R | Right VIIIb |
| | -2.49 | 10 | -52 | -56 | R | Right IX |
| 61 | -3.75 | 8 | -64 | -2 | R | Lingual Gyrus |
| | -3.3 | 6 | -60 | 4 | R | Lingual Gyrus |
| | -2.66 | 14 | -66 | -2 | R | Lingual Gyrus |
| 60 | 3.32 | 22 | -74 | -12 | R | Occipital Fusiform Gyrus |
| | 3.07 | 24 | -76 | -6 | R | Occipital Fusiform Gyrus |
| | 2.85 | 28 | -76 | -16 | R | Occipital Fusiform Gyrus |

| Important Clusters from the PLSDA Model Classifying Romance (movies) | | | | | | |
|---|---|---|---|---|---|---|
| 55 | -2.89 | 64 | -16 | 30 | R | Postcentral Gyrus |
| | -2.62 | 58 | -18 | 24 | R | Postcentral Gyrus |
| | -2.27 | 56 | -14 | 30 | R | Postcentral Gyrus |
| 53 | 4.65 | 18 | -10 | -16 | R | Right Amygdala |
| 52 | 3.27 | -6 | 4 | 52 | L | Juxtapositional Lobule Cortex (formerly Supplementary Motor Cortex) |
| | 2.59 | 0 | 0 | 56 | C | Juxtapositional Lobule Cortex (formerly Supplementary Motor Cortex) |
| 52 | 4.08 | 20 | -52 | -14 | R | Lingual Gyrus |
| | 3.12 | 18 | -58 | -14 | R | Right VI |
| | 2.02 | 20 | -64 | -12 | R | Lingual Gyrus |
| 52 | 3.1 | 0 | 8 | 46 | C | Paracingulate Gyrus |
| | 2.28 | 6 | 6 | 46 | R | Juxtapositional Lobule Cortex (formerly Supplementary Motor Cortex) |
| 49 | -3.1 | -16 | -62 | -4 | L | Lingual Gyrus |
| | -3.1 | -10 | -66 | -2 | L | Lingual Gyrus |
| | -2.53 | -18 | -56 | -6 | L | Lingual Gyrus |
| 49 | 4.25 | -12 | -28 | -16 | L | Brain-Stem |
| | 4.19 | -16 | -30 | -20 | L | Parahippocampal Gyrus, posterior division |
| | 3.4 | -14 | -20 | -28 | L | Brain-Stem |
| 46 | 3.46 | 56 | -40 | 12 | R | Supramarginal Gyrus, posterior division |
| | 2.53 | 60 | -44 | 16 | R | Supramarginal Gyrus, posterior division |
| 46 | -3.16 | -10 | -80 | 10 | L | Intracalcarine Cortex |
| | -2.56 | -8 | -86 | 2 | L | Intracalcarine Cortex |
| 43 | 3.2 | -50 | -78 | 6 | L | Lateral Occipital Cortex, inferior division |
| | 3.01 | -54 | -76 | 10 | L | Lateral Occipital Cortex, inferior division |
| | 2.33 | -44 | -74 | 6 | L | Lateral Occipital Cortex, inferior division |
| 42 | -5.08 | -24 | 0 | -20 | L | Left Amygdala |
| | -3.03 | -24 | 0 | -12 | L | Left Cerebral Cortex |
| | -2.73 | -28 | -4 | -8 | L | Left Putamen |
| 42 | 2.74 | -10 | 50 | -4 | L | Paracingulate Gyrus |
| | 2.35 | -6 | 58 | 6 | L | Frontal Pole |
| | 2.33 | -4 | 56 | -2 | L | Frontal Pole |
| 41 | -3.18 | 18 | 10 | -4 | R | Right Putamen |
| | -2.97 | 26 | 6 | -2 | R | Right Putamen |
| | -2.32 | 24 | 12 | -2 | R | Right Putamen |

| Important Clusters from the PLSDA Model Classifying Romance (movies) | | | | | | |
|---|---|---|---|---|---|---|
| 41 | -5.07 | 44 | -14 | 2 | R | Heschl's Gyrus (includes H1 and H2) |
| | -3.09 | 44 | -8 | -2 | R | Insular Cortex |
| 40 | -4.03 | 2 | -48 | 10 | R | Cingulate Gyrus, posterior division |
| 39 | -4.44 | 46 | -78 | -18 | R | Lateral Occipital Cortex, inferior division |
| | -4.11 | 40 | -82 | -20 | R | Lateral Occipital Cortex, inferior division |
| 37 | 4.77 | -16 | 4 | -26 | L | Parahippocampal Gyrus, anterior division |
| | 2.7 | -30 | 6 | -24 | L | Temporal Pole |
| | 2.65 | -22 | 4 | -24 | L | Temporal Pole |
| 36 | -4.18 | -8 | -54 | -4 | L | Lingual Gyrus |
| 35 | 3.03 | -4 | 26 | -20 | L | Subcallosal Cortex |
| | 3.02 | 0 | 28 | -24 | C | Subcallosal Cortex |
| | 2.53 | 4 | 34 | -26 | R | Frontal Medial Cortex |
| 33 | 3.92 | -38 | -70 | -20 | L | Occipital Fusiform Gyrus |
| 33 | 3.5 | -36 | -78 | -12 | L | Occipital Fusiform Gyrus |
| | 2.26 | -32 | -84 | -14 | L | Occipital Fusiform Gyrus |
| 32 | -2.74 | -44 | 6 | -38 | L | Temporal Pole |
| | -2.19 | -38 | 4 | -36 | L | Temporal Pole |
| 31 | -4.14 | 22 | 0 | -16 | R | Right Amygdala |
| 31 | -4.98 | 28 | -10 | -14 | R | Right Amygdala |
| | -2.91 | 28 | -16 | -12 | R | Right Hippocampus |
| 30 | 4.07 | -10 | -38 | -8 | L | Parahippocampal Gyrus, posterior division |
| | 3.75 | -14 | -32 | -12 | L | Parahippocampal Gyrus, posterior division |
| | 2.87 | -14 | -32 | -4 | L | Left Thalamus |
| 30 | -2.83 | 56 | 2 | -30 | R | Middle Temporal Gyrus, anterior division |
| | -2.66 | 50 | -2 | -34 | R | Inferior Temporal Gyrus, anterior division |
| | -2.42 | 54 | 2 | -36 | R | Middle Temporal Gyrus, anterior division |

*Sadness*

| Important Clusters from the PLSDA Model Classifying Sadness (movies) | | | | | | |
|---|---|---|---|---|---|---|
| **Cluster Size (Voxels)** | **Max Coef** | **MNI** | | | **Hemisphere** | **Anatomical Label** |
| | | **x** | **y** | **z** | | |
| 326 | 5.9 | -10 | -76 | -12 | L | Lingual Gyrus |
| | 5.43 | -8 | -70 | -8 | L | Lingual Gyrus |
| | 4.09 | -14 | -76 | -6 | L | Lingual Gyrus |
| 217 | 8.42 | -8 | -80 | 6 | L | Intracalcarine Cortex |
| | 3.46 | -4 | -88 | -2 | L | Intracalcarine Cortex |
| | 3.01 | -4 | -52 | 6 | L | Precuneous Cortex |
| 191 | 3.69 | 54 | -76 | 0 | R | Lateral Occipital Cortex, inferior division |
| | 3.54 | 46 | -68 | 0 | R | Lateral Occipital Cortex, inferior division |
| | 3.12 | 48 | -78 | 0 | R | Lateral Occipital Cortex, inferior division |
| 150 | -5.38 | 28 | -42 | -10 | R | Lingual Gyrus |
| | -5.34 | 32 | -46 | -8 | R | Temporal Occipital Fusiform Cortex |
| | -4.53 | 28 | -58 | -6 | R | Lingual Gyrus |
| 112 | -4.22 | -14 | -38 | -14 | L | Parahippocampal Gyrus, posterior division |
| | -3.95 | -20 | -54 | -2 | L | Lingual Gyrus |
| | -3.61 | -22 | -42 | -14 | L | Parahippocampal Gyrus, posterior division |
| 92 | -2.95 | -24 | -100 | -12 | L | Occipital Pole |
| | -2.9 | -12 | -100 | -8 | L | Occipital Pole |
| | -2.84 | -18 | -98 | -16 | L | Occipital Pole |
| 88 | 3.48 | -2 | -54 | 48 | L | Precuneous Cortex |
| | 3.12 | 4 | -62 | 44 | R | Precuneous Cortex |
| | 2.59 | 0 | -54 | 38 | C | Precuneous Cortex |
| 79 | -3.99 | 14 | -94 | 0 | R | Occipital Pole |
| | -2.28 | 6 | -90 | 2 | R | Occipital Pole |
| 78 | 3.39 | -46 | -10 | 0 | L | Planum Polare |
| | 3.36 | -42 | -16 | 4 | L | Heschl's Gyrus (includes H1 and H2) |
| | 3 | -46 | -18 | 10 | L | Heschl's Gyrus (includes H1 and H2) |
| 69 | -3.34 | -2 | 58 | 6 | L | Frontal Pole |
| | -2.72 | 2 | 58 | -6 | R | Frontal Pole |
| 66 | -3.32 | 20 | -100 | -10 | R | Occipital Pole |
| | -2.94 | 20 | -96 | -16 | R | Occipital Pole |
| | -2.68 | 26 | -92 | -16 | R | Occipital Pole |
| 65 | -4.19 | -24 | 4 | -22 | L | Temporal Pole |

| Important Clusters from the PLSDA Model Classifying Sadness (movies) | | | | | | |
|---|---|---|---|---|---|---|
| | -3.56 | -14 | 8 | -22 | L | Frontal Orbital Cortex |
| | -2.85 | -8 | 6 | -20 | L | Subcallosal Cortex |
| 63 | 3.74 | 2 | -58 | 24 | R | Precuneous Cortex |
| | 2.94 | 6 | -62 | 18 | R | Precuneous Cortex |
| | 2.4 | 0 | -66 | 22 | C | Precuneous Cortex |
| 57 | -3.16 | 28 | -68 | -16 | R | Occipital Fusiform Gyrus |
| | -2.59 | 22 | -72 | -14 | R | Occipital Fusiform Gyrus |
| | -2.52 | 34 | -74 | -18 | R | Occipital Fusiform Gyrus |
| 56 | -3.85 | 58 | -22 | 16 | R | Parietal Operculum Cortex |
| | -2.94 | 56 | -30 | 20 | R | Parietal Operculum Cortex |
| | -2.55 | 52 | -34 | 22 | R | Parietal Operculum Cortex |
| 53 | -3.35 | 60 | -20 | -4 | R | Superior Temporal Gyrus, posterior division |
| | -3.07 | 68 | -22 | -2 | R | Superior Temporal Gyrus, posterior division |
| | -2.63 | 64 | -26 | 0 | R | Superior Temporal Gyrus, posterior division |
| 52 | -3.43 | -52 | -52 | 16 | L | Angular Gyrus |
| | -2.8 | -52 | -50 | 10 | L | Middle Temporal Gyrus, temporooccipital part |
| 51 | 3.2 | 0 | 6 | 40 | C | Cingulate Gyrus, anterior division |
| | 3.13 | 2 | 12 | 40 | R | Cingulate Gyrus, anterior division |
| | 2.91 | 2 | 14 | 34 | R | Cingulate Gyrus, anterior division |
| 50 | -3.86 | 10 | -74 | 8 | R | Intracalcarine Cortex |
| | -2.51 | 14 | -78 | 6 | R | Intracalcarine Cortex |
| 50 | 5.02 | -4 | -4 | 8 | L | Left Thalamus |
| | 4.87 | -8 | 2 | 6 | L | Left Caudate |
| | 2.47 | -8 | -10 | 10 | L | Left Thalamus |
| 49 | -4.75 | 0 | 4 | -10 | C | Left Cerebral Cortex |
| | -3.48 | 0 | 20 | -6 | C | Subcallosal Cortex |
| | -3.29 | 0 | 12 | -10 | C | Subcallosal Cortex |
| 43 | 3.69 | 46 | -8 | -2 | R | Planum Polare |
| | 3.33 | 40 | -6 | 0 | R | Insular Cortex |
| | 3.19 | 46 | -14 | 2 | R | Heschl's Gyrus (includes H1 and H2) |
| 42 | 4.47 | 16 | -6 | -28 | R | Parahippocampal Gyrus, anterior division |
| | 2.87 | 14 | -4 | -22 | R | Parahippocampal Gyrus, anterior division |
| | 2.11 | 18 | 0 | -28 | R | Parahippocampal Gyrus, anterior division |
| 38 | 3.84 | 52 | -12 | 10 | R | Central Opercular Cortex |
| | 2.61 | 60 | -14 | 10 | R | Central Opercular Cortex |

| Important Clusters from the PLSDA Model Classifying Sadness (movies) | | | | | | |
|---|---|---|---|---|---|---|
| 36 | -2.79 | 58 | -44 | 34 | R | Supramarginal Gyrus, posterior division |
| | -2.43 | 58 | -46 | 28 | R | Angular Gyrus |
| | -2.21 | 60 | -38 | 36 | R | Supramarginal Gyrus, posterior division |
| 35 | -3.91 | 38 | 12 | -14 | R | Insular Cortex |
| 34 | -2.96 | -36 | -68 | -18 | L | Occipital Fusiform Gyrus |
| | -2.69 | -38 | -64 | -22 | L | Left Crus I |
| | -2.44 | -30 | -70 | -18 | L | Occipital Fusiform Gyrus |
| 32 | 4.02 | 6 | -48 | 2 | R | Cingulate Gyrus, posterior division |
| | 3.1 | 2 | -52 | 10 | R | Precuneous Cortex |
| | 2.12 | 10 | -50 | 6 | R | Cingulate Gyrus, posterior division |
| 32 | 3.8 | -2 | -46 | -62 | L | Brain-Stem |
| | 3.37 | 4 | -38 | -62 | R | Brain-Stem |
| | 2.8 | -6 | -48 | -58 | L | Brain-Stem |
| 31 | -3.35 | -54 | -38 | 2 | L | Superior Temporal Gyrus, posterior division |
| | -2.23 | -48 | -36 | 0 | L | Superior Temporal Gyrus, posterior division |
| 30 | -3.46 | -48 | -28 | -4 | L | Superior Temporal Gyrus, posterior division |

*Surprise*

| Important Clusters from the PLSDA Model Classifying Surprise (movies) | | | | | | |
|---|---|---|---|---|---|---|
| Cluster Size (Voxels) | Max Coef | MNI | | | Hemisphere | Anatomical Label |
| | | x | y | z | | |
| 274 | -4.08 | 4 | -50 | 20 | R | Cingulate Gyrus, posterior division |
| | -3.51 | -2 | -58 | 28 | L | Precuneous Cortex |
| | -3.48 | 0 | -64 | 26 | C | Precuneous Cortex |
| 190 | 3.75 | -42 | 6 | 28 | L | Precentral Gyrus |
| | 3.71 | -46 | 2 | 38 | L | Precentral Gyrus |
| | 3.61 | -46 | 4 | 22 | L | Precentral Gyrus |
| 152 | -3.41 | 50 | -74 | -8 | R | Lateral Occipital Cortex, inferior division |
| | -3.29 | 52 | -74 | -2 | R | Lateral Occipital Cortex, inferior division |
| | -2.84 | 44 | -84 | -6 | R | Lateral Occipital Cortex, inferior division |
| 127 | -4.33 | 20 | -26 | -12 | R | Parahippocampal Gyrus, posterior division |
| | -3.99 | 22 | -12 | -12 | R | Right Amygdala |
| | -3.98 | 18 | -8 | -14 | R | Right Amygdala |
| 106 | 3.45 | -58 | -34 | 32 | L | Supramarginal Gyrus, anterior division |
| | 3.19 | -64 | -34 | 36 | L | Supramarginal Gyrus, anterior division |
| | 2.79 | -64 | -34 | 42 | L | Supramarginal Gyrus, anterior division |
| 105 | -5.03 | 2 | 6 | -12 | R | Subcallosal Cortex |
| | -3.54 | -6 | 2 | -14 | L | Left Cerebral Cortex |
| | -2.79 | 4 | 14 | -12 | R | Subcallosal Cortex |
| 97 | -4.86 | -14 | -32 | -16 | L | Parahippocampal Gyrus, posterior division |
| | -3.57 | -10 | -42 | -12 | L | Left I-IV |
| | -3.45 | -18 | -26 | -16 | L | Parahippocampal Gyrus, posterior division |
| 92 | -3.33 | -20 | 48 | 44 | L | Frontal Pole |
| | -2.89 | -22 | 40 | 50 | L | Frontal Pole |
| | -2.78 | -24 | 52 | 40 | L | Frontal Pole |
| 89 | 3.69 | 12 | -64 | 6 | R | Intracalcarine Cortex |
| | 3.2 | 4 | -66 | 6 | R | Lingual Gyrus |
| | 2.63 | 8 | -72 | 2 | R | Lingual Gyrus |
| 89 | 5.14 | -6 | -32 | -6 | L | Brain-Stem |
| | 4.43 | 0 | -34 | -16 | C | Brain-Stem |
| | 3.68 | -4 | -28 | -10 | L | Brain-Stem |
| 72 | -3.15 | -2 | 44 | -18 | L | Frontal Medial Cortex |
| | -2.66 | 0 | 38 | -12 | C | Paracingulate Gyrus |

| Important Clusters from the PLSDA Model Classifying Surprise (movies) | | | | | | |
|---|---|---|---|---|---|---|
| | -2.19 | 4 | 38 | -18 | R | Frontal Medial Cortex |
| 66 | -3.27 | 42 | -48 | -24 | R | Temporal Occipital Fusiform Cortex |
| | -3.05 | 48 | -60 | -20 | R | Temporal Occipital Fusiform Cortex |
| | -3 | 46 | -52 | -22 | R | Temporal Occipital Fusiform Cortex |
| 66 | -3.98 | -42 | -4 | -2 | L | Insular Cortex |
| | -3.48 | -44 | 4 | -6 | L | Insular Cortex |
| | -3.05 | -46 | 14 | -6 | L | Frontal Operculum Cortex |
| 58 | 2.95 | 12 | -90 | 2 | R | Occipital Pole |
| 57 | 3.23 | -10 | -18 | 42 | L | Cingulate Gyrus, posterior division |
| | 3.17 | -14 | -28 | 38 | L | Cingulate Gyrus, posterior division |
| 56 | -3.08 | 0 | 24 | -24 | C | Subcallosal Cortex |
| | -2.58 | 4 | 34 | -22 | R | Frontal Medial Cortex |
| 56 | 4.27 | -18 | -44 | -48 | L | Left VIIIb |
| | 2.85 | -12 | -48 | -48 | L | Left IX |
| | 2.7 | -26 | -42 | -44 | L | Left VIIIa |
| 53 | -4.86 | -10 | -82 | 6 | L | Intracalcarine Cortex |
| 53 | -4.58 | 0 | -22 | 8 | C | Left Thalamus |
| | -3.28 | 0 | -14 | 10 | C | Left Thalamus |
| | -3.13 | 0 | -8 | 8 | C | Left Thalamus |
| 52 | 3.24 | -6 | -94 | 0 | L | Occipital Pole |
| | 2.18 | -6 | -90 | -10 | L | Lingual Gyrus |
| 50 | 2.66 | -14 | -80 | 40 | L | Lateral Occipital Cortex, superior division |
| | 2.62 | -10 | -74 | 42 | L | Precuneous Cortex |
| 50 | 3.76 | 16 | -48 | -52 | R | Right VIIIb |
| | 2.6 | 16 | -54 | -48 | R | Right VIIIb |
| 48 | 3.23 | -8 | -82 | -14 | L | Lingual Gyrus |
| | 2.5 | -14 | -80 | -16 | L | Occipital Fusiform Gyrus |
| | 2.37 | -2 | -88 | -14 | L | Lingual Gyrus |
| 47 | -2.79 | -46 | -58 | 34 | L | Angular Gyrus |
| | -2.72 | -44 | -64 | 38 | L | Lateral Occipital Cortex, superior division |
| | -2.62 | -48 | -68 | 42 | L | Lateral Occipital Cortex, superior division |
| 41 | 3.17 | -56 | -36 | 22 | L | Parietal Operculum Cortex |
| | 3.17 | -62 | -34 | 20 | L | Parietal Operculum Cortex |
| 39 | -3.62 | 32 | 22 | -30 | R | Temporal Pole |
| 37 | 7.76 | -2 | -38 | -66 | L | Brain-Stem |

| Important Clusters from the PLSDA Model Classifying Surprise (movies) | | | | | | |
|---|---|---|---|---|---|---|
| | 5.62 | 0 | -42 | -72 | C | Brain-Stem |
| | 3.63 | 4 | -40 | -68 | R | Brain-Stem |
| 36 | -3.63 | -18 | -38 | 4 | L | Left Hippocampus |
| | -2.23 | -10 | -34 | 6 | L | Left Thalamus |
| | -2.01 | -22 | -34 | -2 | L | Left Thalamus |
| 34 | 3.49 | -28 | -74 | -18 | L | Occipital Fusiform Gyrus |
| 34 | 3 | 0 | -94 | 20 | C | Occipital Pole |
| 34 | -3.23 | -6 | -50 | 2 | L | Cingulate Gyrus, posterior division |
| | -2.83 | -8 | -46 | -2 | L | Cingulate Gyrus, posterior division |
| 32 | 4.03 | 32 | 22 | -10 | R | Frontal Orbital Cortex |
| | 2.32 | 30 | 20 | -4 | R | Insular Cortex |
| 31 | 3.89 | 6 | -18 | -2 | R | Right Thalamus |
| | 2.69 | 12 | -14 | 2 | R | Right Thalamus |
| 31 | 2.98 | -24 | -44 | -16 | L | Temporal Occipital Fusiform Cortex |
| | 2.58 | -30 | -48 | -20 | L | Temporal Occipital Fusiform Cortex |
| | 2.25 | -30 | -44 | -14 | L | Temporal Fusiform Cortex, posterior division |
| 30 | 3.15 | 10 | -20 | 42 | R | Cingulate Gyrus, posterior division |

#### Table S15. Maps of PLSDA Coefficients (Scenarios)

*Amusement*

| Important Clusters from the PLSDA Model Classifying Amusement (scenarios) | | | | | | |
|---|---|---|---|---|---|---|
| Cluster Size (Voxels) | Max Coef | MNI | | | Hemisphere | Anatomical Label |
| | | x | y | z | | |
| 440 | 3.97 | -40 | -28 | -22 | L | Temporal Fusiform Cortex, posterior division |
| | 3.86 | -30 | -46 | -20 | L | Temporal Occipital Fusiform Cortex |
| | 3.81 | -36 | -46 | -22 | L | Temporal Occipital Fusiform Cortex |
| 157 | -4.28 | -2 | -16 | 12 | L | Left Thalamus |
| | -4.28 | 4 | -22 | 12 | R | Right Thalamus |
| | -3.84 | 0 | -14 | 4 | C | Left Thalamus |
| 141 | 5.07 | -38 | 10 | -16 | L | Insular Cortex |
| | 3.5 | -40 | 26 | -24 | L | Temporal Pole |
| | 3.44 | -46 | 14 | -8 | L | Temporal Pole |
| 134 | 3.86 | -20 | -8 | -20 | L | Left Amygdala |
| | 3.35 | -22 | -10 | -14 | L | Left Amygdala |
| | 3.21 | -30 | 0 | -14 | L | Left Amygdala |
| 106 | 4.26 | -36 | 16 | -28 | L | Temporal Pole |
| | 4.04 | -30 | 8 | -28 | L | Temporal Pole |
| | 3.6 | -24 | 10 | -30 | L | Temporal Pole |
| 93 | -3.64 | 64 | -52 | 30 | R | Angular Gyrus |
| | -3.09 | 60 | -50 | 40 | R | Angular Gyrus |
| | -2.73 | 56 | -54 | 44 | R | Angular Gyrus |
| 85 | -3.69 | 28 | 14 | 64 | R | Superior Frontal Gyrus |
| | -2.86 | 32 | 6 | 66 | R | Middle Frontal Gyrus |
| | -2.63 | 24 | 12 | 56 | R | Superior Frontal Gyrus |
| 82 | 5.4 | 30 | 14 | -28 | R | Temporal Pole |
| 75 | 3.93 | -4 | 36 | -30 | L | Frontal Medial Cortex |
| | 3.83 | 2 | 42 | -30 | R | Frontal Medial Cortex |
| | 3.3 | 2 | 32 | -30 | R | Frontal Medial Cortex |
| 71 | 3.02 | -40 | 28 | 22 | L | Middle Frontal Gyrus |
| | 2.92 | -42 | 22 | 24 | L | Middle Frontal Gyrus |
| | 2.71 | -44 | 16 | 28 | L | Inferior Frontal Gyrus, pars opercularis |
| 61 | -4.23 | 26 | 6 | -20 | R | Temporal Pole / Frontal Orbital Cortex |
| | -3.44 | 20 | 6 | -24 | R | Temporal Pole |
| | -2.22 | 34 | 12 | -18 | R | Insular Cortex |

| Important Clusters from the PLSDA Model Classifying Amusement (scenarios) | | | | | | |
|---|---|---|---|---|---|---|
| 58 | -3.18 | -60 | -52 | 38 | L | Supramarginal Gyrus, posterior division |
| | -2.49 | -52 | -54 | 40 | L | Angular Gyrus |
| 57 | -4.02 | -14 | -40 | -50 | L | Left VIIIb |
| | -3.27 | -20 | -38 | -52 | L | Left VIIIb |
| | -3.13 | -18 | -36 | -46 | L | Left X |
| 56 | -5.87 | 0 | -20 | -44 | C | Brain-Stem |
| | -5.25 | -8 | -34 | -54 | L | Brain-Stem |
| | -4.06 | 0 | -26 | -48 | C | Brain-Stem |
| 53 | -3.07 | 56 | 16 | 38 | R | Middle Frontal Gyrus |
| | -2.86 | 48 | 22 | 48 | R | Middle Frontal Gyrus |
| | -2.84 | 52 | 16 | 46 | R | Middle Frontal Gyrus |
| 47 | -3.8 | 46 | -2 | -4 | R | Planum Polare |
| | -2.8 | 46 | -8 | -2 | R | Planum Polare |
| 43 | -4.96 | 12 | -44 | -60 | R | Right VIIIb |
| | -3.32 | 18 | -44 | -58 | R | Right VIIIb |
| | -2.78 | 14 | -44 | -46 | R | Right IX |
| 43 | 3.19 | 36 | -44 | -24 | R | Temporal Occipital Fusiform Cortex |
| | 2.86 | 46 | -44 | -18 | R | Inferior Temporal Gyrus, temporooccipital part / Temporal Occipital Fusiform Cortex |
| 41 | 3.63 | 0 | -98 | -6 | C | Occipital Pole |
| | 3.08 | 2 | -100 | 2 | R | Occipital Pole |
| | 2.76 | 8 | -98 | -6 | R | Occipital Pole |
| 41 | -5.2 | 2 | -50 | -54 | R | Right IX |
| | -3.68 | 6 | -44 | -52 | R | Brain-Stem |
| | -2.54 | 8 | -54 | -54 | R | Right IX |
| 39 | 3.13 | 38 | 10 | 28 | R | Precentral Gyrus |
| 38 | 3.29 | -12 | -46 | 2 | L | Cingulate Gyrus, posterior division |
| | 2.75 | -6 | -54 | -4 | L | Left V |
| 37 | 3.09 | 40 | 18 | -12 | R | Frontal Orbital Cortex |
| | 2.33 | 54 | 28 | -6 | R | Inferior Frontal Gyrus, pars triangularis |
| | 2.3 | 48 | 24 | -8 | R | Frontal Orbital Cortex |
| 37 | 5.05 | 40 | 8 | -18 | R | Temporal Pole |
| | 2.85 | 38 | 14 | -20 | R | Temporal Pole |
| 37 | 3.41 | 22 | -4 | -20 | R | Right Amygdala |
| | 2.78 | 28 | -16 | -16 | R | Right Hippocampus |

| Important Clusters from the PLSDA Model Classifying Amusement (scenarios) | | | | | | |
|---|---|---|---|---|---|---|
| 37 | -3.6 | 22 | 68 | -10 | R | Frontal Pole |
| | -2.82 | 28 | 68 | -4 | R | Frontal Pole |
| 37 | -4.54 | 4 | -14 | -28 | R | Brain-Stem |
| | -3.4 | 4 | -16 | -38 | R | Brain-Stem |
| 37 | 3.42 | 54 | -28 | 28 | R | Parietal Operculum Cortex |
| | 2.39 | 62 | -28 | 26 | R | Supramarginal Gyrus, anterior division |
| 36 | 3.42 | -14 | -40 | -10 | L | Parahippocampal Gyrus, posterior division |
| | 2.88 | -10 | -36 | -2 | L | Left Thalamus |
| | 2.78 | -20 | -40 | -16 | L | Parahippocampal Gyrus, posterior division |
| 35 | 3.94 | -2 | 6 | -16 | L | Subcallosal Cortex |
| | 3.12 | -2 | 14 | -14 | L | Subcallosal Cortex |
| | 2.89 | 0 | 6 | -10 | C | Subcallosal Cortex |
| 35 | -2.78 | -24 | -70 | 62 | L | Lateral Occipital Cortex, superior division |
| | -2.72 | -20 | -70 | 56 | L | Lateral Occipital Cortex, superior division |
| | -2.63 | -18 | -74 | 60 | L | Lateral Occipital Cortex, superior division |
| 34 | 3.34 | 16 | -46 | -4 | R | Lingual Gyrus |
| | 2.95 | 14 | -48 | 2 | R | Cingulate Gyrus, posterior division |
| 33 | 3.52 | -2 | 70 | 6 | L | Frontal Pole |
| | 3.19 | -12 | 68 | 2 | L | Frontal Pole |
| 33 | 3.25 | -56 | 2 | -12 | L | Superior Temporal Gyrus, anterior division |
| | 2.13 | -52 | 0 | -16 | L | Superior Temporal Gyrus, anterior division |
| 32 | 4.21 | -50 | -30 | 14 | L | Parietal Operculum Cortex |
| 32 | 4.38 | 34 | -86 | -18 | R | Lateral Occipital Cortex, inferior division |
| | 2.92 | 28 | -90 | -20 | R | Occipital Fusiform Gyrus |
| 32 | -4.88 | -18 | -24 | -18 | L | Parahippocampal Gyrus, posterior division |
| | -3.1 | -18 | -30 | -22 | L | Parahippocampal Gyrus, posterior division |
| 30 | 8.25 | 6 | -44 | -72 | R | Brain-Stem |
| | 7.88 | -2 | -40 | -70 | L | Brain-Stem |
| | 3.95 | 2 | -38 | -66 | R | Brain-Stem |

*Anger*

| Important Clusters from the PLSDA Model Classifying Anger (scenarios) | | | | | | |
|---|---|---|---|---|---|---|
| **Cluster Size (Voxels)** | **Max Coef** | **MNI** | | | **Hemisphere** | **Anatomical Label** |
| | | **x** | **y** | **z** | | |
| 831 | 4.14 | -4 | -48 | 38 | L | Precuneous Cortex |
| | 4 | 0 | -56 | 44 | C | Precuneous Cortex |
| | 3.68 | 2 | -64 | 46 | R | Precuneous Cortex |
| 538 | 4.63 | 56 | -54 | 22 | R | Angular Gyrus |
| | 4.24 | 60 | -60 | 26 | R | Lateral Occipital Cortex, superior division |
| | 4.11 | 46 | -52 | 22 | R | Angular Gyrus |
| 493 | 5.05 | -44 | -58 | 32 | L | Angular Gyrus |
| | 3.38 | -56 | -62 | 18 | L | Lateral Occipital Cortex, superior division |
| | 3.36 | -56 | -64 | 28 | L | Lateral Occipital Cortex, superior division |
| 349 | -5.26 | -54 | -62 | -20 | L | Inferior Temporal Gyrus, temporooccipital part |
| | -4.16 | -50 | -66 | -18 | L | Lateral Occipital Cortex, inferior division |
| | -3.15 | -36 | -50 | -22 | L | Temporal Occipital Fusiform Cortex |
| 255 | -5.03 | -26 | 4 | -20 | L | Temporal Pole |
| | -4.36 | -24 | -2 | -20 | L | Left Amygdala |
| | -4.22 | -34 | 14 | -24 | L | Temporal Pole |
| 239 | -3.53 | 10 | -84 | 6 | R | Intracalcarine Cortex |
| | -3.44 | -10 | -72 | 10 | L | Intracalcarine Cortex |
| | -3.43 | 12 | -66 | 8 | R | Intracalcarine Cortex |
| 175 | 3.35 | 4 | 56 | 26 | R | Superior Frontal Gyrus |
| | 2.91 | 0 | 62 | 14 | C | Frontal Pole |
| | 2.82 | -4 | 46 | 20 | L | Paracingulate Gyrus |
| 173 | -3.32 | -26 | -80 | 32 | L | Lateral Occipital Cortex, superior division |
| | -3.11 | -28 | -82 | 24 | L | Lateral Occipital Cortex, superior division |
| | -3.01 | -28 | -76 | 24 | L | Lateral Occipital Cortex, superior division |
| 171 | 3.66 | 60 | -18 | -8 | R | Middle Temporal Gyrus, posterior division |
| | 3.3 | 52 | -22 | -10 | R | Middle Temporal Gyrus, posterior division |
| | 3.29 | 68 | -8 | -10 | R | Middle Temporal Gyrus, posterior division |
| 151 | -4.69 | 12 | -2 | -20 | R | Right Amygdala |
| | -4.39 | 28 | 12 | -24 | R | Frontal Orbital Cortex |
| | -4.28 | 10 | -6 | -24 | R | Parahippocampal Gyrus, anterior division |
| 134 | -3.34 | 46 | 6 | 34 | R | Precentral Gyrus |
| | -3.28 | 42 | 10 | 30 | R | Precentral Gyrus |

| Important Clusters from the PLSDA Model Classifying Anger (scenarios) | | | | | | |
|---|---|---|---|---|---|---|
| | -3 | 36 | 6 | 30 | R | Precentral Gyrus |
| 96 | -3.79 | -38 | 8 | 28 | L | Precentral Gyrus |
| | -3.29 | -44 | 8 | 32 | L | Middle Frontal Gyrus |
| | -2.82 | -46 | 2 | 34 | L | Precentral Gyrus |
| 90 | -3.46 | 36 | 36 | -12 | R | Frontal Pole |
| | -3.22 | 26 | 34 | -14 | R | Frontal Pole |
| | -2.71 | 22 | 30 | -20 | R | Frontal Orbital Cortex |
| 80 | 3.36 | 52 | -2 | -36 | R | Inferior Temporal Gyrus, anterior division |
| | 2.74 | 60 | 0 | -30 | R | Middle Temporal Gyrus, anterior division |
| | 2.71 | 46 | 8 | -36 | R | Temporal Pole |
| 68 | 4.85 | -16 | -24 | -20 | L | Parahippocampal Gyrus, posterior division |
| | 3.69 | -18 | -28 | -12 | L | Parahippocampal Gyrus, posterior division |
| 67 | -3.64 | -42 | -22 | 12 | L | Heschl's Gyrus (includes H1 and H2) |
| | -3.56 | -50 | -32 | 14 | L | Parietal Operculum Cortex |
| | -2.24 | -52 | -26 | 12 | L | Parietal Operculum Cortex |
| 59 | -2.97 | -42 | -86 | -12 | L | Lateral Occipital Cortex, inferior division |
| | -2.72 | -36 | -92 | -8 | L | Occipital Pole |
| | -2.15 | -38 | -88 | -16 | L | Lateral Occipital Cortex, inferior division |
| 57 | 4.76 | -4 | -30 | 8 | L | Left Thalamus |
| | 3.17 | -8 | -36 | 4 | L | Left Thalamus |
| | 2.64 | -14 | -32 | 10 | L | Left Thalamus |
| 57 | -2.7 | 32 | -76 | 32 | R | Lateral Occipital Cortex, superior division |
| | -2.51 | 34 | -82 | 26 | R | Lateral Occipital Cortex, superior division |
| 52 | -4.01 | 44 | -16 | 12 | R | Central Opercular Cortex |
| | -2.38 | 40 | -20 | 10 | R | Heschl's Gyrus (includes H1 and H2) |
| 52 | -5.62 | 14 | -34 | -14 | R | Parahippocampal Gyrus, posterior division |
| | -4.34 | 18 | -30 | -20 | R | Parahippocampal Gyrus, posterior division |
| | -3.79 | 12 | -38 | -10 | R | Lingual Gyrus |
| 51 | -4.19 | 0 | 14 | 24 | C | Cingulate Gyrus, anterior division |
| | -3.22 | 0 | 18 | 32 | C | Cingulate Gyrus, anterior division |
| 49 | -3.22 | -18 | -56 | 2 | L | Lingual Gyrus |
| | -3.04 | -14 | -54 | -2 | L | Lingual Gyrus |
| | -2.47 | -20 | -58 | 8 | L | Precuneous Cortex |
| 49 | 3.38 | 8 | -22 | -44 | R | Brain-Stem |
| | 3.21 | -4 | -18 | -42 | L | Brain-Stem |

| Important Clusters from the PLSDA Model Classifying Anger (scenarios) | | | | | | |
|---|---|---|---|---|---|---|
| | 2.95 | 2 | -24 | -48 | R | Brain-Stem |
| 47 | 5.73 | 16 | -14 | -28 | R | Parahippocampal Gyrus, anterior division |
| | 4.67 | 20 | -22 | -26 | R | Parahippocampal Gyrus, anterior division |
| | 3.52 | 18 | -20 | -20 | R | Parahippocampal Gyrus, anterior division |
| 46 | -3.58 | 12 | -20 | 2 | R | Right Thalamus |
| | -3.08 | 8 | -28 | -2 | R | Brain-Stem |
| | -2.69 | 2 | -26 | -4 | R | Brain-Stem |
| 46 | -3.66 | -50 | 42 | 14 | L | Frontal Pole |
| | -2.25 | -42 | 34 | 12 | L | Inferior Frontal Gyrus, pars triangularis |
| 44 | 4.09 | 0 | -16 | 0 | C | Left Thalamus |
| | 3.02 | 6 | -8 | 0 | R | Right Thalamus |
| | 2.78 | -6 | -8 | 0 | L | Left Thalamus |
| 44 | 3.57 | -44 | 0 | -10 | L | Planum Polare |
| | 2.75 | -44 | -6 | -4 | L | Insular Cortex |
| | 2.3 | -46 | 0 | -4 | L | Planum Polare |
| 41 | 4.18 | -40 | 14 | -14 | L | Insular Cortex |
| | 3.06 | -32 | 10 | -20 | L | Frontal Orbital Cortex |
| 40 | 5.33 | -8 | -34 | -50 | L | Brain-Stem |
| | 3.67 | -8 | -40 | -54 | L | Brain-Stem |
| | 2.41 | -2 | -32 | -50 | L | Brain-Stem |
| 39 | 3.84 | 38 | 16 | -16 | R | Insular Cortex / Frontal Orbital Cortex |
| | 3.27 | 42 | 10 | -14 | R | Insular Cortex |
| 38 | 3.18 | 24 | 62 | 30 | R | Frontal Pole |
| | 2.77 | 28 | 60 | 24 | R | Frontal Pole |
| 37 | -4.72 | 4 | -42 | -24 | R | Brain-Stem |
| | -3.62 | 4 | -38 | -18 | R | Brain-Stem |
| | -3.26 | -4 | -42 | -26 | L | Brain-Stem |
| 36 | -3.75 | 36 | 10 | -18 | R | Insular Cortex |
| | -2.69 | 44 | 10 | -18 | R | Temporal Pole |
| 35 | -3.14 | 52 | 14 | -6 | R | Temporal Pole |
| 35 | -5.77 | -14 | -34 | -16 | L | Parahippocampal Gyrus, posterior division |
| | -2.05 | -14 | -38 | -22 | L | Left I-IV |
| 34 | -2.83 | 62 | -48 | -12 | R | Middle Temporal Gyrus, temporooccipital part / Inferior Temporal Gyrus, temporooccipital part |
| | -2.47 | 60 | -42 | -10 | R | Middle Temporal Gyrus, temporooccipital part |

| Important Clusters from the PLSDA Model Classifying Anger (scenarios) | | | | | | |
|---|---|---|---|---|---|---|
| 34 | -3.36 | -34 | 36 | -12 | L | Frontal Orbital Cortex |
| 34 | -3.91 | 2 | 40 | -28 | R | Frontal Medial Cortex |
| | -2.19 | 4 | 32 | -30 | R | Frontal Medial Cortex |
| 33 | 3.09 | -44 | 18 | 52 | L | Middle Frontal Gyrus |
| 32 | -3.68 | 26 | -94 | -18 | R | Occipital Pole |
| | -2.34 | 32 | -88 | -16 | R | Lateral Occipital Cortex, inferior division |
| 32 | -3.05 | 46 | -50 | -12 | R | Inferior Temporal Gyrus, temporooccipital part |
| | -2.49 | 44 | -46 | -20 | R | Temporal Occipital Fusiform Cortex |
| | -2.26 | 48 | -56 | -8 | R | Inferior Temporal Gyrus, temporooccipital part |
| 31 | 3.13 | 0 | -22 | 42 | C | Cingulate Gyrus, posterior division |
| | 2.41 | -2 | -16 | 38 | L | Cingulate Gyrus, posterior division |

*Anxiety*

| Important Clusters from the PLSDA Model Classifying Anxiety (scenarios) | | | | | | |
|---|---|---|---|---|---|---|
| **Cluster Size (Voxels)** | **Max Coef** | **MNI** | | | **Hemisphere** | **Anatomical Label** |
| | | **x** | **y** | **z** | | |
| 305 | 4.59 | -54 | 12 | -22 | L | Temporal Pole |
| | 3.71 | -56 | 8 | -18 | L | Temporal Pole |
| | 3.6 | -54 | -6 | -12 | L | Superior Temporal Gyrus, anterior division |
| 241 | 3.66 | 56 | 4 | -12 | R | Superior Temporal Gyrus, anterior division |
| | 3.49 | 52 | -10 | -14 | R | Superior Temporal Gyrus, posterior division |
| | 3.29 | 48 | -26 | -6 | R | Middle Temporal Gyrus, posterior division |
| 153 | -4.59 | -16 | -2 | -16 | L | Left Amygdala |
| | -4.23 | -26 | 2 | -16 | L | Parahippocampal Gyrus, anterior division |
| | -3.87 | -20 | -10 | -14 | L | Left Amygdala |
| 127 | 4.89 | 2 | -14 | -32 | R | Brain-Stem |
| | 4.73 | -14 | -28 | -42 | L | Brain-Stem |
| | 4.24 | -10 | -18 | -38 | L | Brain-Stem |
| 87 | -3.69 | -22 | 30 | -16 | L | Frontal Orbital Cortex |
| | -3.26 | -30 | 36 | -18 | L | Frontal Pole |
| | -2.4 | -36 | 34 | -14 | L | Frontal Orbital Cortex |
| 67 | -4.31 | -4 | -32 | -14 | L | Brain-Stem |
| | -4.07 | 2 | -30 | -14 | R | Brain-Stem |
| | -2.84 | 0 | -28 | -20 | C | Brain-Stem |
| 65 | -5.17 | 12 | -38 | -10 | R | Lingual Gyrus |
| | -4.3 | 10 | -44 | 0 | R | Cingulate Gyrus, posterior division |
| | -3.41 | 12 | -50 | 2 | R | Lingual Gyrus |
| 59 | -3.49 | -10 | -84 | 46 | L | Lateral Occipital Cortex, superior division |
| | -3.02 | -12 | -78 | 40 | L | Precuneous Cortex |
| 58 | 3.53 | -42 | -56 | 28 | L | Angular Gyrus |
| | 2.95 | -50 | -58 | 28 | L | Angular Gyrus |
| | 2.7 | -52 | -62 | 32 | L | Lateral Occipital Cortex, superior division |
| 58 | 2.88 | 48 | 4 | 36 | R | Precentral Gyrus |
| | 2.84 | 52 | 8 | 38 | R | Precentral Gyrus |
| | 2.68 | 50 | 16 | 34 | R | Middle Frontal Gyrus |
| 58 | 3.7 | 0 | -40 | 22 | C | Cingulate Gyrus, posterior division |
| | 3.14 | 0 | -44 | 14 | C | Cingulate Gyrus, posterior division |
| | 2.95 | 2 | -52 | 22 | R | Cingulate Gyrus, posterior division |

| Important Clusters from the PLSDA Model Classifying Anxiety (scenarios) | | | | | | |
|---|---|---|---|---|---|---|
| 57 | -3.72 | 0 | 24 | 32 | C | Cingulate Gyrus, anterior division |
| | -2.59 | 2 | 26 | 38 | R | Paracingulate Gyrus |
| | -2.59 | 0 | 16 | 36 | C | Cingulate Gyrus, anterior division |
| 56 | -4.68 | 20 | 0 | -18 | R | Right Amygdala |
| | -3.34 | 12 | -8 | -18 | R | Right Hippocampus |
| | -2.56 | 14 | -12 | -26 | R | Parahippocampal Gyrus, anterior division |
| 55 | 3.25 | -2 | -54 | 28 | L | Cingulate Gyrus, posterior division |
| | 2.96 | 0 | -62 | 26 | C | Precuneous Cortex |
| | 2.36 | 4 | -50 | 30 | R | Cingulate Gyrus, posterior division |
| 51 | -3.29 | -18 | 12 | -20 | L | Frontal Orbital Cortex |
| | -3.03 | -20 | 10 | -26 | L | Frontal Orbital Cortex |
| | -2.45 | -14 | 8 | -22 | L | Frontal Orbital Cortex |
| 50 | 4.13 | 48 | 16 | -30 | R | Temporal Pole |
| 49 | -3.91 | 0 | 10 | -4 | C | Subcallosal Cortex |
| | -3.12 | -2 | 2 | -12 | L | Left Cerebral Cortex |
| | -2.97 | -2 | 10 | -12 | L | Subcallosal Cortex |
| 44 | -2.84 | 0 | 36 | 32 | C | Paracingulate Gyrus |
| | -2.35 | -2 | 34 | 42 | L | Superior Frontal Gyrus |
| | -2.2 | -6 | 34 | 30 | L | Paracingulate Gyrus |
| 44 | 4.21 | -4 | -26 | 8 | L | Left Thalamus |
| | 2.6 | -4 | -28 | 2 | L | Left Thalamus |
| | 2.36 | 2 | -20 | 6 | R | Right Thalamus |
| 43 | -5.93 | -36 | 12 | -14 | L | Insular Cortex |
| 43 | 3.19 | -6 | -60 | -46 | L | Left IX |
| | 2.66 | 0 | -56 | -52 | C | Right IX |
| | 2.48 | -4 | -54 | -48 | L | Left IX |
| 39 | -2.85 | 14 | -56 | 16 | R | Precuneous Cortex |
| | -2.76 | 10 | -56 | 10 | R | Precuneous Cortex |
| | -2.51 | 20 | -58 | 16 | R | Precuneous Cortex |
| 38 | 3.7 | -2 | 26 | -14 | L | Subcallosal Cortex |
| | 2.41 | 4 | 18 | -12 | R | Subcallosal Cortex |
| | 2.27 | 0 | 22 | -10 | C | Subcallosal Cortex |
| 38 | -4.28 | 40 | 16 | -14 | R | Insular Cortex |
| | -2.43 | 38 | 8 | -18 | R | Temporal Pole |
| 31 | -3.4 | -24 | -10 | -44 | L | Temporal Fusiform Cortex, posterior division |

| Important Clusters from the PLSDA Model Classifying Anxiety (scenarios) | | | | | | |
|---|---|---|---|---|---|---|
| | -2.44 | -26 | -6 | -38 | L | Parahippocampal Gyrus, anterior division |
| | -2.44 | -26 | -6 | -48 | L | Temporal Fusiform Cortex, anterior division |
| 31 | 3 | -60 | -48 | 6 | L | Middle Temporal Gyrus, temporooccipital part |
| | 2.3 | -64 | -46 | 10 | L | Supramarginal Gyrus, posterior division |
| 31 | -3.47 | -32 | -28 | -20 | L | Parahippocampal Gyrus, posterior division |
| | -2.39 | -34 | -34 | -20 | L | Temporal Fusiform Cortex, posterior division |
| 31 | -3.28 | 22 | -36 | -16 | R | Parahippocampal Gyrus, posterior division |
| | -2.71 | 28 | -40 | -12 | R | Lingual Gyrus |
| 30 | 3.07 | -48 | 10 | -40 | L | Temporal Pole |
| | 2.89 | -50 | 14 | -36 | L | Temporal Pole |
| 30 | -3.63 | -26 | -46 | -16 | L | Temporal Occipital Fusiform Cortex |
| | -2.56 | -24 | -46 | -10 | L | Lingual Gyrus |

*Awe*

| Important Clusters from the PLSDA Model Classifying Awe (scenarios) | | | | | | |
|---|---|---|---|---|---|---|
| **Cluster Size (Voxels)** | **Max Coef** | **MNI** | | | **Hemisphere** | **Anatomical Label** |
| | | **x** | **y** | **z** | | |
| 1362 | -5 | -42 | -56 | 26 | L | Angular Gyrus |
| | -4.84 | -50 | -38 | 0 | L | Middle Temporal Gyrus, posterior division |
| | -4.73 | -56 | -60 | 22 | L | Angular Gyrus |
| 664 | -4.05 | 52 | -12 | -10 | R | Superior Temporal Gyrus, posterior division |
| | -3.99 | 58 | -10 | -8 | R | Superior Temporal Gyrus, posterior division |
| | -3.66 | 48 | -36 | 2 | R | Superior Temporal Gyrus, posterior division |
| 590 | -4.9 | -8 | -74 | 12 | L | Intracalcarine Cortex |
| | -4.48 | 12 | -72 | 12 | R | Intracalcarine Cortex |
| | -4.12 | 14 | -64 | 6 | R | Intracalcarine Cortex |
| 273 | -3.87 | 2 | 54 | 28 | R | Superior Frontal Gyrus |
| | -3.38 | 0 | 52 | 34 | C | Superior Frontal Gyrus |
| | -3.21 | 0 | 50 | 40 | C | Superior Frontal Gyrus |
| 256 | -4.03 | 48 | -54 | 22 | R | Angular Gyrus |
| | -3.98 | 60 | -56 | 20 | R | Angular Gyrus |
| | -3.28 | 54 | -58 | 26 | R | Angular Gyrus |
| 238 | 3.78 | -22 | -44 | -14 | L | Lingual Gyrus |
| | 3.66 | -26 | -38 | -16 | L | Parahippocampal Gyrus, posterior division |
| | 3.24 | -28 | -32 | -22 | L | Temporal Fusiform Cortex, posterior division |
| 208 | -5.14 | 4 | -20 | 8 | R | Right Thalamus |
| | -4.16 | -4 | -24 | 10 | L | Left Thalamus |
| | -3.44 | -2 | -18 | 12 | L | Left Thalamus |
| 163 | -3.11 | 0 | -56 | 40 | C | Precuneous Cortex |
| | -2.86 | 2 | -60 | 44 | R | Precuneous Cortex |
| | -2.85 | -2 | -50 | 38 | L | Precuneous Cortex |
| 154 | 3.64 | 28 | -38 | -14 | R | Parahippocampal Gyrus, posterior division |
| | 3.48 | 22 | -40 | -16 | R | Temporal Occipital Fusiform Cortex |
| | 2.8 | 22 | -34 | -18 | R | Parahippocampal Gyrus, posterior division |
| 145 | 4.5 | 18 | -18 | -26 | R | Parahippocampal Gyrus, anterior division |
| | 4.48 | 12 | -6 | -20 | R | Right Hippocampus |
| | 4.16 | 16 | -12 | -20 | R | Right Hippocampus |
| 133 | 3.91 | 16 | -54 | 18 | R | Precuneous Cortex |
| | 3.54 | 8 | -52 | 6 | R | Precuneous Cortex |

| Important Clusters from the PLSDA Model Classifying Awe (scenarios) | | | | | | |
|---|---|---|---|---|---|---|
| | 3.45 | 8 | -56 | 14 | R | Precuneous Cortex |
| 111 | -3.32 | -46 | 2 | 56 | L | Middle Frontal Gyrus |
| | -3.15 | -42 | -2 | 50 | L | Precentral Gyrus |
| | -2.63 | -36 | -4 | 46 | L | Precentral Gyrus |
| 105 | -4.15 | 38 | 22 | -22 | R | Frontal Orbital Cortex |
| | -3.48 | 44 | 24 | -18 | R | Frontal Orbital Cortex |
| | -2.47 | 44 | 26 | -28 | R | Temporal Pole |
| 104 | -4.02 | 44 | -76 | -18 | R | Lateral Occipital Cortex, inferior division |
| | -2.71 | 44 | -82 | -14 | R | Lateral Occipital Cortex, inferior division |
| | -2.45 | 44 | -76 | -10 | R | Lateral Occipital Cortex, inferior division |
| 102 | 4.99 | -18 | -22 | -18 | L | Parahippocampal Gyrus, anterior division |
| | 3.71 | -12 | -8 | -22 | L | Parahippocampal Gyrus, anterior division |
| | 3.39 | -22 | 2 | -22 | L | Parahippocampal Gyrus, anterior division |
| 93 | -3.28 | -56 | 10 | -22 | L | Temporal Pole |
| | -2.69 | -52 | 12 | -26 | L | Temporal Pole |
| | -2.35 | -46 | 12 | -32 | L | Temporal Pole |
| 89 | 3.49 | 2 | 14 | -12 | R | Subcallosal Cortex |
| | 2.77 | 0 | 26 | -4 | C | Subcallosal Cortex |
| | 2.64 | -4 | 30 | -16 | L | Subcallosal Cortex |
| 87 | 3.72 | -54 | -62 | -22 | L | Inferior Temporal Gyrus, temporooccipital part |
| | 2.79 | -60 | -56 | -12 | L | Middle Temporal Gyrus, temporooccipital part / Inferior Temporal Gyrus, temporooccipital part |
| | 2.39 | -62 | -60 | -8 | L | Middle Temporal Gyrus, temporooccipital part |
| 87 | -3.86 | -36 | 20 | -24 | L | Frontal Orbital Cortex |
| | -2.5 | -30 | 14 | -18 | L | Frontal Orbital Cortex |
| | -2.48 | -34 | 12 | -26 | L | Temporal Pole |
| 85 | -3.15 | 32 | -92 | 6 | R | Occipital Pole |
| | -3.08 | 34 | -92 | 14 | R | Occipital Pole |
| | -2.2 | 26 | -94 | 10 | R | Occipital Pole |
| 81 | -3.05 | -54 | 22 | -4 | L | Inferior Frontal Gyrus, pars triangularis |
| | -2.9 | -54 | 22 | 6 | L | Inferior Frontal Gyrus, pars triangularis |
| | -2.68 | -54 | 32 | 4 | L | Inferior Frontal Gyrus, pars triangularis |
| 78 | -4.06 | -2 | -52 | -36 | L | Vermis IX |
| | -2.48 | 6 | -52 | -42 | R | Right IX |
| 72 | -4.43 | -2 | 8 | 72 | L | Juxtapositional Lobule Cortex (formerly Supplementary Motor Cortex) |

| Important Clusters from the PLSDA Model Classifying Awe (scenarios) | | | | | | |
|---|---|---|---|---|---|---|
| | -3.35 | -2 | 14 | 70 | L | Superior Frontal Gyrus |
| | -2.6 | -2 | -2 | 74 | L | Juxtapositional Lobule Cortex (formerly Supplementary Motor Cortex) |
| 65 | 3.99 | -12 | -58 | 16 | L | Precuneous Cortex |
| | 2.97 | -18 | -60 | 18 | L | Precuneous Cortex |
| 64 | 3.47 | 64 | -18 | 12 | R | Planum Temporale |
| 63 | -3.42 | 28 | -90 | -20 | R | Occipital Fusiform Gyrus |
| | -2.95 | 30 | -84 | -20 | R | Occipital Fusiform Gyrus |
| | -2.36 | 34 | -92 | -14 | R | Occipital Pole |
| 53 | 4.39 | -44 | -4 | -8 | L | Planum Polare |
| | 3.19 | -44 | 6 | -14 | L | Temporal Pole |
| 51 | -2.93 | 44 | 24 | 22 | R | Middle Frontal Gyrus |
| | -2.36 | 42 | 20 | 28 | R | Middle Frontal Gyrus |
| 50 | 2.84 | -58 | -6 | 26 | L | Precentral Gyrus |
| | 2.43 | -64 | -2 | 26 | L | Precentral Gyrus |
| | 2.19 | -58 | -6 | 18 | L | Precentral Gyrus |
| 48 | -3.04 | 50 | 10 | 38 | R | Middle Frontal Gyrus |
| | -2.39 | 54 | 18 | 34 | R | Middle Frontal Gyrus |
| | -2.32 | 46 | 4 | 36 | R | Precentral Gyrus |
| 44 | -2.91 | -56 | 12 | 18 | L | Inferior Frontal Gyrus, pars opercularis |
| | -2.31 | -54 | 12 | 10 | L | Inferior Frontal Gyrus, pars opercularis |
| | -2.29 | -58 | 20 | 22 | L | Inferior Frontal Gyrus, pars opercularis |
| 44 | 2.75 | 6 | 36 | -16 | R | Frontal Medial Cortex |
| | 2.58 | 0 | 48 | -10 | C | Frontal Medial Cortex |
| | 2.18 | 2 | 40 | -12 | R | Paracingulate Gyrus |
| 42 | -2.74 | -56 | -2 | 46 | L | Precentral Gyrus |
| | -2.62 | -52 | 6 | 48 | L | Middle Frontal Gyrus / Precentral Gyrus |
| | -2.46 | -48 | 6 | 42 | L | Middle Frontal Gyrus |
| 41 | 3.42 | -48 | 42 | 14 | L | Frontal Pole |
| 41 | -3.66 | 4 | -30 | -2 | R | Brain-Stem |
| | -2.26 | -6 | -26 | -4 | L | Brain-Stem |
| 39 | 3.97 | 44 | 4 | -10 | R | Insular Cortex |
| | 2.24 | 48 | 2 | -6 | R | Planum Polare |
| 34 | 3.54 | 18 | -24 | -18 | R | Parahippocampal Gyrus, posterior division |
| | 2.67 | 22 | -18 | -12 | R | Right Hippocampus |

| Important Clusters from the PLSDA Model Classifying Awe (scenarios) | | | | | | |
|---|---|---|---|---|---|---|
| 32 | 3.99 | -38 | -22 | 4 | L | Heschl's Gyrus (includes H1 and H2) |
| 32 | -3.15 | -40 | -88 | -16 | L | Lateral Occipital Cortex, inferior division |
| | -2.39 | -42 | -82 | -18 | L | Lateral Occipital Cortex, inferior division |
| 31 | 3.63 | 24 | 4 | -18 | R | Parahippocampal Gyrus, anterior division |
| | 2.54 | 30 | 6 | -16 | R | Frontal Orbital Cortex |

*Calmness*

| Important Clusters from the PLSDA Model Classifying Calmness (scenarios) | | | | | | |
|---|---|---|---|---|---|---|
| **Cluster Size (Voxels)** | **Max Coef** | **MNI** | | | **Hemisphere** | **Anatomical Label** |
| | | **x** | **y** | **z** | | |
| 232 | 4.69 | 14 | -54 | 16 | R | Precuneous Cortex |
| | 3.63 | 8 | -40 | 4 | R | Cingulate Gyrus, posterior division |
| | 3.5 | -16 | -58 | 14 | L | Precuneous Cortex |
| 224 | 3.48 | 50 | -68 | 38 | R | Lateral Occipital Cortex, superior division |
| | 3.34 | 44 | -70 | 38 | R | Lateral Occipital Cortex, superior division |
| | 3.02 | 46 | -74 | 34 | R | Lateral Occipital Cortex, superior division |
| 214 | 4.25 | 0 | 58 | 2 | C | Frontal Pole |
| | 4.13 | 2 | 64 | 0 | R | Frontal Pole |
| | 3.09 | 10 | 40 | -4 | R | Paracingulate Gyrus |
| 132 | -3.27 | 4 | 48 | 32 | R | Superior Frontal Gyrus |
| | -3.26 | -2 | 52 | 30 | L | Superior Frontal Gyrus |
| | -3.14 | -6 | 60 | 38 | L | Frontal Pole |
| 128 | 4.1 | 2 | 2 | -8 | R | Right Cerebral Cortex |
| | 4.03 | 0 | 6 | -4 | C | Subcallosal Cortex |
| | 3.68 | -4 | 2 | -14 | L | Left Cerebral Cortex |
| 127 | -3.84 | -32 | 14 | -20 | L | Frontal Orbital Cortex |
| | -3.65 | -46 | 20 | -12 | L | Frontal Orbital Cortex |
| | -3.47 | -28 | 18 | -14 | L | Frontal Orbital Cortex |
| 102 | -3.13 | -50 | -60 | 30 | L | Angular Gyrus |
| | -3.07 | -58 | -58 | 20 | L | Angular Gyrus |
| | -2.62 | -50 | -54 | 14 | L | Angular Gyrus |
| 95 | 3.82 | -8 | -70 | 36 | L | Precuneous Cortex |
| | 3.46 | -10 | -68 | 30 | L | Precuneous Cortex |
| | 2.62 | -12 | -76 | 38 | L | Precuneous Cortex |
| 80 | 3.48 | 0 | -64 | 50 | C | Precuneous Cortex |
| | 3.15 | 0 | -68 | 56 | C | Precuneous Cortex |
| | 2.5 | 2 | -60 | 54 | R | Precuneous Cortex |
| 78 | 4.76 | 0 | 32 | 14 | C | Cingulate Gyrus, anterior division |
| | 3.73 | 2 | 34 | 6 | R | Cingulate Gyrus, anterior division |
| | 2.28 | 0 | 42 | 2 | C | Cingulate Gyrus, anterior division |
| 72 | 3.47 | 4 | -52 | -54 | R | Right IX |
| | 3.35 | 10 | -50 | -50 | R | Right IX |

| Important Clusters from the PLSDA Model Classifying Calmness (scenarios) | | | | | | |
|---|---|---|---|---|---|---|
| | 3.2 | 2 | -42 | -50 | R | Brain-Stem |
| 72 | -4.63 | 28 | 8 | -18 | R | Frontal Orbital Cortex |
| | -3.11 | 30 | 16 | -18 | R | Frontal Orbital Cortex |
| | -2.63 | 32 | 14 | -28 | R | Temporal Pole |
| 68 | 2.79 | 32 | 64 | -2 | R | Frontal Pole |
| | 2.63 | 28 | 58 | -2 | R | Frontal Pole |
| | 2.62 | 30 | 64 | 6 | R | Frontal Pole |
| 63 | 5.92 | -16 | -28 | -20 | L | Parahippocampal Gyrus, posterior division |
| | 3.3 | -18 | -24 | -16 | L | Parahippocampal Gyrus, posterior division |
| | 2.61 | -24 | -32 | -22 | L | Parahippocampal Gyrus, posterior division |
| 63 | 4.86 | 20 | -28 | -24 | R | Parahippocampal Gyrus, posterior division |
| | 2.93 | 20 | -22 | -16 | R | Right Hippocampus |
| | 2.81 | 26 | -30 | -32 | R | Right I-IV |
| 60 | -2.7 | -46 | 14 | -34 | L | Temporal Pole |
| | -2.56 | -44 | 8 | -36 | L | Temporal Pole |
| 59 | 4.02 | 22 | 10 | -26 | R | Frontal Orbital Cortex |
| | 3.34 | 28 | 2 | -26 | R | Right Amygdala |
| 58 | -2.8 | -54 | 12 | 22 | L | Inferior Frontal Gyrus, pars opercularis |
| | -2.54 | -54 | 16 | 30 | L | Inferior Frontal Gyrus, pars opercularis |
| 56 | 4.14 | 14 | -36 | -8 | R | Parahippocampal Gyrus, posterior division |
| | 2.93 | 18 | -30 | -10 | R | Parahippocampal Gyrus, posterior division |
| | 2.55 | 22 | -34 | -12 | R | Parahippocampal Gyrus, posterior division |
| 56 | -2.81 | -52 | 8 | -24 | L | Temporal Pole |
| | -2.79 | -54 | 0 | -16 | L | Superior Temporal Gyrus, anterior division |
| 55 | -4.37 | 0 | 32 | -28 | C | Frontal Medial Cortex |
| | -2.89 | 4 | 42 | -28 | R | Frontal Medial Cortex |
| 52 | -3.56 | 0 | 44 | 46 | C | Superior Frontal Gyrus |
| 51 | -3.03 | -54 | 26 | 10 | L | Inferior Frontal Gyrus, pars triangularis |
| | -2.86 | -48 | 24 | 8 | L | Inferior Frontal Gyrus, pars triangularis |
| | -2.4 | -54 | 22 | 4 | L | Inferior Frontal Gyrus, pars triangularis |
| 47 | -3.14 | 2 | -56 | -36 | R | Vermis IX |
| | -2.65 | -4 | -56 | -36 | L | Vermis IX |
| 47 | -3.21 | 0 | 32 | 54 | C | Superior Frontal Gyrus |
| | -2.54 | -8 | 40 | 56 | L | Frontal Pole |
| 46 | -2.98 | 62 | -28 | -2 | R | Superior Temporal Gyrus, posterior division |

| Important Clusters from the PLSDA Model Classifying Calmness (scenarios) | | | | | | |
|---|---|---|---|---|---|---|
| | -2.69 | 56 | -30 | -2 | R | Superior Temporal Gyrus, posterior division |
| | -2.36 | 56 | -24 | -4 | R | Superior Temporal Gyrus, posterior division |
| 39 | -2.99 | 40 | 20 | -14 | R | Frontal Orbital Cortex |
| | -2.94 | 46 | 20 | -10 | R | Frontal Orbital Cortex |
| | -2.21 | 38 | 20 | -6 | R | Insular Cortex |
| 34 | -3.52 | 44 | 4 | -10 | R | Insular Cortex |
| | -2.86 | 42 | 8 | -14 | R | Insular Cortex |
| 33 | -3.05 | 46 | -26 | -4 | R | Superior Temporal Gyrus, posterior division |
| | -2.3 | 48 | -32 | -4 | R | Middle Temporal Gyrus, posterior division |
| 33 | -2.89 | 46 | 4 | -38 | R | Inferior Temporal Gyrus, anterior division |
| | -2.54 | 48 | 6 | -32 | R | Temporal Pole |
| | -2.46 | 40 | 6 | -40 | R | Temporal Pole |
| 32 | -3.68 | 2 | -26 | -2 | R | Brain-Stem |
| | -2.93 | 6 | -28 | 4 | R | Right Thalamus |
| 31 | -3.34 | 6 | -2 | 4 | R | Right Thalamus |
| 31 | 4.63 | 0 | -48 | -42 | C | Vermis X |
| | 3.23 | -10 | -48 | -44 | L | Left IX |
| 30 | 2.93 | 4 | -84 | -14 | R | Lingual Gyrus |
| | 2.84 | 6 | -90 | -14 | R | Lingual Gyrus |
| | 2.67 | 14 | -94 | -14 | R | Occipital Pole |
| 30 | -2.92 | -48 | 34 | -4 | L | Inferior Frontal Gyrus, pars triangularis |
| | -2.18 | -52 | 28 | -4 | L | Inferior Frontal Gyrus, pars triangularis |

*Craving*

| Important Clusters from the PLSDA Model Classifying Craving (scenarios) | | | | | | |
|---|---|---|---|---|---|---|
| Cluster Size (Voxels) | Max Coef | MNI | | | Hemisphere | Anatomical Label |
| | | x | y | z | | |
| 770 | 3.85 | -38 | -68 | -20 | L | Occipital Fusiform Gyrus |
| | 3.83 | -50 | -64 | -22 | L | Inferior Temporal Gyrus, temporooccipital part |
| | 3.61 | -44 | -74 | -18 | L | Lateral Occipital Cortex, inferior division |
| 745 | 5.02 | 10 | -84 | 4 | R | Intracalcarine Cortex |
| | 4.34 | 8 | -100 | -4 | R | Occipital Pole |
| | 4.24 | 10 | -72 | 12 | R | Intracalcarine Cortex |
| 625 | 7.29 | 46 | -76 | -18 | R | Lateral Occipital Cortex, inferior division |
| | 4.04 | 32 | -90 | 4 | R | Occipital Pole |
| | 3.86 | 28 | -90 | -20 | R | Occipital Fusiform Gyrus |
| 395 | 4 | -10 | -74 | 12 | L | Intracalcarine Cortex |
| | 3.66 | -10 | -68 | 6 | L | Intracalcarine Cortex |
| | 3.62 | -18 | -64 | 4 | L | Intracalcarine Cortex |
| 274 | 4.42 | -48 | -44 | 6 | L | Middle Temporal Gyrus, temporooccipital part |
| | 4.06 | -50 | -38 | 2 | L | Superior Temporal Gyrus, posterior division |
| | 3.41 | -62 | -42 | 6 | L | Superior Temporal Gyrus, posterior division |
| 200 | -3.47 | 2 | 62 | 14 | R | Frontal Pole |
| | -3.29 | 2 | 56 | 24 | R | Superior Frontal Gyrus |
| | -2.89 | 0 | 50 | 18 | C | Paracingulate Gyrus |
| 155 | 3.93 | -36 | 8 | 28 | L | Precentral Gyrus |
| | 3.68 | -44 | 8 | 24 | L | Inferior Frontal Gyrus, pars opercularis |
| | 3.4 | -38 | 2 | 32 | L | Precentral Gyrus |
| 130 | 3.58 | -14 | -2 | -28 | L | Parahippocampal Gyrus, anterior division |
| | 3.45 | -18 | 8 | -26 | L | Frontal Orbital Cortex |
| | 3.29 | -24 | 6 | -20 | L | Frontal Orbital Cortex |
| 110 | -3.53 | 2 | -62 | 34 | R | Precuneous Cortex |
| | -2.65 | 6 | -54 | 40 | R | Precuneous Cortex |
| | -2.58 | 2 | -60 | 44 | R | Precuneous Cortex |
| 91 | 5.37 | 28 | 6 | -22 | R | Temporal Pole |
| | 3.79 | 16 | 4 | -26 | R | Parahippocampal Gyrus, anterior division |
| | 3.76 | 34 | 12 | -22 | R | Temporal Pole |
| 89 | -4.69 | 46 | 0 | -6 | R | Planum Polare |
| | -4.08 | 46 | -6 | -2 | R | Heschl's Gyrus (includes H1 and H2) |

| Important Clusters from the PLSDA Model Classifying Craving (scenarios) | | | | | | |
|---|---|---|---|---|---|---|
| | -3.05 | 44 | -14 | 0 | R | Heschl's Gyrus (includes H1 and H2) |
| 81 | 3.37 | -58 | -2 | -10 | L | Superior Temporal Gyrus, anterior division |
| | 2.99 | -58 | -10 | -8 | L | Superior Temporal Gyrus, posterior division |
| | 2.57 | -52 | -2 | -16 | L | Superior Temporal Gyrus, anterior division |
| 78 | 3 | -44 | -2 | 58 | L | Precentral Gyrus |
| | 2.98 | -46 | -10 | 58 | L | Precentral Gyrus |
| | 2.66 | -36 | -2 | 48 | L | Precentral Gyrus |
| 76 | 3 | 20 | -100 | 12 | R | Occipital Pole |
| | 2.74 | 16 | -94 | 18 | R | Occipital Pole |
| | 2.5 | 14 | -96 | 28 | R | Occipital Pole |
| 73 | 3.25 | -52 | 4 | 46 | L | Precentral Gyrus |
| | 3.11 | -54 | 10 | 44 | L | Middle Frontal Gyrus |
| | 2.43 | -56 | -2 | 46 | L | Precentral Gyrus |
| 67 | 4.13 | -22 | 30 | -16 | L | Frontal Orbital Cortex |
| | 3.47 | -24 | 34 | -12 | L | Frontal Orbital Cortex |
| 66 | 3.69 | -28 | -76 | 24 | L | Lateral Occipital Cortex, superior division |
| | 3.05 | -26 | -72 | 30 | L | Lateral Occipital Cortex, superior division |
| | 2.71 | -26 | -80 | 20 | L | Lateral Occipital Cortex, superior division |
| 64 | 4.46 | -36 | 34 | -12 | L | Frontal Orbital Cortex |
| | 2.97 | -40 | 28 | -12 | L | Frontal Orbital Cortex |
| 57 | 4.9 | 38 | 8 | -16 | R | Insular Cortex |
| | 2.53 | 40 | 2 | -14 | R | Insular Cortex |
| | 2.33 | 40 | 2 | -8 | R | Insular Cortex |
| 51 | 3.88 | 50 | -36 | 6 | R | Superior Temporal Gyrus, posterior division |
| | 2.98 | 54 | -32 | 2 | R | Superior Temporal Gyrus, posterior division |
| 49 | 3.11 | 54 | 14 | -4 | R | Temporal Pole |
| | 3.05 | 54 | 12 | -22 | R | Temporal Pole |
| | 2.64 | 54 | 18 | -14 | R | Temporal Pole |
| 44 | -3.42 | 2 | -68 | 52 | R | Precuneous Cortex |
| | -2.67 | 0 | -62 | 54 | C | Precuneous Cortex |
| 41 | 3.92 | 34 | -52 | -18 | R | Temporal Occipital Fusiform Cortex |
| | 2.96 | 32 | -44 | -22 | R | Temporal Occipital Fusiform Cortex |
| 40 | 2.85 | 34 | -74 | 28 | R | Lateral Occipital Cortex, superior division |
| | 2.52 | 32 | -66 | 30 | R | Lateral Occipital Cortex, superior division |
| 40 | 2.68 | 30 | -54 | 60 | R | Superior Parietal Lobule |

| Important Clusters from the PLSDA Model Classifying Craving (scenarios) | | | | | | |
|---|---|---|---|---|---|---|
| | 2.56 | 28 | -60 | 66 | R | Lateral Occipital Cortex, superior division |
| | 2.31 | 24 | -58 | 58 | R | Lateral Occipital Cortex, superior division |
| 39 | 2.88 | 0 | -36 | 26 | C | Cingulate Gyrus, posterior division |
| | 2.85 | 0 | -34 | 32 | C | Cingulate Gyrus, posterior division |
| | 2.41 | 0 | -28 | 34 | C | Cingulate Gyrus, posterior division |
| 39 | 3.28 | 24 | 32 | -16 | R | Frontal Orbital Cortex |
| 38 | -3.49 | 24 | -32 | -32 | R | Right I-IV |
| | -2.87 | 30 | -34 | -32 | R | Right V |
| | -2.38 | 30 | -38 | -38 | R | Right VI |
| 38 | 3.44 | 6 | -28 | -2 | R | Brain-Stem |
| | 3.4 | 0 | -24 | -8 | C | Brain-Stem |
| 36 | -3.14 | -6 | -2 | 6 | L | Left Thalamus |
| | -3 | -6 | -4 | 12 | L | Left Thalamus |
| | -2.71 | -8 | 4 | 6 | L | Left Caudate |
| 34 | 3.94 | 8 | -48 | 0 | R | Lingual Gyrus |
| | 3.3 | 10 | -44 | -4 | R | Lingual Gyrus |
| 33 | 3.34 | 8 | -80 | -44 | R | Right Crus II |
| | 2.22 | 10 | -80 | -50 | R | Right VIIb |
| 33 | -2.87 | -2 | 34 | -28 | L | Frontal Medial Cortex |
| 32 | -4.79 | -44 | -2 | -6 | L | Insular Cortex |
| | -2.41 | -44 | 4 | -12 | L | Temporal Pole |

*Disgust*

| Important Clusters from the PLSDA Model Classifying Disgust (scenarios) | | | | | | |
|---|---|---|---|---|---|---|
| Cluster Size (Voxels) | Max Coef | MNI | | | Hemisphere | Anatomical Label |
| | | x | y | z | | |
| 203 | 5.95 | -24 | 34 | -14 | L | Frontal Orbital Cortex |
| | 5.42 | -32 | 38 | -10 | L | Frontal Pole |
| | 3.64 | -30 | 38 | -16 | L | Frontal Pole |
| 161 | 3.82 | -42 | 36 | 12 | L | Frontal Pole |
| | 3.44 | -46 | 36 | 18 | L | Frontal Pole |
| | 3.08 | -54 | 34 | 16 | L | Inferior Frontal Gyrus, pars triangularis |
| 160 | 6.09 | 46 | -2 | -2 | R | Insular Cortex |
| | 4.02 | 40 | 4 | -12 | R | Insular Cortex |
| | 3.88 | 44 | 2 | -8 | R | Insular Cortex |
| 110 | -3.29 | -54 | -28 | 12 | L | Planum Temporale |
| | -3.25 | -54 | -20 | 10 | L | Heschl's Gyrus (includes H1 and H2) |
| | -3.1 | -48 | -34 | 20 | L | Parietal Operculum Cortex |
| 107 | 5.33 | 20 | 0 | -18 | R | Right Amygdala |
| | 4 | 24 | -4 | -12 | R | Right Amygdala |
| | 3.17 | 14 | -2 | -26 | R | Parahippocampal Gyrus, anterior division |
| 98 | -3.95 | 0 | 44 | -4 | C | Paracingulate Gyrus |
| | -3.19 | -8 | 42 | -12 | L | Frontal Medial Cortex |
| | -3.09 | 0 | 44 | 2 | C | Cingulate Gyrus, anterior division |
| 97 | 4.59 | 22 | 32 | -16 | R | Frontal Orbital Cortex |
| | 2.27 | 28 | 34 | -20 | R | Frontal Pole |
| 95 | 3.34 | -42 | 6 | -6 | L | Insular Cortex |
| | 3.26 | -42 | 12 | -14 | L | Insular Cortex |
| | 3.11 | -36 | 6 | -6 | L | Insular Cortex |
| 95 | 4.6 | -28 | 6 | -26 | L | Temporal Pole |
| | 4.44 | -28 | 6 | -18 | L | Frontal Orbital Cortex |
| | 2.87 | -22 | 8 | -22 | L | Frontal Orbital Cortex |
| 86 | -4.72 | -40 | -22 | 0 | L | Planum Polare |
| | -3.33 | -40 | -22 | 10 | L | Heschl's Gyrus (includes H1 and H2) |
| | -2.79 | -38 | -16 | 18 | L | Central Opercular Cortex |
| 77 | -3.64 | 50 | -30 | 18 | R | Parietal Operculum Cortex |
| | -3.25 | 60 | -24 | 14 | R | Planum Temporale |
| | -2.42 | 56 | -20 | 12 | R | Planum Temporale |

| Important Clusters from the PLSDA Model Classifying Disgust (scenarios) | | | | | | |
|---|---|---|---|---|---|---|
| 76 | -2.98 | -30 | 62 | 6 | L | Frontal Pole |
| | -2.94 | -30 | 60 | 0 | L | Frontal Pole |
| | -2.5 | -38 | 48 | -4 | L | Frontal Pole |
| 76 | 4.59 | -18 | -4 | -14 | L | Left Amygdala |
| | 3.29 | -14 | -4 | -20 | L | Left Amygdala |
| | 2.26 | -22 | -10 | -14 | L | Left Amygdala |
| 75 | 5.03 | -32 | -28 | -22 | L | Temporal Fusiform Cortex, posterior division |
| | 3.34 | -28 | -34 | -24 | L | Temporal Fusiform Cortex, posterior division |
| 74 | 5.11 | 0 | -4 | 32 | C | Cingulate Gyrus, anterior division |
| | 3.66 | 0 | -6 | 40 | C | Cingulate Gyrus, anterior division |
| | 2.35 | 4 | -12 | 38 | R | Cingulate Gyrus, anterior division |
| 61 | -3.68 | 38 | -16 | -4 | R | Insular Cortex |
| | -3.44 | 38 | -20 | 2 | R | Insular Cortex |
| | -2.3 | 38 | -18 | 8 | R | Insular Cortex |
| 58 | 4.27 | -6 | -20 | 0 | L | Left Thalamus |
| | 3.15 | -2 | -14 | 4 | L | Left Thalamus |
| 53 | -3.34 | -2 | 60 | 0 | L | Frontal Pole |
| | -2.82 | 0 | 56 | -8 | C | Frontal Pole |
| 51 | -3.99 | 52 | 20 | -8 | R | Frontal Orbital Cortex |
| | -3.89 | 46 | 20 | -2 | R | Frontal Operculum Cortex |
| | -2.33 | 40 | 22 | 0 | R | Frontal Operculum Cortex |
| 51 | -4.09 | 0 | 12 | -10 | C | Subcallosal Cortex |
| | -3.56 | 0 | 6 | -12 | C | Subcallosal Cortex |
| | -3.48 | 0 | 18 | -6 | C | Subcallosal Cortex |
| 48 | -5.05 | -14 | -18 | -26 | L | Brain-Stem |
| | -4.82 | -12 | -10 | -22 | L | Parahippocampal Gyrus, anterior division |
| | -3.72 | -16 | -18 | -20 | L | Left Hippocampus |
| 47 | 5.63 | 8 | -14 | -32 | R | Brain-Stem |
| | 3.58 | 0 | -16 | -34 | C | Brain-Stem |
| | 2.67 | -6 | -16 | -38 | L | Brain-Stem |
| 46 | 4.35 | -2 | -24 | 8 | L | Left Thalamus |
| | 4.01 | 6 | -22 | 8 | R | Right Thalamus |
| | 3.77 | -4 | -32 | 6 | L | Left Thalamus |
| 45 | 2.96 | -44 | 22 | 0 | L | Frontal Operculum Cortex |
| | 2.94 | -42 | 22 | 6 | L | Frontal Operculum Cortex |

| Important Clusters from the PLSDA Model Classifying Disgust (scenarios) | | | | | | |
|---|---|---|---|---|---|---|
| | 2.51 | -40 | 30 | -2 | L | Frontal Orbital Cortex |
| 45 | 3.77 | -48 | -50 | -18 | L | Inferior Temporal Gyrus, temporooccipital part |
| | 3.19 | -48 | -56 | -24 | L | Inferior Temporal Gyrus, temporooccipital part |
| 45 | -8.04 | -14 | -40 | -56 | L | Left VIIIb |
| | -3.83 | -8 | -42 | -56 | L | Brain-Stem |
| | -3.08 | -16 | -40 | -48 | L | Left X |
| 42 | -3.06 | 2 | -88 | -10 | R | Lingual Gyrus |
| | -3.05 | -2 | -88 | -16 | L | Lingual Gyrus |
| | -2.42 | 2 | -92 | -2 | R | Occipital Pole |
| 41 | 3.92 | -8 | -32 | -6 | L | Brain-Stem |
| | 2.68 | -16 | -34 | -4 | L | Left Thalamus |
| 39 | 3.12 | 4 | -74 | 8 | R | Intracalcarine Cortex |
| | 3.02 | 2 | -76 | 0 | R | Lingual Gyrus |
| | 2.32 | 6 | -74 | 18 | R | Intracalcarine Cortex |
| 37 | 3.93 | -12 | -74 | 10 | L | Intracalcarine Cortex |
| | 2.91 | -4 | -72 | 12 | L | Intracalcarine Cortex |
| 37 | 3.62 | 10 | -32 | -8 | R | Brain-Stem |
| | 3.21 | 12 | -28 | -14 | R | Brain-Stem |
| | 2.91 | -2 | -30 | -10 | L | Brain-Stem |
| 37 | 3.54 | 2 | 6 | 28 | R | Cingulate Gyrus, anterior division |
| 36 | 3.06 | 20 | -38 | 4 | R | Right Hippocampus |
| | 2.9 | 16 | -34 | 6 | R | Right Thalamus |
| | 2.67 | 12 | -30 | 4 | R | Right Thalamus |
| 35 | -5.63 | -4 | -42 | -70 | L | Brain-Stem |
| | -4.46 | 4 | -40 | -66 | R | Brain-Stem |
| | -2.24 | -2 | -38 | -64 | L | Brain-Stem |
| 35 | 3.62 | -24 | 6 | -40 | L | Temporal Pole |
| 34 | 3.67 | -30 | -10 | -32 | L | Parahippocampal Gyrus, anterior division |
| 31 | -4.01 | 42 | 6 | -18 | R | Temporal Pole |
| | -3.21 | 38 | 8 | -22 | R | Temporal Pole |
| 30 | 2.78 | 40 | 34 | -14 | R | Frontal Pole |
| 30 | 2.77 | -62 | -18 | 26 | L | Postcentral Gyrus |
| | 2.36 | -56 | -18 | 26 | L | Postcentral Gyrus |
| 30 | 3.11 | 0 | -56 | -36 | C | Vermis IX |
| | 2.24 | -4 | -62 | -38 | L | Vermis VIIIb |

| Important Clusters from the PLSDA Model Classifying Disgust (scenarios) | | | | | | |
|---|---|---|---|---|---|---|
| 30 | 3.99 | 0 | -32 | 26 | C | Cingulate Gyrus, posterior division |
| | 2.87 | 2 | -26 | 28 | R | Cingulate Gyrus, posterior division |
| | 2.38 | 4 | -36 | 28 | R | Cingulate Gyrus, posterior division |

*Excitement*

| Important Clusters from the PLSDA Model Classifying Excitement (scenarios) | | | | | | |
|---|---|---|---|---|---|---|
| **Cluster Size (Voxels)** | **Max Coef** | **MNI** | | | **Hemisphere** | **Anatomical Label** |
| | | **x** | **y** | **z** | | |
| 388 | 4.25 | -36 | -66 | 58 | L | Lateral Occipital Cortex, superior division |
| | 4.13 | -44 | -58 | 58 | L | Lateral Occipital Cortex, superior division |
| | 3.89 | -50 | -58 | 52 | L | Angular Gyrus |
| 282 | -3.81 | 54 | -22 | -8 | R | Middle Temporal Gyrus, posterior division |
| | -3.33 | 54 | 2 | -16 | R | Superior Temporal Gyrus, anterior division |
| | -3.32 | 50 | -18 | -10 | R | Middle Temporal Gyrus, posterior division |
| 211 | -3.17 | -60 | -60 | 14 | L | Angular Gyrus |
| | -3.16 | -60 | -54 | 12 | L | Angular Gyrus |
| | -3.04 | -54 | -48 | 6 | L | Middle Temporal Gyrus, temporooccipital part |
| 203 | -3.37 | -10 | -70 | 6 | L | Intracalcarine Cortex |
| | -3.26 | -6 | -72 | 10 | L | Intracalcarine Cortex |
| | -3.23 | 10 | -74 | 14 | R | Intracalcarine Cortex |
| 175 | -3.24 | 60 | -56 | 20 | R | Angular Gyrus |
| | -3.21 | 54 | -56 | 22 | R | Angular Gyrus |
| | -2.98 | 60 | -44 | 10 | R | Middle Temporal Gyrus, temporooccipital part |
| 147 | -3.6 | -42 | -34 | -20 | L | Temporal Fusiform Cortex, posterior division |
| | -3 | -30 | -38 | -22 | L | Temporal Fusiform Cortex, posterior division |
| | -2.84 | -36 | -40 | -24 | L | Temporal Fusiform Cortex, posterior division |
| 142 | -4.25 | -4 | -32 | -2 | L | Brain-Stem |
| | -3.47 | 8 | -30 | -2 | R | Brain-Stem |
| | -3.33 | 2 | -30 | -2 | R | Brain-Stem |
| 116 | -3.68 | -44 | 0 | 52 | L | Precentral Gyrus |
| | -3.37 | -38 | -4 | 46 | L | Precentral Gyrus |
| | -2.56 | -44 | 2 | 58 | L | Middle Frontal Gyrus |
| 114 | 3.87 | 0 | 46 | -2 | C | Paracingulate Gyrus |
| | 2.95 | -2 | 50 | -8 | L | Frontal Medial Cortex |
| | 2.44 | -8 | 52 | -2 | L | Paracingulate Gyrus |
| 112 | -3.75 | -12 | -90 | -22 | L | Occipital Fusiform Gyrus |
| | -3.69 | -10 | -98 | -16 | L | Occipital Pole |
| | -2.9 | -2 | -96 | -14 | L | Occipital Pole |
| 104 | -3.49 | 4 | -98 | 2 | R | Occipital Pole |
| | -3.24 | 4 | -98 | -4 | R | Occipital Pole |

| Important Clusters from the PLSDA Model Classifying Excitement (scenarios) | | | | | | |
|---|---|---|---|---|---|---|
| | -3.13 | 10 | -98 | -6 | R | Occipital Pole |
| 99 | -3.32 | -14 | -102 | 6 | L | Occipital Pole |
| | -2.84 | -10 | -92 | 0 | L | Occipital Pole |
| | -2.48 | -8 | -86 | 0 | L | Intracalcarine Cortex |
| 85 | -4.79 | -30 | 10 | -18 | L | Frontal Orbital Cortex |
| | -3.38 | -32 | 6 | -24 | L | Temporal Pole |
| | -3.37 | -24 | 6 | -20 | L | Frontal Orbital Cortex |
| 73 | 3.24 | -2 | -38 | 32 | L | Cingulate Gyrus, posterior division |
| | 3.08 | -2 | -30 | 36 | L | Cingulate Gyrus, posterior division |
| | 3.02 | 4 | -28 | 36 | R | Cingulate Gyrus, posterior division |
| 67 | -4.78 | 22 | -92 | -18 | R | Occipital Pole |
| | -2.92 | 30 | -88 | -18 | R | Occipital Fusiform Gyrus |
| | -2.59 | 20 | -98 | -16 | R | Occipital Pole |
| 60 | 3.27 | 54 | -22 | 14 | R | Parietal Operculum Cortex |
| | 3.11 | 68 | -18 | 12 | R | Superior Temporal Gyrus, posterior division |
| | 2.8 | 60 | -20 | 14 | R | Planum Temporale |
| 58 | 2.97 | 2 | 60 | -4 | R | Frontal Pole |
| | 2.85 | -4 | 72 | 8 | L | Frontal Pole |
| | 2.67 | -6 | 66 | 10 | L | Frontal Pole |
| 57 | 3.59 | 40 | 54 | -4 | R | Frontal Pole |
| | 2.57 | 42 | 52 | 6 | R | Frontal Pole |
| 55 | -3.11 | -38 | 8 | 28 | L | Precentral Gyrus |
| | -2.66 | -46 | 6 | 20 | L | Precentral Gyrus |
| | -2.3 | -36 | 16 | 26 | L | Inferior Frontal Gyrus, pars opercularis |
| 54 | -2.69 | -60 | -8 | -4 | L | Superior Temporal Gyrus, anterior division |
| | -2.67 | -54 | -14 | -8 | L | Superior Temporal Gyrus, posterior division |
| | -2.57 | -54 | -6 | -8 | L | Superior Temporal Gyrus, anterior division |
| 48 | 3.86 | -2 | -48 | 8 | L | Cingulate Gyrus, posterior division |
| | 3.58 | -4 | -52 | 20 | L | Cingulate Gyrus, posterior division |
| 43 | 3.16 | 26 | -62 | -8 | R | Lingual Gyrus |
| | 3.14 | 20 | -62 | -4 | R | Lingual Gyrus |
| 43 | -2.99 | 2 | 56 | 38 | R | Superior Frontal Gyrus |
| | -2.42 | 2 | 52 | 30 | R | Superior Frontal Gyrus |
| | -2.26 | 4 | 50 | 36 | R | Superior Frontal Gyrus |
| 41 | -2.98 | -10 | -50 | 0 | L | Lingual Gyrus |

| Important Clusters from the PLSDA Model Classifying Excitement (scenarios) | | | | | | |
|---|---|---|---|---|---|---|
| | -2.94 | -8 | -56 | 6 | L | Precuneous Cortex |
| | -2.5 | -16 | -52 | 0 | L | Lingual Gyrus |
| 41 | 3.34 | -28 | 26 | 58 | L | Superior Frontal Gyrus |
| | 3.33 | -24 | 32 | 56 | L | Superior Frontal Gyrus |
| | 2.21 | -22 | 28 | 60 | L | Superior Frontal Gyrus |
| 41 | 3.04 | -28 | -12 | -2 | L | Left Putamen |
| 39 | 2.82 | -40 | 52 | 2 | L | Frontal Pole |
| | 2.53 | -44 | 54 | -2 | L | Frontal Pole |
| | 2.09 | -40 | 58 | 6 | L | Frontal Pole |
| 39 | 2.64 | -32 | 58 | -6 | L | Frontal Pole |
| | 2.27 | -32 | 60 | 0 | L | Frontal Pole |
| 39 | -3.32 | 48 | 32 | -6 | R | Frontal Orbital Cortex |
| | -2.74 | 52 | 26 | -6 | R | Inferior Frontal Gyrus, pars triangularis |
| | -2.36 | 54 | 32 | -8 | R | Frontal Pole |
| 38 | -2.82 | -44 | -60 | -20 | L | Temporal Occipital Fusiform Cortex |
| | -2.25 | -44 | -54 | -22 | L | Temporal Occipital Fusiform Cortex |
| 37 | 3.23 | 0 | 24 | -16 | C | Subcallosal Cortex |
| | 2.55 | -4 | 30 | -14 | L | Subcallosal Cortex |
| | 2.37 | 0 | 36 | -14 | C | Frontal Medial Cortex / Paracingulate Gyrus |
| 37 | 2.52 | 40 | -68 | -40 | R | Right Crus I |
| | 2.37 | 30 | -70 | -40 | R | Right Crus II |
| 32 | -3.38 | 26 | 8 | -18 | R | Frontal Orbital Cortex |
| | -2.9 | 26 | 16 | -22 | R | Frontal Orbital Cortex |
| 31 | 3.04 | -44 | -12 | 0 | L | Insular Cortex |
| | 2.65 | -46 | -14 | 8 | L | Heschl's Gyrus (includes H1 and H2) |
| | 2.34 | -42 | -6 | 2 | L | Insular Cortex |
| 30 | 2.99 | 6 | -64 | -50 | R | Right IX |
| | 2.51 | 2 | -60 | -54 | R | Right IX |

*Fear*

| Important Clusters from the PLSDA Model Classifying Fear (scenarios) | | | | | | |
|---|---|---|---|---|---|---|
| Cluster Size (Voxels) | Max Coef | MNI | | | Hemisphere | Anatomical Label |
| | | x | y | z | | |
| 460 | -4.59 | 0 | 62 | -6 | C | Frontal Pole |
| | -4.11 | 2 | 64 | 4 | R | Frontal Pole |
| | -4.06 | 2 | 68 | 16 | R | Frontal Pole |
| 424 | 4.53 | 52 | -44 | 32 | R | Angular Gyrus |
| | 4.5 | 60 | -48 | 26 | R | Angular Gyrus |
| | 4.09 | 60 | -46 | 32 | R | Angular Gyrus |
| 367 | -4.28 | 36 | -66 | 58 | R | Lateral Occipital Cortex, superior division |
| | -4.24 | 38 | -56 | 64 | R | Superior Parietal Lobule |
| | -4.02 | 48 | -62 | 52 | R | Lateral Occipital Cortex, superior division |
| 332 | 3.78 | -2 | -56 | 58 | L | Precuneous Cortex |
| | 3.63 | -2 | -54 | 52 | L | Precuneous Cortex |
| | 3.39 | -2 | -42 | 50 | L | Precuneous Cortex |
| 319 | 4 | -60 | -48 | 44 | L | Supramarginal Gyrus, posterior division |
| | 3.88 | -60 | -52 | 32 | L | Supramarginal Gyrus, posterior division |
| | 3.33 | -64 | -40 | 40 | L | Supramarginal Gyrus, anterior division |
| 251 | -3.68 | 0 | -72 | 36 | C | Precuneous Cortex |
| | -3.38 | 2 | -68 | 28 | R | Precuneous Cortex |
| | -3.31 | 2 | -62 | 28 | R | Precuneous Cortex |
| 208 | -3.73 | 0 | -34 | 26 | C | Cingulate Gyrus, posterior division |
| | -3.5 | 0 | -46 | 28 | C | Cingulate Gyrus, posterior division |
| | -3.4 | 0 | -36 | 32 | C | Cingulate Gyrus, posterior division |
| 161 | 5.63 | -26 | 2 | -20 | L | Parahippocampal Gyrus, anterior division |
| | 5.41 | -30 | 4 | -24 | L | Temporal Pole |
| | 4.22 | -36 | 8 | -22 | L | Temporal Pole |
| 129 | 7.08 | 0 | -10 | -2 | C | Left Thalamus |
| | 3.5 | 4 | -4 | 4 | R | Right Thalamus |
| | 3.27 | 8 | -10 | 0 | R | Right Thalamus |
| 122 | 3.5 | 2 | 50 | 38 | R | Superior Frontal Gyrus |
| | 3.34 | 0 | 32 | 52 | C | Superior Frontal Gyrus |
| | 2.75 | 0 | 42 | 46 | C | Superior Frontal Gyrus |
| 118 | 3.36 | -58 | -64 | 14 | L | Lateral Occipital Cortex, superior division |
| | 2.74 | -62 | -62 | 8 | L | Middle Temporal Gyrus, temporooccipital part |

| Important Clusters from the PLSDA Model Classifying Fear (scenarios) | | | | | | |
|---|---|---|---|---|---|---|
| | 2.7 | -60 | -64 | -2 | L | Lateral Occipital Cortex, inferior division |
| 116 | 3.16 | -38 | -84 | 32 | L | Lateral Occipital Cortex, superior division |
| | 2.83 | -42 | -84 | 26 | L | Lateral Occipital Cortex, superior division |
| | 2.73 | -30 | -86 | 38 | L | Lateral Occipital Cortex, superior division |
| 96 | -7.75 | 2 | -46 | -42 | R | Right IX |
| | -5.58 | 2 | -50 | -52 | R | Right IX |
| | -3 | 4 | -42 | -38 | R | Brain-Stem |
| 86 | -3.53 | 2 | 38 | -18 | R | Frontal Medial Cortex |
| | -3.4 | 0 | 32 | -26 | C | Frontal Medial Cortex |
| | -2.95 | 2 | 32 | -14 | R | Subcallosal Cortex |
| 58 | 4.8 | 12 | -8 | -16 | R | Right Amygdala |
| | 3.22 | 18 | -2 | -14 | R | Right Amygdala |
| | 2.92 | 18 | -8 | -18 | R | Right Amygdala |
| 54 | -4.89 | 0 | 6 | -10 | C | Subcallosal Cortex |
| | -3.17 | 0 | 12 | -12 | C | Subcallosal Cortex |
| | -2.83 | -4 | 8 | -16 | L | Subcallosal Cortex |
| 53 | 3.17 | 58 | -22 | -8 | R | Middle Temporal Gyrus, posterior division |
| | 3.1 | 54 | -26 | -6 | R | Middle Temporal Gyrus, posterior division |
| | 2.49 | 56 | -32 | -2 | R | Middle Temporal Gyrus, posterior division |
| 52 | -3.3 | 4 | -58 | 10 | R | Precuneous Cortex |
| | -2.62 | -10 | -60 | 4 | L | Lingual Gyrus |
| | -2.56 | -4 | -66 | 8 | L | Intracalcarine Cortex |
| 50 | 3.82 | 32 | 8 | -20 | R | Temporal Pole |
| | 3.43 | 26 | 4 | -16 | R | Parahippocampal Gyrus, anterior division |
| 48 | -3.55 | -46 | 18 | -34 | L | Temporal Pole |
| | -2.84 | -44 | 12 | -42 | L | Temporal Pole |
| 44 | -6.62 | 2 | -14 | -24 | R | Brain-Stem |
| | -2.55 | 2 | -14 | -30 | R | Brain-Stem |
| 44 | -3.18 | 22 | 70 | 4 | R | Frontal Pole |
| | -2.78 | 24 | 66 | -4 | R | Frontal Pole |
| 42 | 3.45 | -48 | 22 | 10 | L | Inferior Frontal Gyrus, pars triangularis |
| 40 | 4.11 | -20 | -22 | -18 | L | Left Hippocampus |
| | 3.18 | -18 | -16 | -22 | L | Left Hippocampus |
| | 2.42 | -14 | -14 | -18 | L | Left Hippocampus |
| 40 | -3.33 | 48 | -38 | 62 | R | Postcentral Gyrus |

| Important Clusters from the PLSDA Model Classifying Fear (scenarios) | | | | | | |
|---|---|---|---|---|---|---|
| | -2.92 | 52 | -34 | 60 | R | Supramarginal Gyrus, anterior division / Supramarginal Gyrus, posterior division |
| | -2.2 | 46 | -36 | 56 | R | Supramarginal Gyrus, posterior division |
| 40 | 7.68 | -2 | -38 | -66 | L | Brain-Stem |
| | 7.3 | -4 | -42 | -70 | L | Brain-Stem |
| | 5.65 | 4 | -38 | -66 | R | Brain-Stem |
| 39 | 2.66 | 4 | -20 | 44 | R | Cingulate Gyrus, posterior division |
| | 2.58 | -2 | -22 | 44 | L | Cingulate Gyrus, posterior division |
| | 2.38 | 8 | -14 | 42 | R | Cingulate Gyrus, anterior division / Cingulate Gyrus, posterior division |
| 38 | 3.15 | -48 | -54 | 20 | L | Angular Gyrus |
| | 2.24 | -52 | -50 | 18 | L | Angular Gyrus |
| 37 | 3.15 | 48 | -76 | 26 | R | Lateral Occipital Cortex, superior division |
| 36 | -3.36 | -40 | -68 | 54 | L | Lateral Occipital Cortex, superior division |
| | -2.49 | -46 | -66 | 50 | L | Lateral Occipital Cortex, superior division |
| | -2.43 | -34 | -66 | 58 | L | Lateral Occipital Cortex, superior division |
| 35 | -6.94 | -14 | -32 | -16 | L | Parahippocampal Gyrus, posterior division |
| | -4.62 | -14 | -32 | -22 | L | Left I-IV |
| 33 | -2.8 | -10 | -50 | 30 | L | Cingulate Gyrus, posterior division |
| | -2.66 | -6 | -52 | 24 | L | Cingulate Gyrus, posterior division |
| | -2.29 | -2 | -54 | 28 | L | Cingulate Gyrus, posterior division |
| 33 | 2.48 | 62 | -58 | 12 | R | Middle Temporal Gyrus, temporooccipital part |
| | 2.31 | 64 | -52 | 14 | R | Angular Gyrus |
| | 2.3 | 62 | -58 | 6 | R | Middle Temporal Gyrus, temporooccipital part |
| 33 | 3.81 | 18 | -46 | 2 | R | Cingulate Gyrus, posterior division |
| | 2.61 | 20 | -54 | 4 | R | Lingual Gyrus |
| 32 | 3.59 | -42 | -22 | 12 | L | Heschl's Gyrus (includes H1 and H2) |
| | 2.88 | -40 | -30 | 12 | L | Planum Temporale |
| 32 | 3.05 | 52 | 40 | -8 | R | Frontal Pole |
| | 2.86 | 52 | 34 | -8 | R | Frontal Pole |
| 31 | -2.97 | 32 | -72 | 38 | R | Lateral Occipital Cortex, superior division |
| 30 | -2.73 | 46 | 52 | 16 | R | Frontal Pole |
| | -2.57 | 38 | 60 | 18 | R | Frontal Pole |
| | -2.22 | 44 | 50 | 22 | R | Frontal Pole |
| 30 | -3.34 | 18 | -60 | 72 | R | Lateral Occipital Cortex, superior division |
| | -2.7 | 24 | -58 | 70 | R | Lateral Occipital Cortex, superior division |

| **Important Clusters from the PLSDA Model Classifying Fear (scenarios)** | | | | | | |
|---|---|---|---|---|---|---|
| | -2.39 | 10 | -56 | 74 | R | Lateral Occipital Cortex, superior division |

*Horror*

| Important Clusters from the PLSDA Model Classifying Horror (scenarios) | | | | | | |
|---|---|---|---|---|---|---|
| Cluster Size (Voxels) | Max Coef | MNI | | | Hemisphere | Anatomical Label |
| | | x | y | z | | |
| 415 | 4.45 | -64 | -40 | 32 | L | Supramarginal Gyrus, anterior division |
| | 4.19 | -66 | -34 | 32 | L | Supramarginal Gyrus, anterior division |
| | 3.59 | -56 | -36 | 36 | L | Supramarginal Gyrus, anterior division |
| 306 | -4.62 | 0 | -48 | 24 | C | Cingulate Gyrus, posterior division |
| | -3.95 | 2 | -40 | 24 | R | Cingulate Gyrus, posterior division |
| | -3.8 | 0 | -48 | 32 | C | Cingulate Gyrus, posterior division |
| 242 | 3.66 | -40 | -84 | 28 | L | Lateral Occipital Cortex, superior division |
| | 3.43 | -36 | -84 | 36 | L | Lateral Occipital Cortex, superior division |
| | 3.24 | -34 | -88 | 30 | L | Lateral Occipital Cortex, superior division |
| 163 | -3.94 | -44 | 16 | -34 | L | Temporal Pole |
| | -3.81 | -52 | 14 | -24 | L | Temporal Pole |
| | -2.82 | -48 | 10 | -26 | L | Temporal Pole |
| 161 | 4.15 | 2 | 46 | 46 | R | Superior Frontal Gyrus |
| | 3.73 | 4 | 46 | 32 | R | Paracingulate Gyrus |
| | 3.18 | -2 | 54 | 24 | L | Superior Frontal Gyrus |
| 158 | 4.36 | -24 | 0 | -18 | L | Left Amygdala |
| | 4.23 | -12 | -2 | -22 | L | Parahippocampal Gyrus, anterior division |
| | 4.1 | -22 | -2 | -24 | L | Left Amygdala |
| 142 | 3.58 | -58 | -60 | 2 | L | Middle Temporal Gyrus, temporooccipital part |
| | 3.24 | -56 | -64 | -2 | L | Lateral Occipital Cortex, inferior division |
| | 2.98 | -58 | -62 | -8 | L | Lateral Occipital Cortex, inferior division |
| 140 | 3.55 | 50 | -44 | 32 | R | Angular Gyrus |
| | 3.3 | 64 | -44 | 34 | R | Supramarginal Gyrus, posterior division |
| | 3.29 | 64 | -50 | 26 | R | Angular Gyrus |
| 130 | 4.03 | 54 | -26 | -4 | R | Superior Temporal Gyrus, posterior division |
| | 3.16 | 60 | -18 | -10 | R | Middle Temporal Gyrus, posterior division |
| | 3.06 | 62 | -28 | -4 | R | Middle Temporal Gyrus, posterior division |
| 108 | 3.75 | -2 | -38 | 46 | L | Cingulate Gyrus, posterior division |
| | 2.83 | -6 | -44 | 56 | L | Precuneous Cortex |
| | 2.55 | -6 | -46 | 64 | L | Precuneous Cortex |
| 85 | -3.63 | 34 | 16 | 8 | R | Insular Cortex |
| | -3.33 | 34 | 22 | 8 | R | Frontal Operculum Cortex |

| Important Clusters from the PLSDA Model Classifying Horror (scenarios) | | | | | | |
|---|---|---|---|---|---|---|
| | -2.39 | 32 | 26 | 2 | R | Insular Cortex |
| 72 | 3.93 | -2 | 0 | 36 | L | Cingulate Gyrus, anterior division |
| | 3.28 | 0 | 6 | 32 | C | Cingulate Gyrus, anterior division |
| | 2.15 | 2 | -8 | 42 | R | Cingulate Gyrus, anterior division |
| 68 | -3.49 | 46 | 16 | -32 | R | Temporal Pole |
| | -2.76 | 36 | 18 | -38 | R | Temporal Pole |
| 58 | 4.27 | -32 | -36 | -16 | L | Temporal Fusiform Cortex, posterior division |
| | 2.78 | -30 | -30 | -18 | L | Parahippocampal Gyrus, posterior division |
| | 2.5 | -36 | -26 | -22 | L | Temporal Fusiform Cortex, posterior division |
| 57 | -3.81 | -54 | 12 | -12 | L | Temporal Pole |
| | -3.07 | -46 | 2 | -12 | L | Planum Polare |
| 52 | -4.13 | 42 | 16 | -16 | R | Temporal Pole |
| | -3.77 | 42 | 10 | -14 | R | Insular Cortex |
| 51 | -2.78 | -2 | -62 | 28 | L | Precuneous Cortex |
| | -2.76 | -4 | -70 | 26 | L | Precuneous Cortex |
| | -2.09 | 0 | -68 | 30 | C | Precuneous Cortex |
| 50 | -3.91 | 14 | 2 | -26 | R | Parahippocampal Gyrus, anterior division |
| | -3.47 | 18 | 6 | -22 | R | Frontal Orbital Cortex |
| | -3.34 | 28 | 12 | -26 | R | Temporal Pole |
| 50 | 3.67 | -42 | -40 | -20 | L | Temporal Fusiform Cortex, posterior division |
| | 2.85 | -46 | -46 | -20 | L | Inferior Temporal Gyrus, temporooccipital part |
| | 2.84 | -48 | -40 | -20 | L | Inferior Temporal Gyrus, posterior division |
| 50 | 3.96 | -22 | 0 | -40 | L | Parahippocampal Gyrus, anterior division |
| | 3.06 | -30 | 0 | -36 | L | Parahippocampal Gyrus, anterior division |
| | 2.66 | -28 | -4 | -32 | L | Parahippocampal Gyrus, anterior division |
| 50 | 4.89 | 20 | -2 | -16 | R | Right Amygdala |
| | 2.96 | 16 | -6 | -14 | R | Right Amygdala |
| 45 | 4.73 | -10 | -36 | -6 | L | Parahippocampal Gyrus, posterior division |
| | 4.69 | -14 | -32 | -12 | L | Parahippocampal Gyrus, posterior division |
| | 2.64 | -14 | -32 | -2 | L | Left Thalamus |
| 43 | 3.88 | -12 | -30 | 38 | L | Cingulate Gyrus, posterior division |
| | 2.98 | -8 | -26 | 40 | L | Cingulate Gyrus, posterior division |
| | 2.97 | -10 | -20 | 38 | L | Cingulate Gyrus, posterior division |
| 42 | 2.88 | -46 | 4 | 22 | L | Precentral Gyrus |
| | 2.85 | -50 | 8 | 18 | L | Inferior Frontal Gyrus, pars opercularis |

| Important Clusters from the PLSDA Model Classifying Horror (scenarios) | | | | | | |
|---|---|---|---|---|---|---|
| 41 | -3.9 | 62 | -16 | 16 | R | Central Opercular Cortex |
| | -2.08 | 56 | -16 | 14 | R | Central Opercular Cortex |
| 41 | 3.19 | -50 | 40 | 14 | L | Frontal Pole |
| | 3.07 | -46 | 38 | 10 | L | Frontal Pole |
| 39 | -2.88 | -2 | 36 | -16 | L | Frontal Medial Cortex |
| | -2.63 | -4 | 46 | -16 | L | Frontal Medial Cortex |
| 37 | -3.08 | -22 | -90 | -20 | L | Occipital Fusiform Gyrus |
| | -2.74 | -28 | -84 | -20 | L | Occipital Fusiform Gyrus |
| 35 | -2.87 | 2 | 22 | 48 | R | Paracingulate Gyrus |
| | -2.63 | 8 | 16 | 48 | R | Paracingulate Gyrus |
| 34 | 3.1 | 28 | 0 | -36 | R | Parahippocampal Gyrus, anterior division |
| | 2.97 | 18 | -2 | -36 | R | Parahippocampal Gyrus, anterior division |
| 34 | 3.24 | 16 | -46 | -50 | R | Right VIIIb |
| | 3.14 | 12 | -48 | -42 | R | Right IX |
| | 2.4 | 22 | -40 | -48 | R | Right X |
| 33 | -4.89 | -8 | -50 | -8 | L | Left V |
| 33 | -3.22 | -10 | 72 | 10 | L | Frontal Pole |
| | -3.09 | -4 | 72 | 10 | L | Frontal Pole |
| 32 | 3.04 | -32 | -28 | -28 | L | Temporal Fusiform Cortex, posterior division |
| | 2.89 | -32 | -20 | -26 | L | Parahippocampal Gyrus, anterior division |
| | 2.23 | -26 | -28 | -30 | L | Parahippocampal Gyrus, posterior division |
| 32 | -3.41 | 24 | 24 | -22 | R | Frontal Orbital Cortex |
| | -3.34 | 16 | 24 | -26 | R | Frontal Orbital Cortex |
| 32 | 3.91 | -4 | -14 | 0 | L | Left Thalamus |
| 31 | 4.9 | 2 | -18 | -26 | R | Brain-Stem |
| | 2.35 | -2 | -24 | -34 | L | Brain-Stem |
| | 2.13 | 0 | -28 | -30 | C | Brain-Stem |
| 30 | -3.01 | -46 | 34 | -16 | L | Frontal Orbital Cortex |

*Joy*

| Important Clusters from the PLSDA Model Classifying Joy (scenarios) | | | | | | |
|---|---|---|---|---|---|---|
| Cluster Size (Voxels) | Max Coef | MNI | | | Hemisphere | Anatomical Label |
| | | x | y | z | | |
| 764 | -4.36 | -54 | -40 | 2 | L | Superior Temporal Gyrus, posterior division |
| | -4.26 | -50 | -44 | 4 | L | Middle Temporal Gyrus, temporooccipital part |
| | -3.56 | -56 | -52 | 10 | L | Middle Temporal Gyrus, temporooccipital part |
| 604 | 4.68 | 2 | -58 | 28 | R | Precuneous Cortex |
| | 3.78 | 0 | -62 | 34 | C | Precuneous Cortex |
| | 3.74 | -4 | -54 | 28 | L | Cingulate Gyrus, posterior division |
| 372 | 4.11 | -2 | 62 | 4 | L | Frontal Pole |
| | 3.77 | 0 | 64 | -4 | C | Frontal Pole |
| | 3.63 | 0 | 60 | 12 | C | Frontal Pole |
| 354 | -4.7 | 46 | -30 | -2 | R | Superior Temporal Gyrus, posterior division |
| | -4.29 | 50 | -16 | -10 | R | Superior Temporal Gyrus, posterior division |
| | -3.94 | 56 | -10 | -8 | R | Superior Temporal Gyrus, posterior division |
| 275 | -4.02 | 48 | 20 | 30 | R | Middle Frontal Gyrus |
| | -4.01 | 36 | 8 | 30 | R | Precentral Gyrus |
| | -3.4 | 42 | 10 | 24 | R | Inferior Frontal Gyrus, pars opercularis |
| 268 | -3.91 | -40 | 4 | 36 | L | Precentral Gyrus |
| | -3.33 | -42 | 6 | 24 | L | Precentral Gyrus |
| | -3.28 | -48 | 12 | 30 | L | Middle Frontal Gyrus |
| 236 | -3.29 | -48 | -58 | -18 | L | Inferior Temporal Gyrus, temporooccipital part |
| | -3.15 | -44 | -52 | -14 | L | Inferior Temporal Gyrus, temporooccipital part |
| | -2.95 | -54 | -54 | -24 | L | Inferior Temporal Gyrus, temporooccipital part |
| 176 | 4.74 | 42 | -14 | 0 | R | Insular Cortex |
| | 3.65 | 44 | -8 | 2 | R | Insular Cortex |
| | 3.07 | 46 | -10 | 8 | R | Central Opercular Cortex |
| 174 | 3.64 | 62 | -28 | 28 | R | Supramarginal Gyrus, anterior division |
| | 3.3 | 54 | -24 | 16 | R | Parietal Operculum Cortex |
| | 2.86 | 50 | -28 | 20 | R | Parietal Operculum Cortex |
| 172 | 4.05 | 0 | 34 | 22 | C | Cingulate Gyrus, anterior division |
| | 3.59 | 2 | 36 | 12 | R | Cingulate Gyrus, anterior division |
| | 3.12 | 0 | 24 | 24 | C | Cingulate Gyrus, anterior division |
| 138 | -3.07 | -36 | -68 | -18 | L | Occipital Fusiform Gyrus |
| | -2.98 | -38 | -76 | -18 | L | Occipital Fusiform Gyrus |

| Important Clusters from the PLSDA Model Classifying Joy (scenarios) | | | | | | |
|---|---|---|---|---|---|---|
| | -2.91 | -36 | -82 | -20 | L | Lateral Occipital Cortex, inferior division / Occipital Fusiform Gyrus |
| 99 | -4.82 | -4 | -14 | -28 | L | Brain-Stem |
| | -3.72 | 6 | -14 | -26 | R | Brain-Stem |
| | -3.5 | 14 | -20 | -24 | R | Brain-Stem |
| 97 | 3.79 | -36 | 12 | 2 | L | Insular Cortex |
| | 3.75 | -40 | 6 | 2 | L | Insular Cortex |
| | 2.49 | -36 | 8 | 10 | L | Central Opercular Cortex |
| 96 | -3.6 | -8 | -82 | 4 | L | Intracalcarine Cortex |
| | -3.22 | -6 | -74 | 12 | L | Intracalcarine Cortex |
| | -3.01 | -12 | -74 | 10 | L | Intracalcarine Cortex |
| 92 | -3.43 | -48 | 2 | 56 | L | Middle Frontal Gyrus |
| | -3.01 | -52 | 2 | 50 | L | Precentral Gyrus |
| | -2.97 | -54 | 6 | 44 | L | Precentral Gyrus |
| 85 | -3.04 | 14 | -72 | 10 | R | Intracalcarine Cortex |
| | -3 | 10 | -74 | 14 | R | Intracalcarine Cortex |
| | -2.96 | 12 | -82 | 4 | R | Intracalcarine Cortex |
| 80 | -6.34 | 8 | -92 | -14 | R | Lingual Gyrus / Occipital Pole |
| | -3.46 | 14 | -94 | -14 | R | Occipital Pole |
| 73 | 3.19 | 14 | -66 | 34 | R | Precuneous Cortex |
| | 2.8 | 18 | -66 | 28 | R | Precuneous Cortex |
| | 2.34 | 20 | -58 | 24 | R | Precuneous Cortex |
| 71 | -4.76 | 8 | -34 | 4 | R | Right Thalamus |
| | -3.81 | -2 | -32 | -2 | L | Brain-Stem |
| | -3.03 | 10 | -30 | -2 | R | Right Thalamus |
| 71 | -3.1 | 2 | 42 | 44 | R | Superior Frontal Gyrus |
| | -2.7 | 2 | 50 | 40 | R | Superior Frontal Gyrus |
| 67 | -4.44 | -32 | -46 | -22 | L | Temporal Occipital Fusiform Cortex |
| | -3.35 | -24 | -40 | -14 | L | Parahippocampal Gyrus, posterior division |
| | -3.08 | -30 | -34 | -18 | L | Temporal Fusiform Cortex, posterior division |
| 65 | 3.34 | 0 | -26 | 30 | C | Cingulate Gyrus, posterior division |
| | 3.24 | 0 | -38 | 26 | C | Cingulate Gyrus, posterior division |
| | 2.41 | -4 | -42 | 24 | L | Cingulate Gyrus, posterior division |
| 62 | -3.09 | 58 | 6 | -14 | R | Temporal Pole |
| | -2.55 | 52 | 0 | -16 | R | Superior Temporal Gyrus, anterior division |

| Important Clusters from the PLSDA Model Classifying Joy (scenarios) | | | | | | |
|---|---|---|---|---|---|---|
| | -2.42 | 58 | 0 | -10 | R | Superior Temporal Gyrus, anterior division |
| 61 | -3.02 | 56 | 28 | 4 | R | Inferior Frontal Gyrus, pars triangularis |
| | -2.86 | 48 | 32 | 18 | R | Inferior Frontal Gyrus, pars triangularis |
| | -2.42 | 56 | 34 | 18 | R | Frontal Pole / Inferior Frontal Gyrus, pars triangularis |
| 60 | -5.67 | 20 | -26 | -26 | R | Parahippocampal Gyrus, posterior division |
| | -3.38 | 16 | -16 | -26 | R | Parahippocampal Gyrus, anterior division |
| | -2.6 | 24 | -20 | -20 | R | Right Hippocampus |
| 56 | -4.09 | -28 | 16 | -26 | L | Frontal Orbital Cortex |
| | -3.62 | -24 | 10 | -26 | L | Frontal Orbital Cortex |
| 54 | 4.76 | 28 | 8 | -18 | R | Frontal Orbital Cortex |
| | 4.03 | 40 | 14 | -18 | R | Temporal Pole |
| | 2.78 | 42 | 12 | -12 | R | Insular Cortex |
| 50 | -3.49 | -2 | -96 | -14 | L | Occipital Pole |
| | -2.41 | -10 | -94 | -14 | L | Occipital Pole |
| | -2.34 | -4 | -94 | -6 | L | Occipital Pole |
| 49 | 3 | -40 | 48 | 26 | L | Frontal Pole |
| 45 | 4.71 | -44 | -14 | -2 | L | Planum Polare |
| 43 | -2.81 | 52 | 16 | -24 | R | Temporal Pole |
| | -2.76 | 54 | 14 | -18 | R | Temporal Pole |
| | -2.52 | 50 | 12 | -28 | R | Temporal Pole |
| 43 | 2.78 | -38 | 58 | -4 | L | Frontal Pole |
| | 2.77 | -32 | 62 | -4 | L | Frontal Pole |
| | 2.63 | -26 | 58 | 0 | L | Frontal Pole |
| 42 | -5.97 | -18 | -26 | -24 | L | Parahippocampal Gyrus, posterior division |
| | -3.08 | -22 | -22 | -30 | L | Parahippocampal Gyrus, anterior division |
| | -2.66 | -14 | -20 | -26 | L | Brain-Stem |
| 41 | -4.03 | 20 | -54 | 2 | R | Lingual Gyrus |
| | -3.29 | 12 | -56 | 6 | R | Precuneous Cortex |
| 40 | -2.68 | -54 | 4 | -16 | L | Temporal Pole |
| | -2.36 | -54 | -4 | -12 | L | Superior Temporal Gyrus, anterior division |
| 36 | -4.47 | 18 | -40 | -48 | R | Right X |
| 36 | 2.86 | 62 | 4 | 4 | R | Planum Polare |
| | 2.64 | 62 | 10 | 4 | R | Precentral Gyrus |
| | 2.17 | 56 | 8 | 0 | R | Temporal Pole |

| Important Clusters from the PLSDA Model Classifying Joy (scenarios) | | | | | | |
|---|---|---|---|---|---|---|
| 35 | 2.86 | -44 | 52 | 6 | L | Frontal Pole |
| 35 | 3.24 | 36 | 4 | 10 | R | Insular Cortex |
| | 2.46 | 42 | 2 | 12 | R | Central Opercular Cortex |
| 32 | 3.24 | -56 | -2 | 6 | L | Central Opercular Cortex |
| | 2.03 | -52 | 0 | 2 | L | Central Opercular Cortex |
| 32 | -3.19 | -12 | -48 | 2 | L | Cingulate Gyrus, posterior division |
| | -2.88 | -16 | -50 | -2 | L | Lingual Gyrus |
| 32 | -2.54 | -28 | -72 | 28 | L | Lateral Occipital Cortex, superior division |
| | -2.5 | -28 | -82 | 24 | L | Lateral Occipital Cortex, superior division |
| 31 | -3.64 | -18 | -38 | -46 | L | Left X |
| | -2.61 | -16 | -44 | -50 | L | Left VIIIb |

*Neutral*

| Important Clusters from the PLSDA Model Classifying Neutral (scenarios) | | | | | | |
|---|---|---|---|---|---|---|
| Cluster Size (Voxels) | Max Coef | MNI | | | Hemisphere | Anatomical Label |
| | | x | y | z | | |
| 1475 | 4.09 | 50 | 44 | -12 | R | Frontal Pole |
| | 4 | 48 | 50 | -6 | R | Frontal Pole |
| | 3.9 | 48 | 42 | 16 | R | Frontal Pole |
| 862 | -4.11 | 8 | -92 | -10 | R | Occipital Pole |
| | -4.02 | -10 | -80 | 10 | L | Intracalcarine Cortex |
| | -3.86 | 8 | -84 | 2 | R | Intracalcarine Cortex |
| 652 | 4.45 | 46 | -54 | 60 | R | Angular Gyrus |
| | 4.01 | 52 | -46 | 56 | R | Angular Gyrus |
| | 3.96 | 48 | -48 | 60 | R | Angular Gyrus |
| 348 | -4.04 | -56 | -4 | -10 | L | Superior Temporal Gyrus, anterior division |
| | -4.03 | -58 | 4 | -10 | L | Temporal Pole |
| | -3.73 | -58 | -8 | -6 | L | Superior Temporal Gyrus, anterior division |
| 266 | -4.72 | -20 | -14 | -14 | L | Left Amygdala |
| | -4.5 | -20 | -2 | -16 | L | Left Amygdala |
| | -4.06 | -12 | -8 | -20 | L | Left Hippocampus |
| 232 | 3.49 | -46 | -50 | 58 | L | Supramarginal Gyrus, posterior division |
| | 3.34 | -48 | -44 | 60 | L | Supramarginal Gyrus, posterior division |
| | 3.02 | -52 | -40 | 56 | L | Supramarginal Gyrus, posterior division |
| 221 | -4.08 | 58 | -8 | -8 | R | Superior Temporal Gyrus, posterior division |
| | -3.14 | 60 | 6 | -12 | R | Temporal Pole |
| | -2.99 | 58 | 0 | -14 | R | Superior Temporal Gyrus, anterior division |
| 189 | -3.06 | -40 | -86 | -16 | L | Lateral Occipital Cortex, inferior division |
| | -3.05 | -34 | -94 | -14 | L | Occipital Pole |
| | -3.02 | -40 | -74 | -8 | L | Lateral Occipital Cortex, inferior division |
| 170 | 3.3 | 2 | 36 | 38 | R | Paracingulate Gyrus |
| | 2.87 | 2 | 20 | 50 | R | Paracingulate Gyrus |
| | 2.76 | 4 | 28 | 46 | R | Paracingulate Gyrus |
| 162 | -3.55 | 12 | -60 | 10 | R | Precuneous Cortex |
| | -3.32 | -4 | -52 | 18 | L | Cingulate Gyrus, posterior division |
| | -3.06 | 2 | -52 | 20 | R | Cingulate Gyrus, posterior division |
| 119 | -3.57 | -2 | 64 | 4 | L | Frontal Pole |
| | -3.39 | -4 | 70 | 12 | L | Frontal Pole |

| Important Clusters from the PLSDA Model Classifying Neutral (scenarios) | | | | | | |
|---|---|---|---|---|---|---|
| | -3.32 | -2 | 64 | 14 | L | Frontal Pole |
| 97 | -4.62 | 48 | -68 | -20 | R | Lateral Occipital Cortex, inferior division |
| | -2.81 | 44 | -84 | -10 | R | Lateral Occipital Cortex, inferior division |
| | -2.66 | 46 | -74 | -14 | R | Lateral Occipital Cortex, inferior division |
| 95 | 3.32 | 28 | 68 | -6 | R | Frontal Pole |
| | 3.27 | 34 | 66 | -2 | R | Frontal Pole |
| | 2.87 | 34 | 66 | -8 | R | Frontal Pole |
| 80 | -3.39 | -40 | -48 | -24 | L | Temporal Occipital Fusiform Cortex |
| | -2.8 | -36 | -40 | -26 | L | Temporal Fusiform Cortex, posterior division |
| | -2.74 | -42 | -52 | -20 | L | Temporal Occipital Fusiform Cortex |
| 79 | -3.73 | -50 | -10 | 56 | L | Precentral Gyrus |
| | -3.1 | -46 | -12 | 60 | L | Precentral Gyrus |
| | -2.64 | -54 | -8 | 52 | L | Precentral Gyrus |
| 73 | 3.58 | -38 | 52 | -12 | L | Frontal Pole |
| | 2.78 | -34 | 48 | -16 | L | Frontal Pole |
| | 2.37 | -44 | 50 | -12 | L | Frontal Pole |
| 65 | 2.78 | 34 | 40 | -14 | R | Frontal Pole |
| | 2.54 | 26 | 40 | -18 | R | Frontal Pole |
| | 2.49 | 18 | 42 | -20 | R | Frontal Pole |
| 63 | -3.05 | -48 | -40 | 2 | L | Superior Temporal Gyrus, posterior division |
| | -3 | -50 | -34 | 0 | L | Superior Temporal Gyrus, posterior division |
| | -2.95 | -50 | -46 | 6 | L | Middle Temporal Gyrus, temporooccipital part |
| 56 | 3.53 | 44 | 22 | -6 | R | Frontal Orbital Cortex |
| | 2.63 | 54 | 20 | -2 | R | Inferior Frontal Gyrus, pars triangularis |
| | 2.29 | 44 | 22 | -12 | R | Frontal Orbital Cortex |
| 50 | -4.07 | -10 | -32 | -4 | L | Left Thalamus |
| | -3.64 | -10 | -38 | -4 | L | Parahippocampal Gyrus, posterior division |
| | -2.49 | -12 | -42 | 4 | L | Left Hippocampus |
| 50 | -2.75 | 2 | 4 | 52 | R | Juxtapositional Lobule Cortex (formerly Supplementary Motor Cortex) |
| | -2.71 | 0 | -2 | 54 | C | Juxtapositional Lobule Cortex (formerly Supplementary Motor Cortex) |
| | -2.45 | -2 | -8 | 46 | L | Cingulate Gyrus, anterior division |
| 46 | 3.04 | -46 | 48 | -4 | L | Frontal Pole |
| | 2.55 | -46 | 44 | -10 | L | Frontal Pole |
| | 2.06 | -48 | 40 | -14 | L | Frontal Pole |

| Important Clusters from the PLSDA Model Classifying Neutral (scenarios) | | | | | | |
|---|---|---|---|---|---|---|
| 41 | -3.77 | -14 | -54 | 4 | L | Precuneous Cortex |
| | -2.18 | -16 | -50 | -2 | L | Lingual Gyrus |
| 40 | -3.12 | -6 | -14 | 4 | L | Left Thalamus |
| | -2.63 | 0 | -12 | 10 | C | Left Thalamus |
| | -2.24 | -6 | -14 | 10 | L | Left Thalamus |
| 38 | -3.77 | 24 | -20 | -14 | R | Right Hippocampus |
| | -2.46 | 28 | -26 | -14 | R | Right Hippocampus |
| 37 | -3.3 | 34 | -46 | -22 | R | Temporal Occipital Fusiform Cortex |
| | -2.24 | 40 | -46 | -18 | R | Temporal Occipital Fusiform Cortex |
| 37 | -2.9 | -50 | -64 | 16 | L | Lateral Occipital Cortex, superior division |
| | -2.65 | -52 | -60 | 10 | L | Middle Temporal Gyrus, temporooccipital part |
| | -2.45 | -40 | -62 | 18 | L | Lateral Occipital Cortex, superior division |
| 35 | -4.18 | 38 | 10 | -18 | R | Temporal Pole |
| 35 | -3.1 | 14 | -50 | 4 | R | Cingulate Gyrus, posterior division |
| 34 | -5.1 | 12 | -34 | -2 | R | Right Thalamus |
| | -2.29 | 10 | -40 | 2 | R | Cingulate Gyrus, posterior division |
| 34 | -3.59 | -32 | -36 | -16 | L | Temporal Fusiform Cortex, posterior division |
| | -3.38 | -24 | -36 | -16 | L | Parahippocampal Gyrus, posterior division |
| 33 | -3.15 | 0 | -62 | 36 | C | Precuneous Cortex |
| 32 | -2.48 | 48 | -28 | -8 | R | Middle Temporal Gyrus, posterior division |
| | -2.43 | 48 | -32 | -2 | R | Middle Temporal Gyrus, posterior division |
| 30 | -2.71 | -38 | -68 | -22 | L | Left Crus I |
| | -2.69 | -38 | -74 | -20 | L | Occipital Fusiform Gyrus |

*Romance*

| Important Clusters from the PLSDA Model Classifying Romance (scenarios) | | | | | | |
|---|---|---|---|---|---|---|
| **Cluster Size (Voxels)** | **Max Coef** | **MNI** | | | **Hemisphere** | **Anatomical Label** |
| | | **x** | **y** | **z** | | |
| 209 | 4.11 | 4 | 30 | -8 | R | Subcallosal Cortex |
| | 3.78 | 0 | 38 | -6 | C | Cingulate Gyrus, anterior division |
| | 3.63 | -2 | 30 | -2 | L | Subcallosal Cortex |
| 193 | -4.51 | 14 | -54 | 16 | R | Precuneous Cortex |
| | -4.35 | 6 | -54 | 6 | R | Precuneous Cortex |
| | -3.73 | 8 | -48 | -2 | R | Lingual Gyrus |
| 188 | -4.46 | -20 | -58 | -12 | L | Lingual Gyrus |
| | -3.29 | -22 | -72 | -18 | L | Left VI |
| | -3.28 | -20 | -66 | -16 | L | Left VI |
| 164 | 6.14 | 2 | -20 | 0 | R | Right Thalamus |
| | 5.17 | -4 | -32 | -4 | L | Brain-Stem |
| | 4.07 | 0 | -34 | -16 | C | Brain-Stem |
| 129 | 3.47 | 12 | -74 | 12 | R | Intracalcarine Cortex |
| | 3.4 | 16 | -62 | 6 | R | Intracalcarine Cortex |
| | 3.17 | 20 | -68 | 6 | R | Intracalcarine Cortex |
| 115 | -3.33 | 0 | 36 | 48 | C | Superior Frontal Gyrus |
| | -2.93 | 0 | 36 | 42 | C | Superior Frontal Gyrus |
| | -2.84 | 2 | 26 | 54 | R | Superior Frontal Gyrus |
| 107 | 3.85 | -50 | 2 | 52 | L | Precentral Gyrus |
| | 3.6 | -54 | 6 | 46 | L | Precentral Gyrus |
| | 3.42 | -46 | 2 | 58 | L | Middle Frontal Gyrus |
| 98 | 3.46 | -6 | -94 | -10 | L | Occipital Pole |
| | 3.15 | -8 | -84 | 4 | L | Intracalcarine Cortex |
| | 2.76 | -10 | -92 | -2 | L | Occipital Pole |
| 92 | -3.85 | -2 | -44 | 40 | L | Cingulate Gyrus, posterior division |
| | -3.73 | 0 | -60 | 52 | C | Precuneous Cortex |
| | -2.65 | 2 | -50 | 54 | R | Precuneous Cortex |
| 84 | 4.88 | -56 | -50 | 8 | L | Middle Temporal Gyrus, temporooccipital part |
| | 2.33 | -58 | -42 | 8 | L | Superior Temporal Gyrus, posterior division |
| 79 | -4.28 | -16 | -58 | 18 | L | Precuneous Cortex |
| | -3.59 | -10 | -58 | 14 | L | Precuneous Cortex |
| | -3.07 | -4 | -60 | 8 | L | Precuneous Cortex |

| Important Clusters from the PLSDA Model Classifying Romance (scenarios) | | | | | | |
|---|---|---|---|---|---|---|
| 66 | -3.84 | 0 | -72 | 2 | C | Lingual Gyrus |
| | -3.05 | 0 | -78 | -4 | C | Lingual Gyrus |
| | -2.58 | -6 | -76 | -8 | L | Lingual Gyrus |
| 61 | 4 | -36 | -62 | 62 | L | Lateral Occipital Cortex, superior division |
| | 3.79 | -42 | -58 | 60 | L | Lateral Occipital Cortex, superior division |
| | 2.73 | -48 | -56 | 56 | L | Angular Gyrus |
| 60 | 3.32 | 46 | -36 | 4 | R | Superior Temporal Gyrus, posterior division |
| | 3.06 | 50 | -38 | 10 | R | Supramarginal Gyrus, posterior division |
| | 2.98 | 56 | -40 | 8 | R | Supramarginal Gyrus, posterior division |
| 58 | -3.16 | 24 | -96 | -2 | R | Occipital Pole |
| | -2.9 | 30 | -96 | -8 | R | Occipital Pole |
| | -2.81 | 20 | -100 | -6 | R | Occipital Pole |
| 57 | 3.57 | -2 | 60 | 4 | L | Frontal Pole |
| | 3.12 | -2 | 66 | 4 | L | Frontal Pole |
| | 2.47 | 0 | 64 | -4 | C | Frontal Pole |
| 54 | -3.91 | -4 | -48 | 6 | L | Cingulate Gyrus, posterior division |
| | -3.91 | -10 | -54 | 4 | L | Precuneous Cortex |
| | -3.2 | -6 | -44 | 2 | L | Cingulate Gyrus, posterior division |
| 53 | 3.04 | 42 | 8 | 24 | R | Inferior Frontal Gyrus, pars opercularis |
| | 2.72 | 48 | 14 | 26 | R | Inferior Frontal Gyrus, pars opercularis |
| | 2.46 | 40 | 6 | 30 | R | Precentral Gyrus |
| 53 | 3.32 | 30 | 14 | -20 | R | Frontal Orbital Cortex |
| | 2.76 | 28 | 12 | -14 | R | Insular Cortex |
| | 2.67 | 26 | 8 | -20 | R | Frontal Orbital Cortex |
| 49 | 3.5 | -38 | -58 | -22 | L | Temporal Occipital Fusiform Cortex |
| | 3.49 | -30 | -56 | -18 | L | Temporal Occipital Fusiform Cortex |
| 49 | 3.07 | -58 | -38 | 4 | L | Superior Temporal Gyrus, posterior division |
| | 2.75 | -52 | -30 | 0 | L | Superior Temporal Gyrus, posterior division |
| | 2.69 | -52 | -40 | 0 | L | Middle Temporal Gyrus, posterior division |
| 48 | 4.05 | 10 | -84 | 4 | R | Intracalcarine Cortex |
| 47 | 3.83 | -6 | -72 | 12 | L | Intracalcarine Cortex |
| | 2.8 | -8 | -78 | 14 | L | Intracalcarine Cortex |
| | 2.74 | -10 | -68 | 10 | L | Intracalcarine Cortex |
| 46 | 6.84 | 12 | -44 | -60 | R | Right VIIIb |
| | 2.84 | 6 | -44 | -54 | R | Brain-Stem |

| Important Clusters from the PLSDA Model Classifying Romance (scenarios) | | | | | | |
|---|---|---|---|---|---|---|
| | 2.59 | 6 | -46 | -62 | R | Brain-Stem |
| 44 | -3.74 | -42 | 2 | -14 | L | Planum Polare |
| | -3.26 | -38 | 6 | -16 | L | Insular Cortex |
| | -2.8 | -42 | 2 | -20 | L | Planum Polare |
| 44 | 3.16 | 14 | -36 | -12 | R | Parahippocampal Gyrus, posterior division |
| | 2.78 | 10 | -42 | -12 | R | Right I-IV |
| | 2.63 | 12 | -40 | -6 | R | Lingual Gyrus |
| 44 | 3.27 | 0 | -72 | 36 | C | Precuneous Cortex |
| | 2.38 | 2 | -74 | 42 | R | Precuneous Cortex |
| | 2.25 | 0 | -70 | 46 | C | Precuneous Cortex |
| 42 | 6.58 | -2 | 6 | -14 | L | Subcallosal Cortex |
| | 3.06 | 0 | 12 | -8 | C | Subcallosal Cortex |
| 41 | 4.82 | 6 | -26 | 10 | R | Right Thalamus |
| | 3.26 | 4 | -20 | 14 | R | Right Thalamus |
| 41 | -3.82 | 48 | -2 | 4 | R | Central Opercular Cortex |
| | -3.54 | 52 | -6 | 2 | R | Heschl's Gyrus (includes H1 and H2) |
| 40 | 4.8 | 0 | 36 | 12 | C | Cingulate Gyrus, anterior division |
| | 2.43 | 6 | 32 | 12 | R | Cingulate Gyrus, anterior division |
| 40 | 3.19 | 0 | -14 | 32 | C | Cingulate Gyrus, anterior division |
| | 2.94 | 0 | -20 | 34 | C | Cingulate Gyrus, posterior division |
| 40 | 3.58 | 0 | -36 | 24 | C | Cingulate Gyrus, posterior division |
| | 3.18 | 0 | -42 | 26 | C | Cingulate Gyrus, posterior division |
| | 2.68 | 4 | -46 | 30 | R | Cingulate Gyrus, posterior division |
| 39 | -3.37 | 10 | -50 | -54 | R | Right IX |
| | -3.05 | 10 | -56 | -50 | R | Right IX |
| 36 | -3.33 | 32 | -40 | -46 | R | Right VIIIa |
| | -2.89 | 38 | -40 | -42 | R | Right VIIb |
| | -2.48 | 42 | -42 | -46 | R | Right VIIb |
| 34 | 3.03 | -26 | -10 | 48 | L | Precentral Gyrus |
| | 2.84 | -32 | -6 | 46 | L | Precentral Gyrus |
| 34 | 4.6 | 4 | -2 | 6 | R | Right Thalamus |
| 34 | 4.72 | 20 | -24 | -26 | R | Parahippocampal Gyrus, posterior division |
| 33 | 3.13 | -6 | 4 | 56 | L | Juxtapositional Lobule Cortex (formerly Supplementary Motor Cortex) |
| | 2.33 | -4 | 2 | 48 | L | Juxtapositional Lobule Cortex (formerly Supplementary Motor Cortex) |

| Important Clusters from the PLSDA Model Classifying Romance (scenarios) | | | | | | |
|---|---|---|---|---|---|---|
| 33 | -3.34 | -26 | 8 | -36 | L | Temporal Pole |
| | -2.59 | -22 | 4 | -42 | L | Temporal Pole |
| | -2.55 | -28 | 6 | -44 | L | Temporal Pole |
| 32 | -3.57 | -32 | -34 | -16 | L | Parahippocampal Gyrus, posterior division |
| | -2.91 | -30 | -34 | -24 | L | Temporal Fusiform Cortex, posterior division |
| 32 | -4.26 | 0 | -10 | -2 | C | Left Thalamus |
| | -3.33 | 4 | -16 | -2 | R | Right Thalamus |
| | -3.17 | -6 | -8 | 0 | L | Left Thalamus |
| 31 | -3.07 | -26 | -82 | 36 | L | Lateral Occipital Cortex, superior division |
| | -2.36 | -24 | -86 | 40 | L | Lateral Occipital Cortex, superior division |
| | -2.15 | -30 | -88 | 34 | L | Lateral Occipital Cortex, superior division |
| 31 | -3 | -24 | 36 | -14 | L | Frontal Orbital Cortex |
| | -2.68 | -30 | 36 | -12 | L | Frontal Orbital Cortex |
| 30 | -3.18 | -30 | -10 | -34 | L | Parahippocampal Gyrus, anterior division |
| | -3.14 | -28 | -2 | -34 | L | Parahippocampal Gyrus, anterior division |

*Sadness*

| Important Clusters from the PLSDA Model Classifying Sadness (scenarios) | | | | | | |
|---|---|---|---|---|---|---|
| **Cluster Size (Voxels)** | **Max Coef** | **MNI** | | | **Hemisphere** | **Anatomical Label** |
| | | **x** | **y** | **z** | | |
| 267 | 4.64 | -42 | -56 | 26 | L | Angular Gyrus |
| | 4.55 | -44 | -60 | 34 | L | Lateral Occipital Cortex, superior division |
| | 2.98 | -58 | -62 | 18 | L | Lateral Occipital Cortex, superior division |
| 266 | 3.41 | 52 | 8 | -24 | R | Temporal Pole |
| | 3.31 | 46 | 22 | -30 | R | Temporal Pole |
| | 3.22 | 58 | -2 | -12 | R | Superior Temporal Gyrus, anterior division |
| 236 | 4.64 | 48 | 22 | -10 | R | Frontal Orbital Cortex |
| | 4.48 | 40 | 22 | -20 | R | Frontal Orbital Cortex |
| | 3.68 | 34 | 18 | -24 | R | Temporal Pole |
| 215 | 4.63 | -32 | 14 | -22 | L | Frontal Orbital Cortex |
| | 4.43 | -40 | 4 | -18 | L | Temporal Pole |
| | 4.03 | -40 | 18 | -20 | L | Frontal Orbital Cortex |
| 174 | 3.48 | -2 | -54 | 28 | L | Cingulate Gyrus, posterior division |
| | 3.19 | 0 | -64 | 30 | C | Precuneous Cortex |
| | 2.82 | 0 | -72 | 30 | C | Precuneous Cortex |
| 137 | 3.76 | 54 | -56 | 24 | R | Angular Gyrus |
| | 3.72 | 60 | -56 | 22 | R | Angular Gyrus |
| | 2.97 | 48 | -54 | 22 | R | Angular Gyrus |
| 133 | -5.23 | -52 | -64 | -20 | L | Inferior Temporal Gyrus, temporooccipital part |
| | -3.38 | -48 | -72 | -18 | L | Lateral Occipital Cortex, inferior division |
| | -3.02 | -38 | -66 | -20 | L | Occipital Fusiform Gyrus |
| 114 | -4.77 | -52 | 40 | 14 | L | Frontal Pole |
| | -3.09 | -42 | 34 | 14 | L | Inferior Frontal Gyrus, pars triangularis |
| 102 | -3.15 | 48 | 40 | 12 | R | Frontal Pole |
| | -2.94 | 42 | 38 | 18 | R | Frontal Pole |
| | -2.8 | 42 | 38 | 10 | R | Frontal Pole |
| 87 | 3.58 | -62 | -16 | -6 | L | Superior Temporal Gyrus, posterior division |
| | 3.22 | -58 | -24 | -6 | L | Superior Temporal Gyrus, posterior division |
| | 2.19 | -52 | -26 | -4 | L | Superior Temporal Gyrus, posterior division |
| 84 | -4.77 | 26 | 6 | -20 | R | Temporal Pole / Frontal Orbital Cortex |
| | -4.31 | 14 | 2 | -24 | R | Parahippocampal Gyrus, anterior division |
| | -3.81 | 30 | 4 | -24 | R | Temporal Pole |

| Important Clusters from the PLSDA Model Classifying Sadness (scenarios) | | | | | | |
|---|---|---|---|---|---|---|
| 79 | 3.01 | 0 | 58 | 26 | C | Frontal Pole |
| | 2.51 | 4 | 48 | 16 | R | Paracingulate Gyrus |
| | 2.44 | 0 | 58 | 18 | C | Frontal Pole |
| 61 | -3.15 | -44 | -48 | -20 | L | Inferior Temporal Gyrus, temporooccipital part / Temporal Occipital Fusiform Cortex |
| | -2.54 | -50 | -48 | -20 | L | Inferior Temporal Gyrus, temporooccipital part |
| | -2.51 | -48 | -54 | -24 | L | Inferior Temporal Gyrus, temporooccipital part |
| 59 | -3.67 | 14 | -94 | -18 | R | Occipital Pole |
| | -3.44 | 14 | -96 | -12 | R | Occipital Pole |
| | -2.83 | 8 | -96 | -6 | R | Occipital Pole |
| 58 | 3.55 | -2 | 60 | -6 | L | Frontal Pole |
| | 3.32 | 2 | 64 | -10 | R | Frontal Pole |
| | 2.07 | 6 | 68 | -12 | R | Frontal Pole |
| 50 | -3.46 | -60 | -36 | 46 | L | Supramarginal Gyrus, anterior division |
| | -2.52 | -64 | -24 | 40 | L | Postcentral Gyrus |
| 49 | 3.37 | -54 | 32 | -6 | L | Inferior Frontal Gyrus, pars triangularis |
| | 2.56 | -46 | 30 | -6 | L | Frontal Orbital Cortex |
| | 2.55 | -38 | 24 | -2 | L | Frontal Orbital Cortex |
| 49 | -3.77 | -42 | -6 | 0 | L | Insular Cortex |
| | -3.21 | -40 | -10 | -4 | L | Insular Cortex |
| 46 | 3.03 | 56 | 24 | 8 | R | Inferior Frontal Gyrus, pars triangularis |
| 41 | -3.99 | -58 | 8 | 0 | L | Precentral Gyrus |
| | -2.91 | -64 | -2 | 2 | L | Superior Temporal Gyrus, anterior division |
| 41 | -3.92 | 32 | -86 | -18 | R | Lateral Occipital Cortex, inferior division |
| | -2.99 | 28 | -92 | -18 | R | Occipital Pole |
| | -2.96 | 38 | -82 | -20 | R | Lateral Occipital Cortex, inferior division |
| 40 | -3.03 | 2 | -38 | 50 | R | Precuneous Cortex |
| | -2.54 | 4 | -34 | 46 | R | Cingulate Gyrus, posterior division |
| 36 | 3.02 | 0 | -58 | 42 | C | Precuneous Cortex |
| | 2.02 | 0 | -64 | 40 | C | Precuneous Cortex |
| 35 | 2.75 | -48 | 16 | -34 | L | Temporal Pole |
| | 2.61 | -42 | 16 | -34 | L | Temporal Pole |
| 34 | 2.9 | 4 | 52 | 44 | R | Superior Frontal Gyrus |
| | 2.34 | 2 | 46 | 34 | R | Superior Frontal Gyrus |
| 33 | 3.42 | 54 | -16 | 14 | R | Central Opercular Cortex |

| Important Clusters from the PLSDA Model Classifying Sadness (scenarios) | | | | | | |
|---|---|---|---|---|---|---|
| | 2.88 | 60 | -12 | 8 | R | Central Opercular Cortex |
| 32 | -5.68 | -12 | -28 | -18 | L | Brain-Stem |
| | -3.83 | -14 | -34 | -18 | L | Left I-IV |
| 32 | -4.47 | 42 | 8 | -20 | R | Temporal Pole |
| | -2.24 | 38 | 16 | -20 | R | Temporal Pole |
| 31 | 2.66 | 0 | 44 | 44 | C | Superior Frontal Gyrus |
| | 2.3 | 2 | 42 | 50 | R | Superior Frontal Gyrus |
| 30 | -4.12 | -26 | 4 | -26 | L | Temporal Pole |
| | -2.36 | -30 | 2 | -22 | L | Temporal Pole |
| | 4.55 | -44 | -60 | 34 | L | Lateral Occipital Cortex, superior division |
| | 2.98 | -58 | -62 | 18 | L | Lateral Occipital Cortex, superior division |

*Surprise*

| Important Clusters from the PLSDA Model Classifying Surprise (scenarios) | | | | | | |
|---|---|---|---|---|---|---|
| Cluster Size (Voxels) | Max Coef | MNI | | | Hemisphere | Anatomical Label |
| | | x | y | z | | |
| 307 | -5.27 | -18 | 2 | -22 | L | Parahippocampal Gyrus, anterior division |
| | -4.86 | -26 | 2 | -18 | L | Parahippocampal Gyrus, anterior division |
| | -4.47 | -40 | 6 | -16 | L | Temporal Pole |
| 202 | -4.53 | 2 | 4 | -12 | R | Right Cerebral Cortex |
| | -4.46 | -2 | 14 | -16 | L | Subcallosal Cortex |
| | -3.85 | 0 | 12 | -8 | C | Subcallosal Cortex |
| 201 | 4.2 | 46 | -52 | 58 | R | Angular Gyrus |
| | 3.58 | 48 | -58 | 52 | R | Lateral Occipital Cortex, superior division |
| | 3.31 | 56 | -54 | 46 | R | Angular Gyrus |
| 170 | -4.04 | -4 | -52 | 20 | L | Cingulate Gyrus, posterior division |
| | -3.54 | -2 | -58 | 20 | L | Precuneous Cortex |
| | -3.17 | 6 | -52 | 26 | R | Cingulate Gyrus, posterior division |
| 149 | 3.69 | 26 | 12 | 56 | R | Superior Frontal Gyrus |
| | 3.48 | 28 | 16 | 60 | R | Superior Frontal Gyrus |
| | 2.88 | 30 | 8 | 66 | R | Superior Frontal Gyrus |
| 141 | 3.8 | 42 | 56 | -4 | R | Frontal Pole |
| | 3.27 | 46 | 52 | -8 | R | Frontal Pole |
| | 3.19 | 32 | 62 | 0 | R | Frontal Pole |
| 97 | -3.37 | 0 | 44 | -12 | C | Frontal Medial Cortex |
| | -3.3 | 2 | 52 | -6 | R | Paracingulate Gyrus |
| | -2.13 | -4 | 52 | 2 | L | Paracingulate Gyrus |
| 88 | -3.19 | 4 | 8 | 46 | R | Paracingulate Gyrus |
| | -3.17 | 0 | 4 | 42 | C | Cingulate Gyrus, anterior division |
| | -2.44 | 4 | 4 | 52 | R | Juxtapositional Lobule Cortex (formerly Supplementary Motor Cortex) |
| 81 | 3.23 | 64 | -50 | -8 | R | Middle Temporal Gyrus, temporooccipital part |
| | 2.74 | 58 | -50 | -6 | R | Middle Temporal Gyrus, temporooccipital part |
| | 2.74 | 64 | -46 | -14 | R | Inferior Temporal Gyrus, temporooccipital part |
| 78 | 3.5 | 68 | -32 | -4 | R | Middle Temporal Gyrus, posterior division |
| | 2.67 | 66 | -36 | 0 | R | Middle Temporal Gyrus, posterior division |
| | 2.52 | 60 | -24 | -4 | R | Superior Temporal Gyrus, posterior division |
| 68 | -2.71 | -2 | -6 | 58 | L | Juxtapositional Lobule Cortex (formerly Supplementary Motor Cortex) |

| Important Clusters from the PLSDA Model Classifying Surprise (scenarios) | | | | | | |
|---|---|---|---|---|---|---|
| | -2.6 | 2 | -14 | 60 | R | Precentral Gyrus |
| | -2.57 | 0 | -12 | 52 | C | Juxtapositional Lobule Cortex (formerly Supplementary Motor Cortex) |
| 62 | 3.01 | -32 | -42 | -42 | L | Left VIIb |
| | 2.81 | -34 | -46 | -56 | L | Left VIIIa |
| | 2.69 | -26 | -42 | -52 | L | Left VIIIb |
| 60 | 3.32 | -16 | -80 | -22 | L | Left Crus I |
| | 2.75 | -8 | -80 | -26 | L | Left Crus I |
| | 2.46 | -10 | -78 | -32 | L | Left Crus II |
| 58 | -2.97 | 12 | -74 | 38 | R | Precuneous Cortex |
| | -2.72 | 14 | -70 | 34 | R | Precuneous Cortex |
| 57 | 3.9 | 6 | 30 | 64 | R | Superior Frontal Gyrus |
| | 3.15 | 6 | 36 | 60 | R | Superior Frontal Gyrus |
| | 2.63 | 2 | 34 | 56 | R | Superior Frontal Gyrus |
| 57 | -2.91 | 0 | -76 | 34 | C | Precuneous Cortex |
| | -2.62 | -8 | -78 | 38 | L | Precuneous Cortex |
| | -2.61 | 0 | -70 | 28 | C | Precuneous Cortex |
| 56 | -4.26 | 2 | 70 | 4 | R | Frontal Pole |
| | -2.66 | 0 | 64 | 2 | C | Frontal Pole |
| 49 | 3.46 | 10 | -2 | 10 | R | Right Thalamus |
| | 3.06 | 14 | -2 | 18 | R | Right Caudate |
| 45 | 3 | -58 | -46 | -14 | L | Inferior Temporal Gyrus, temporooccipital part |
| | 2.79 | -58 | -50 | -8 | L | Middle Temporal Gyrus, temporooccipital part |
| 44 | 5.01 | -2 | -42 | -38 | L | Brain-Stem |
| | 3.67 | 0 | -42 | -44 | C | Brain-Stem |
| | 2.49 | 4 | -38 | -36 | R | Brain-Stem |
| 43 | -2.73 | 36 | -94 | -2 | R | Occipital Pole |
| | -2.38 | 32 | -96 | 2 | R | Occipital Pole |
| | -2.26 | 40 | -90 | -6 | R | Lateral Occipital Cortex, inferior division |
| 41 | 3.2 | 28 | -72 | -26 | R | Right Crus I |
| | 2.24 | 26 | -78 | -24 | R | Right Crus I |
| | 2.17 | 34 | -72 | -22 | R | Right Crus I |
| 40 | 8.84 | -4 | -42 | -70 | L | Brain-Stem |
| | 5.22 | 6 | -44 | -70 | R | Brain-Stem |
| | 2.42 | -4 | -38 | -64 | L | Brain-Stem |

| Important Clusters from the PLSDA Model Classifying Surprise (scenarios) | | | | | | |
|---|---|---|---|---|---|---|
| 39 | -4.07 | -4 | 30 | -20 | L | Subcallosal Cortex |
| | -2.37 | 0 | 24 | -26 | C | Subcallosal Cortex |
| | -2.23 | -2 | 34 | -14 | L | Paracingulate Gyrus |
| 37 | -4.03 | 8 | -50 | 2 | R | Lingual Gyrus |
| 37 | 2.63 | -30 | -74 | -24 | L | Left Crus I |
| | 2.55 | -24 | -70 | -30 | L | Left Crus I |
| | 2.31 | -20 | -66 | -34 | L | Left VI |
| 35 | 2.9 | 40 | -46 | -40 | R | Right Crus I |
| | 2.38 | 46 | -44 | -40 | R | Right Crus I |
| 35 | -2.77 | -2 | -38 | 50 | L | Precuneous Cortex |
| | -2.62 | -8 | -32 | 46 | L | Precentral Gyrus |
| 34 | 2.83 | -26 | -72 | -38 | L | Left Crus II |
| 34 | -3.45 | 2 | 24 | 28 | R | Cingulate Gyrus, anterior division |
| 34 | -6.77 | -16 | -30 | -20 | L | Parahippocampal Gyrus, posterior division |
| | -3.66 | -10 | -36 | -18 | L | Left I-IV |
| | -3.41 | -20 | -26 | -26 | L | Parahippocampal Gyrus, posterior division |
| 34 | 3.91 | -20 | -54 | -12 | L | Lingual Gyrus |
| 32 | 2.59 | -44 | 52 | -6 | L | Frontal Pole |
| | 2.57 | -36 | 56 | -8 | L | Frontal Pole |
| | 2.2 | -46 | 46 | -2 | L | Frontal Pole |
| 32 | -4.4 | -44 | -10 | 0 | L | Insular Cortex |
| 32 | -4.26 | -8 | -94 | -18 | L | Occipital Pole |
| 31 | 3.8 | -4 | -26 | -12 | L | Brain-Stem |
| | 2.72 | -6 | -32 | -8 | L | Brain-Stem |
| 30 | -3.82 | 0 | 36 | 6 | C | Cingulate Gyrus, anterior division |
| | -2.78 | 2 | 34 | 14 | R | Cingulate Gyrus, anterior division |
